# Long-range single-molecule mapping of chromatin accessibility in eukaryotes

**DOI:** 10.1101/504662

**Authors:** Zohar Shipony, Georgi K. Marinov, Matthew P. Swaffer, Nasa A. Sinott-Armstrong, Jan M. Skotheim, Anshul Kundaje, William J. Greenleaf

## Abstract

Active regulatory elements in eukaryotes are typically characterized by an open, nucleosome-depleted chromatin structure; mapping areas of open chromatin has accordingly emerged as a widely used tool in the arsenal of modern functional genomics. However, existing approaches for profiling chromatin accessibility are limited by their reliance on DNA fragmentation and short read sequencing, which leaves them unable to provide information about the state of chromatin on larger scales or reveal coordination between the chromatin state of individual distal regulatory elements. To address these limitations, we have developed a method for profiling accessibility of individual chromatin fibers at multi-kilobase length scale (SMAC-seq, or <u>S</u>ingle-<u>M</u>olecule long-read <u>A</u>ccessible <u>C</u>hromatin mapping <u>seq</u>uencing assay), enabling the simultaneous, high-resolution, single-molecule assessment of the chromatin state of distal genomic elements. Our strategy is based on combining the preferential methylation of open chromatin regions by DNA methyltransferases (CpG and GpC 5-methylcytosine (5mC) and N^6^-methyladenosine (m^6^A) enzymes) and the ability of long-read single-molecule nanopore sequencing to directly read out the methylation state of individual DNA bases. Applying SMAC-seq to the budding yeast *Saccharomyces cerevisiae*, we demonstrate that aggregate SMAC-seq signals match bulk-level accessibility measurements, observe single-molecule protection footprints of nucleosomes and transcription factors, and quantify the correlation between the chromatin states of distal genomic elements.

---

The packaging of DNA by nucleosomes into chromatin is a major organizing principle of genome organization in eukaryotes. The majority of the genome is tightly packaged by nucleosomal particles that wrap around DNA (usually ~147bp), thus making it inaccessible to binding by most regulatory proteins. Conversely, regions of open chromatin tend to be strongly associated with regulatory elements (REs), such as enhancers, promoters, and insulators, and nucleosomes often exhibit characteristic occupancy patterns in their vicinity. These biological properties have proven highly useful for identifying candidate such elements (cREs), and in turn for understanding the functional organization of genomes. Regions of open chromatin have greatly increased sensitivity to cleavage by nucleases such as DNAse I, as already noted nearly four decades ago for promoter and enhancer elements around individual genes ^1–3^. Subsequent advances in microarray ^4,5^ and DNA sequencing technologies^6,7^ have enabled DNAse hypersensitivity-based mapping of cREs genome-wide. Similarly, digestion of DNA is inhibited by nucleosome occupancy, and MNase digestion of chromatin (MNase-seq) has accordingly become a widely used tool to map nucleosome positioning throughout genomes ^9^. More recently, the Tn5 transposase has also been used as a facile probe of chromatin accessibility ^8^. However, while short read-based assays provide immensely useful information about the identity of cREs and positioned nucleosomes, they give little insight into the long-range physical organization of individual chromatin fibers. Because cleavage-based approaches remove the linkage between distal segments of DNA molecules, classical functional genomic assays return estimates of the relative chromatin states only at short localized regions of the genome. To what extent these states are coordinated across distance and what the distribution of chromatin states looks like at the kilo– to multikilobase scale is largely unknown. Well established tools exist for mapping direct interactions between distal genomic elements in the context of three-dimensional genome architecture^10–12^ but they only capture pairwise interactions within large cell populations, not the chromatin state of the interacting regions, and they are often of limited resolution. Similar limitations apply to emerging methods for studying local chromatin structure^13^. Super-resolution microscopy using highly multiplexed fluorescent probes is a powerful tool for revealing the folding of the chromatin fiber at the single cell level ^14^, but this approach is limited to a small number of loci and does not provide high-resolution base-pair level information about regulatory states. Cryoelectron microscopy-based approaches ^15^ allow direct observations of the nucleosomal state along individual chromatin fibers, but at present the underlying DNA sequence cannot be linked to these measurements. More recently, long-read sequencing was used to map MNase cleavage events at multinucleosomal lengths^21^; however, such approaches only provide information about two points in genomic space leaving the state of chromatin in the intervening sequence unknown.

These technological limitations prevent the investigation of the manner in which chromatin states of adjacent elements are correlated and how such coordination might play a role in gene regulation (for example, by creating self-reinforcing or mutually exclusive epigenetic states). To address these technological limitations, we have developed SMAC-seq, a versatile, single molecule method that directly assays long-range nucleosome positioning and accessibility states within the chromatin fiber. We use this method to study chromatin architecture and co-accessibility states in the yeast *Saccharomyces cerevisiae*, both under normal growth and conditions of cellular stress. We use SMAC-seq to assess the degree of coordination between the positions of nearby nucleosome particles, enumerate mutually exclusive regulatory states along individual loci, and observe coordinated changes in nucleosome positioning and chromatin accessibility upon transcriptional activation. SMAC-seq also enables high-resolution footprinting of transcription factor (TF) occupancy, and provides strand-specific information about the exposure of DNA bound to nucleosomes and other proteins. We expect future applications of, improvements on, and extensions of the SMAC-seq approach to enable novel insights into the dynamics of chromatin states in the context of a wide variety of experimental systems and biological questions.

## Results

### SMAC-seq maps chromatin accessibility and nucleosome positioning at the multi-kb scale

SMAC-seq is built on the conceptual foundations of the NOMe-seq assay ^16,17^. In its original form, NOMe-seq relies on the preferential modification of bases within accessible DNA with M.CviPI (a GpC-specific 5mC methyltransferase), followed by bisulfite conversion and Illumina-based sequencing readout. A more recent variation, dSMF ^18^ (**d**ual-enzyme **S**ingle **M**olecule **F**ootprinting), utilized an additional CpG-specific 5mC methyltransferase (M.SssI) to map promoter states in *Drosophila* cells in finer detail. Such an approach is applicable in the *Drosophila* context thanks to the absence of endogenous CpG methylation in flies.

We build on the dSMF approach by adding a third DNA-modifying enzyme, the m^6^A methyltransferase EcoGII, and adapting the protocol to nanopore sequencing (Figure 1A). Briefly, isolated nuclei are treated with the three enzymes, which preferentially modify DNA within accessible chromatin. High-molecular weight (HMW) DNA is then isolated and subjected to long-read single-molecule sequencing using the Oxford Nanopore platform. Using the ability of nanopore sequencing to directly read modified DNA bases^19,20^, we then obtain methylation maps for individual DNA molecules on a multikilobase scale (see the Methods section for details), which we then interpret in terms of chromatin accessibility.

**Figure 1:**
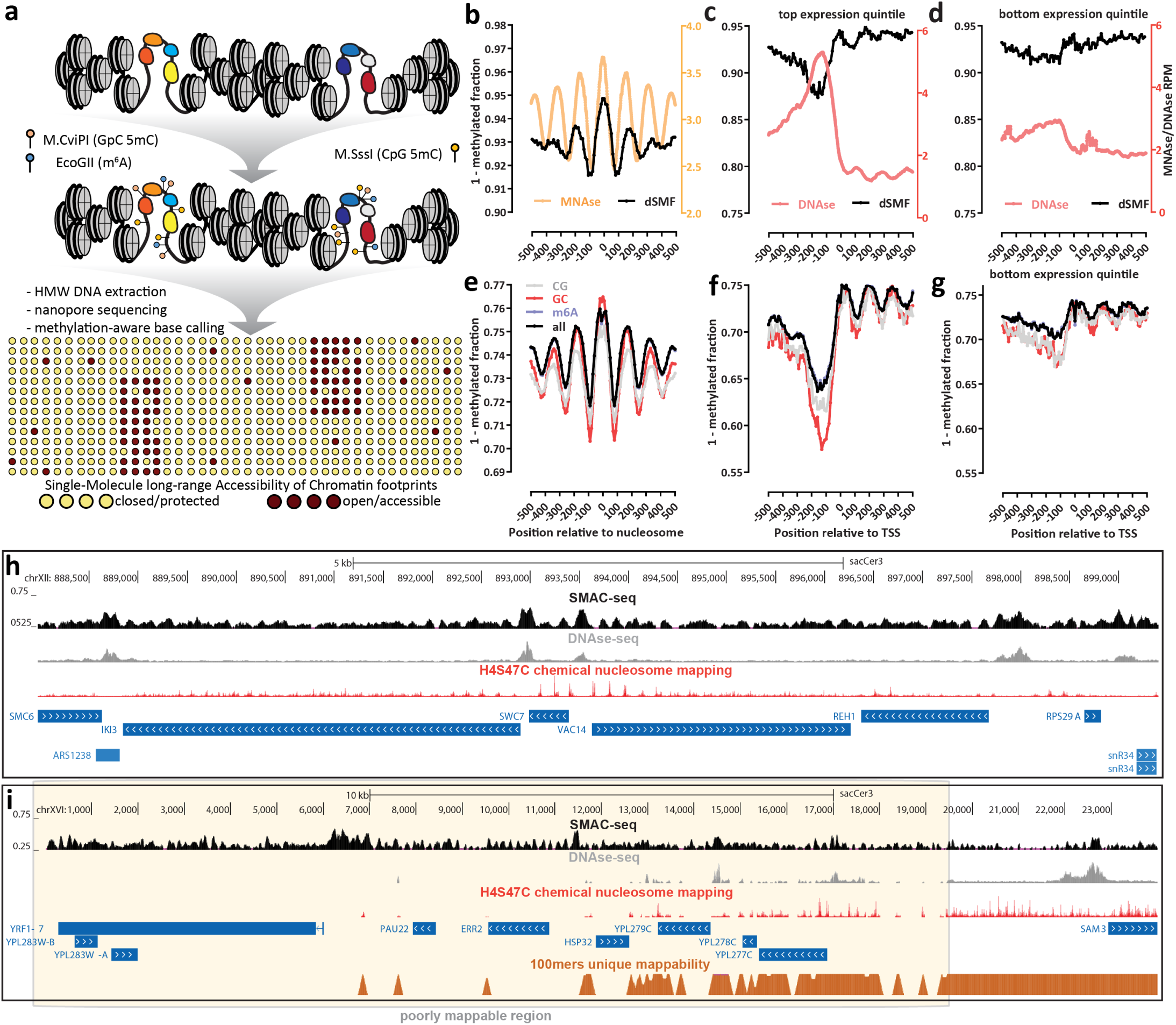
The SMAC-seq assay for profiling chromatin accessibility and nucleosome positioning at the multikilobase scale. (a) Outline of the SMAC-seq assay. Intact chromatin is treated with m^6^A and CpG and GpC 5mC methyltransferases, which preferentially methylate DNA bases in open chromatin regions. HMW DNA is then isolated, subjected to nanopore sequencing and methylated bases are used to reconstruct the open chromatin state within individual molecules. (b-h) SMAC-seq faithfully captures chromatin accessibility around promoters and positioned nucleosomes in *S. cerevisiae*; (b) MNAse-seq and dSMF profiles around chemically mapped positioned nucleosome dyads; (c) DNAse-seq and dSMF profiles around the top 20% highly expressed genes in *S. cerevisiae*; (d) DNAse-seq and dSMF profiles around the bottom 20% expressed genes in *S. cerevisiae*; (e) SMAC-seq profile around chemically mapped positioned nucleosomes dyads (shown is the “diamide 0 min rep2” sample); (f) SMAC-seq profile around the top 20% highly expressed genes in *S. cerevisiae*; (g) SMAC-seq profile around the bottom 20% expressed genes in *S. cerevisiae*; (h) SMAC-seq correlates closely with both DNAse-seq and nucleosome occupancy profiling at the level of individual loci, and provides a combined readout of accessibility and nucleosome positioning. Shown is aggregate SMAC-seq signal along the genome (aggregated over 50-bp windows sliding every 5 bp; see Methods for details) together with DNAse-seq, nucleosome chemical mapping data, and transcriptional activity (measured by PRO-seq and PRO-cap). Large aggregate SMAC-seq signal enrichments match closely with DNAse accessibility peaks, while smaller aggregate SMAC-seq peaks are inversely correlated with positioned nucleosomes; (i) SMAC-seq profiles chromatin accessibility in repetitive regions of the genomes that are “invisible” to short reads. Shown is the telomeric region of chrXVI

The addition of m^6^A methylation is of key importance to making SMAC-seq a high-resolution broadly applicable assay. Many eukaryote genomes are endogenously methylated at 5mC positions^22,23^. This is usually in a CpG context, in particular in metazoans, but it is not always limited to CpG dinucleotides. For example, in plants methylation in CHG and CHH contexts is also a common occurrence^24^, thus even results obtained with M.CviPI alone are confounded by endogenous methylation. In addition, CpG and GpC dinucleotides are relatively rare in the genome; the average resolution achieved by the combination of CpG and GpC methyltransferases is *>*10 bp in *Drosophila* and ~15 bp in yeast. The addition of m^6^A dramatically increases the resolution of SMAC-seq, to ~3 bp in all main model organisms (Supplementary Figures 1, 4, and 5).

We initially developed and optimized the method in the budding yeast *Saccharomyces cerevisiae* under normal growth conditions, as *Saccharomyces cerevisiae* has no endogenous DNA methylation (allowing the simultaneous use of all three enzymes) and has a small genome (~12 Mbp), making it possible to achieve very high depth of nanopore sequencing coverage. To verify the specificity and efficiency of the enzymatic treatments, we carried out both dSMF experiments (i.e M.CviPI + M.SssI treatment) on yeast chromatin and M.CviPI + M.SssI + EcoGII reactions on naked yeast DNA (as well as untreated naked DNA controls), and subjected the resulting material to whole-genome bisulfite sequencing (see the Methods section for more details). We observe ≥95% methylation in both the CpG and GpC contexts when starting with naked DNA, ≤10% when working with chromatin, and nearly 0% on naked DNA (Supplementary Figure 3). Comparing dSMF profiles to DNase-seq and MNase-seq profiles around transcription start sites (TSSs) and positioned nucleosomes (obtained from previous H4S47C chemical mapping studies ^25^) revealed the expected nucleosome depletion and nucleosomal patterns, respectively (Figure 1b-d).

We then carried out a SMAC-seq experiment on unsynchronized *S. cerevisiae* cells using all three enzymes (experimental details described in the Methods section). We isolated HMW DNA and performed nanopore sequencing on the MinION platform. To call methylated bases, we first used Albacore for raw base calling, and then applied the Tombo ^26^ algorithm in “de novo” mode (running on top of the Minimap aligner ^27^) for “resquiggling” of raw nanopore signal to the genomic sequence, and calling of methylated bases. After mapping, we obtained reads with a median length of ~1.5 kbp from this initial experiment (“Sample 1”; Supplementary Table 1), which allows the capture of multiple promoter regions per fragment for much of the yeast genome (Supplementary Figure 2). We also analyzed our initial dataset with Nanopolish ^19^, an alternative algorithm for calling methylated bases, which is capable of identifying 5mC events in CpG and GpC context.

Unlike bisulfite-based conversion followed by sequencing-by-synthesis, nanopore-based direct measurement of nucleotide modifications does not provide unambiguous binary calls for methylated bases. Instead, methylation probabilities are obtained for each base. We therefore examined multiple strategies for binarizing these methylation calls within each read. We observed that while per-base methylation probabilities are skewed towards the two ends of the [0, 1] interval, a substantial number of bases have probabilities in between those extremes (Supplementary Figure 6). We find that binarization at *p* = 0.5 delivers the most optimal results (Supplementary Figures 7 and 8) in terms of signal-to-noise ratio. Using a simple average of these binarized per-base methylation values, we compared SMAC-seq to dSMF, MNase-seq, and DNase-seq profiles, as well as signal from ChIP-seq for RNA Polymerase (Pol2) and transcription initiation factors around known chromatin features such as active promoters and well positioned nucleosomes (Figure 1e-g, Supplementary Figures 13 and 16). SMAC-seq faithfully reproduces nucleosomal positioning throughout the genome and nucleosome depletion around promoters, and, strikingly, has a larger observed dynamic range than dSMF data (possibly due to the higher mapping efficiency of long nanopore reads). Using comparisons with dSMF data we estimate the false positive rate of methylation base calling to be on the order of 20-25% for Tombo and around 10-15% for Nanopolish (Supplementary Figure 12). We also examined potential sequence biases inherent to the combination of methylation enzymes and base calling algorithm. We find modest differences in methylation levels for different *k*-mers in the genome (less than two-fold for *k* = 6; Supplementary Figures 10 and 11).

In practice, the biologically relevant length scale of accessibility measurements is usually larger than an individual base. Furthermore, given noise intrinsic to single molecule sequencing methods, we reasoned that sharing methylation information between adjacent bases should improve the reliability of overall accessibility measurements. We therefore developed a simple Bayesian procedure to aggregate the methylation probabilities at individual bases and derive accessibility calls at the level of single reads over windows of arbitrary size (described in detail in the Methods section and thereafter referred to as “aggregate” signal as opposed to “average” signal, which refers to simple probability averaging and binarization).

We also note that sometimes we observe a subpopulation of reads that appear to be either entirely fully methylated or fully methylated over large segments of their length (Supplementary Figure 9 and Figure 2a). We interpret these as originating from naked DNA molecules most likely deriving from dead cells. As such reads can confound many analyses, in particular when measuring coaccessibility within single reads, we devised a procedure for filtering them out (the resulting sets of reads are referred to as “filtered reads”; (Supplementary Figures 19 and 20). However, as discussed below, in certain situations chromatin is indeed largely nucleosome-free over specific regions in vivo, and in such cases filtering out fully methylated reads removes real biological signal. For these unique special case loci, we do not eliminate reads based on a very high fraction of accessibility (see Methods).

**Figure 2:**
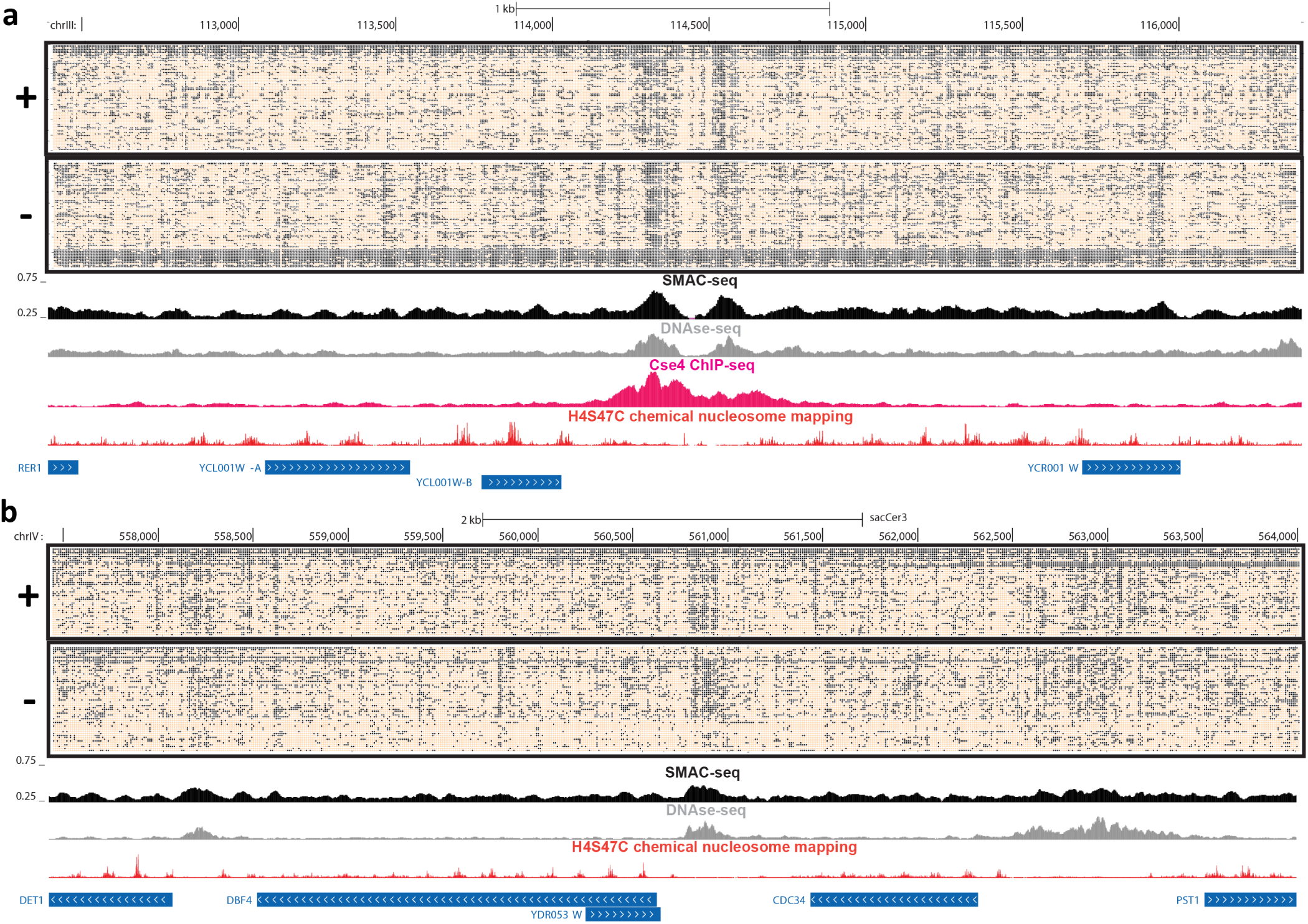
SMAC-seq provides a single-molecule linked-read view of the chromatin landscape. (a) Unfiltered nanopore reads fully spanning the 4-kilobase neighborhood of the centromere of *S. cerevisiae* chrIII (“aggregate” signal “Sample 1”). (b) Unfiltered nanopore reads fully spanning a 6.6-kilobase neighborhood encompassing several genes on chrIV (“aggregate” signal from “Sample 1”). In both cases, accessibility is shown at 10-bp resolution (see Methods section for details) for the single-molecule display, and aggregated over sliding (every 5 bases) 50-bp windows for the average SMAC-seq track.

We then compared average SMAC-seq profiles against chemical maps of positioned nucleosomes (generated using H4S47C substitutions and copper-induced cleavage ^25^), DNase-seq, and maps of transcriptional activity at the level of individual loci in the genome (Figure 1h). Qualitatively, we observe that large peaks in SMAC-seq signal profiles match very closely with DNase-seq peaks, while smaller “bumps” in the SMAC-seq signal profile are inversely correlate with positioned nucleosomes, consistent with labeling of linker DNA. We also observe positive correlation between average SMAC-seq methylation levels and DNase-seq and ATAC-seq coverage over promoter regions (Supplementary Figure 15). Thus SMAC-seq simultaneously probes both regions of “open” chromatin, as well as the position of nucleosomes.

The long reads of nanopore sequencing allow SMAC-seq to provide accessibility maps for the whole yeast genome, not just for the portions of it that are uniquely mappable with short reads (Supplementary Figure 18). For example, SMAC-seq maps chromatin and nucleosomes in the otherwise not uniquely mappable telomere of chrXVI (Figure 1i), which is revealed to contain several active promoters and numerous well positioned nucleosomes. We also used SMAC-seq to characterize chromatin accessibility around multiple transposable elements (Supplementary Figures 17), for several of which we observe open chromatin peaks around their promoters.

### SMAC-seq provides single-molecule accessibility profiles on individual chromatin fibers

Unlike short-read methods for probing chromatin, SMAC-seq allows the profiling of nucleosome positioning and open chromatin within individual long molecules. To demonstrate this capacity, we investigated all reads spanning the 4-kb neighborhood around the centromere of chrIII (Figure 2a). Centromeres in *S. cerevisiae* are specified by a precisely defined sequence element, are occupied by a single strongly positioned nucleosome containing the H3 histone variant Cse4, and are also associated with the CBF3 complex, an essential component of the kinetochore^28^. Yeast centromeric nucleosomes are thought to be nearly perfectly positioned ^28,29^ and thus represent a good case system to study nucleosome positioning at the single-molecule level. We indeed observe strong nucleosomal positioning using SMAC-seq, with nearly all individual reads exhibiting the expected from the presence of a strongly positioned centromere nucleosomal pattern. We also find hints of substructure within the centromeric nucleosome in the form of accessibility traces inside the protected centromeric region and potential protection footprints in its immediate open chromatin vicinity. We find similarly strong positioning for most other centromeric nucleosomes (Supplementary Figures 21-28), but not all (chrX, chrXII and chrXIII appear to be exceptions to this general strong footprinting pattern).

We also illustrate the ability of SMAC-seq to capture accessibility at long-range multikilobase scales with an example from a more generic genomic neighborhood in Figure 2b). This ~6.6-kb span of chrIX contains five genes and three open chromatin regions, one of them fairly large and diffuse. In contrast to the more localized accessibility observed elsewhere, this region exhibits considerable heterogeneity in its accessibility suggesting a complex landscape of protein occupancy.

We next asked if SMAC-seq could reveal binary states of chromatin accessibility. To approach this question we investigated SMAC-seq profiles at ribosomal DNA (rDNA). In yeast, rDNA is organized into multicopy arrays, each unit of which contains a copy of the 35S precursor pre-rRNA, transcribed by Pol I and later processed into mature 18S, 5.8S and 25S rRNAs, as well as a copy of the 5S RNA gene, transcribed by Pol III, and an origin of replication (ARS element in yeast) located in non-transcribed (NTS) regions of the array. Each array unit is ~9.1 kb in length, and each cell’s genome has an array of ~150 copies of this unit^30^. However, this number can vary from cell to cell, and the widely used sacCer3 *S. cerevisiae* genome assembly only contains a single locus with two array copies. Chromatin structure at the rDNA locus has long been known to adopt two distinct conformations ^31–33^, depending on whether or not an individual unit is being transcribed. The active state is thought to be largely devoid of nucleosomes due to the extremely high levels of active transcription^32^; the high-mobility group protein Hmo1 is proposed to replace nucleosomes ^30,34^. However, other studies have alternatively suggested that nucleosomes are found over actively transcribed rDNA arrays ^35^. Of note, rDNA indeed appears to be extremely accessible in short-read assays – around half of reads in a typical ATAC-seq dataset in *S. cerevisiae* are not uniquely mappable even though only a small fraction of the yeast genome consists of repetitive elements; these reads originate primarily from rDNA arrays (Supplementary Figure 29a-e). The same phenomenon is also observed in other yeast species, such as *Schizosaccharomyces pombe* (Supplementary Figure 29f). Active and inactive rDNA arrays are usually estimated to exist in roughly equal proportions in untreated, normally growing cells ^35^. However, methods to precisely observe these alternative states at the population level and relate them to sequence in fine detail have not been available.

SMAC-seq reveals a striking picture of the two alternative mutually exclusive rDNA states at the single molecule level (Figure 3a). About a quarter of full-length molecules exhibit near-full accessibility in the region spanning the 35S transcript, but not in the non-transcribed sequence (NTS) between the 35S and the 5S gene. The rest of the molecules show a typical nucleosomal state with several clearly accessible regions. We observe a broadly similar picture in all of our high-quality SMAC-seq samples (Supplementary Figures 30–33). We note that while this picture is in contrast with the usually reported ~50% of arrays being fully accessible, it is possible that the fully accessible and nucleosomal states fragment differentially during DNA isolation; the fully accessible fraction appears to be in the 50% range when a shorter window around the 35S promoter is examined (Figure 3c). We also observe a localized accessible region just upstream of the 35S transcriptional unit that is present in the nucleosomal subpopulation but is not in an open state in the fully accessible population (in addition to a nearby open chromatin region present in all molecules), suggesting the possibility of a regulatory switch associated with that element. Finally, we also observe at least two (previously unreported) accessible regions located within the 35S transcriptional unit that exhibit strong accessibility in the nucleosome-protected fraction (Figure 3a).

**Figure 3:**
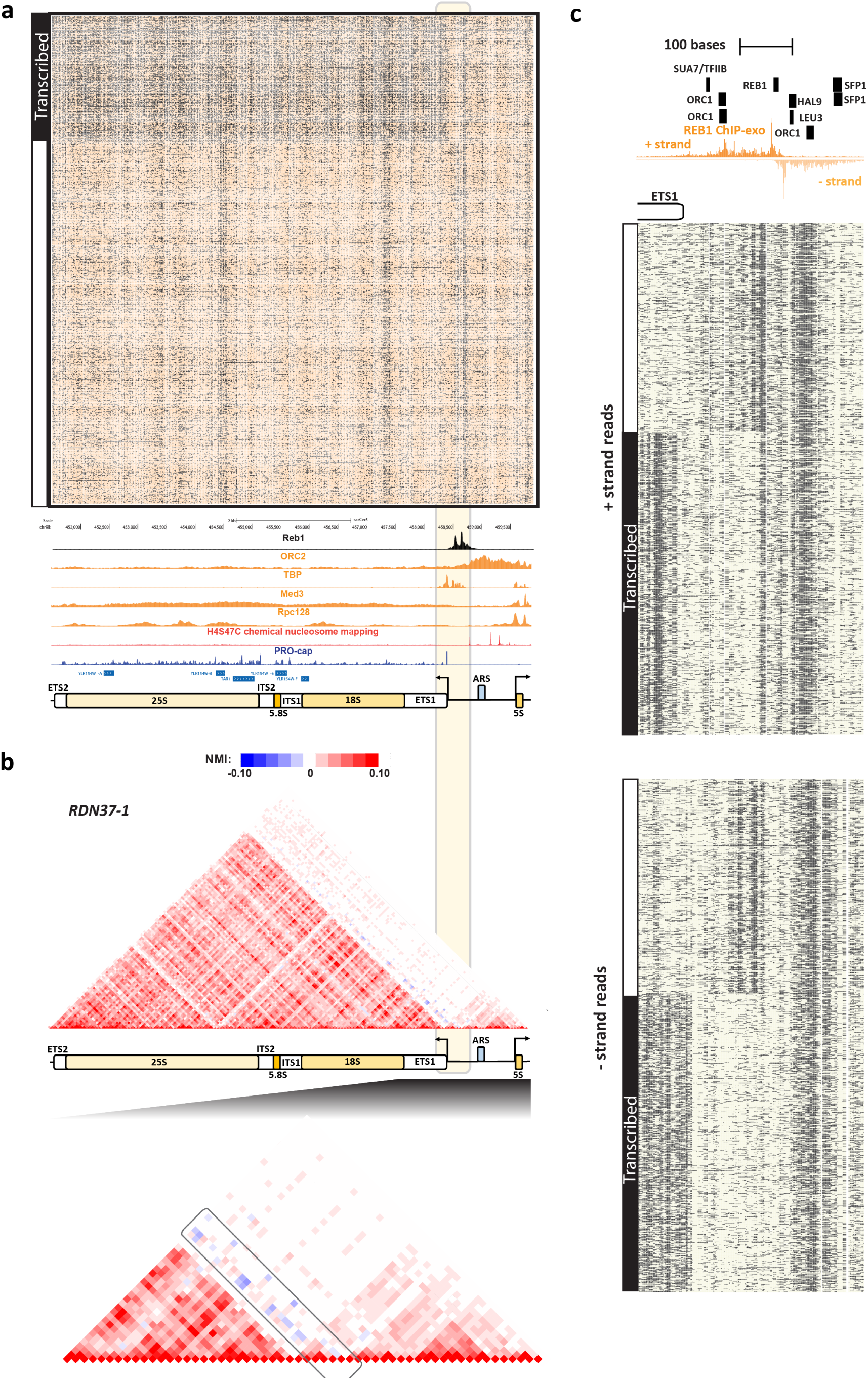
SMAC-seq’s single-molecule readout provides insights into the distribution and relationship between mutually exclusive chromatin yeast rDNA states. (a) SMAC-seq reveals the distribution of alternative chromatin states of rDNA arrays. Shown are all reads covering the *RDN37-1* array in the *RDN1* locus in the “diamide 30 min rep1” experiment (unfiltered reads, “aggregate” signal). See Supplementary Figures 30–33 for additional details. ChIP-seq and ChIP-exo tracks were generated by including and normalizing all multimappers rather than the usual unique-only policy (See the Methods section for more details). (b) Normalized mutual information profiles for the *RDN37-1* array show anti-correlation between the accessibility peaks immediately upstream of the *35S* TSS and the nucleosome-free state over the *35S* transcriptional unit. (c) High-resolution SMAC-seq profiles reveal regulatory protein footprints in the immediate vicinity of the 35S TSS and the Reb1 binding site in the rDNA NTS region (shown are 3000 randomly sampled reads using 10-bp aggregate SMAC-seq signal at 1-bp resolution).

To quantify the extent of (anti-)correlation between chromatin states, we developed a modified normalized mutual information (NMI) metric for assessing the degree of correlation between segments of the genome (see the Methods section for further details). NMI analysis of the rDNA arrays confirmed our observations of the inverse correlation between the 35S open-chromatin state and the accessibility of the upstream element (Figure 3b).

What factors might be behind the observed chromatin state switch? Silencing of rDNA in yeast is thought to be mediated by a Sir2-containing complex called RENT^36^, and a Reb1 binding site in NTS1 has been suggested to recruit corepressors to rDNA repeats^35^. We took a higher-resolution view of NTS1 by integrating SMAC-seq data with available ChIP-exo data for Reb1 and transcription factor motif maps in the region. We find a clear pattern of protection from methylation around the Reb1 motif, which is concordant with ChIP-exo data (Figure 3c), and we also observe signal consistent with footprinting from several other TF motifs. However, the anti-correlated accessibility profile seems to not be exclusively associated with Reb1 binding but rather with the region closer to the 35S TSS. It is likely that other proteins are responsible for establishing this state, but no currently annotated transcription factor recognition motifs are found in the underlying sequence.

### SMAC-seq provides a high-resolution strand-specific view of protein occupancy on DNA

We next asked if SMAC-seq can generally identify transcription factor footprints. We anticipated that the use of m^6^A methylation ought to provide sufficient resolution to footprint most yeast TFs, which is confirmed by analysis of their consensus recognition motifs (Supplementary Figure 34). Averaging genome-wide SMAC-seq profiles over occupied motifs indeed revealed strong protection footprints for several factors, such as Reb1, Rap1 and ORC1 (Origin of Replication Complex) as shown in Figure 4b-c and Supplementary Figure 35. Examination of individual sites confirmed these observations (Supplementary Figures 36– 44). We note that we did not observe strong footprinting for all TFs (e.g. Abf1 and Cbf1; Supplementary Figure 35), suggesting that TF footprinting is probably dependent on the biophysical properties of individual factors.

**Figure 4:**
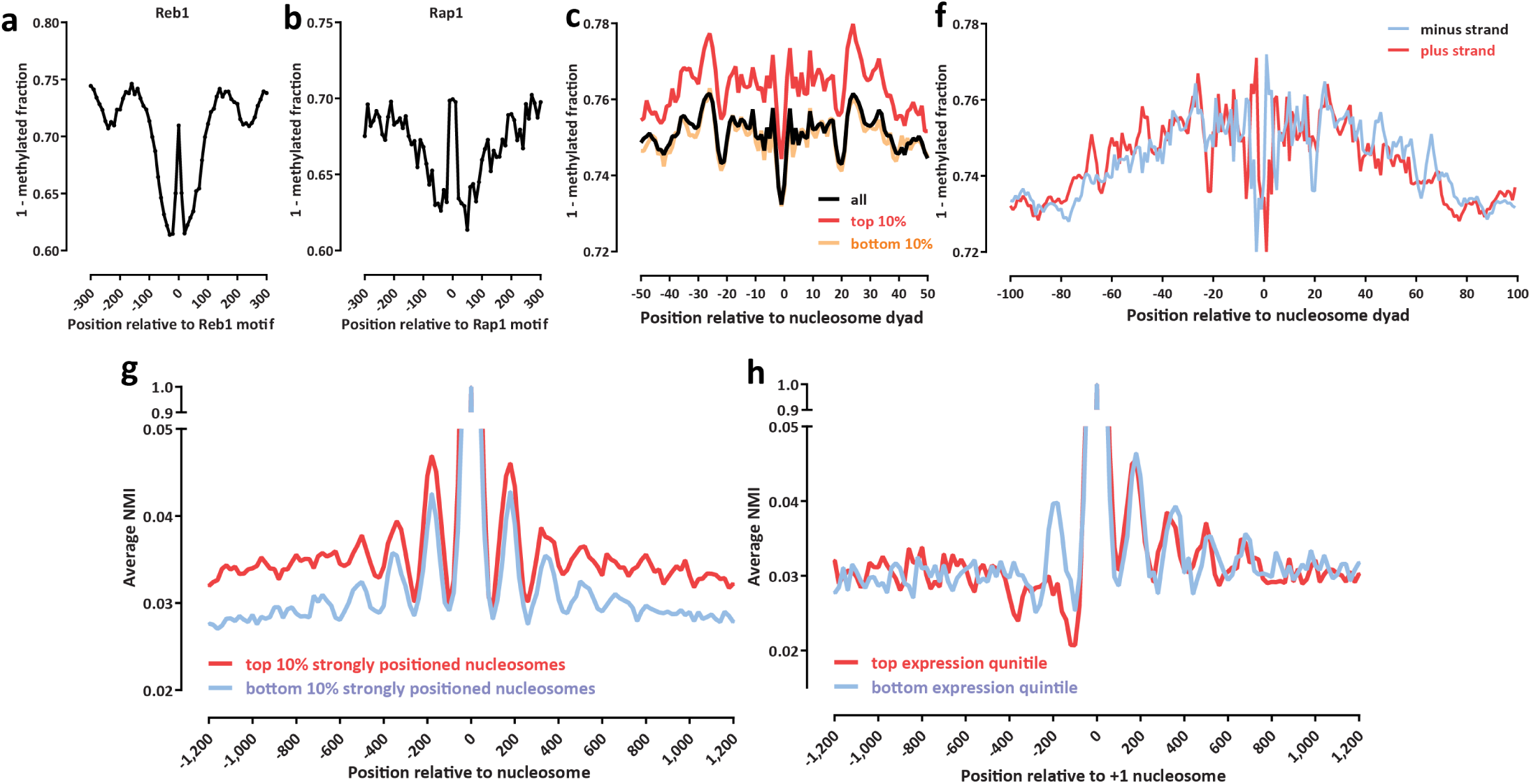
SMAC-seq provides a high-resolution strand-specific view of genomic occupancy by DNA-binding proteins and complexes. (a-b) SMAC-seq allows for precise footprinting of transcription factor binding events. Shown is aggregate genome-wide SMAC-seq signal around occupied (as measured by ChIP-exo) Reb1 (a), and Rap1 (b) sequence recognition motifs. (c) SMAC-seq profiles around positioned nucleosome dyads reveal increased accessibility in the dyad and increased protection at the points of contact with the nucleosome (see Supplementary Figure 16 for additional details.) (d) SMAC-seq provides a strand-specific view of nucleosome occupancy and reveals differential accessibility between the two DNA strands depending on their position on the nucleosomal particle. (e-f) Coordination between the positions of individual nucleosomes at the level of single chromatin fibers. (e) Shown is the average normalized mutual information between each strongly or poorly positioned nucleosome in the yeast genome and its immediate genomic neighborhood (measured for windows of 10 bp length tiling at every genomic position centered on the nucleosome dyad). (f) Shown is the average normalized mutual information between each +1 nucleosome and its immediate genomic neighborhood in highly expressed and in mostly silent genes (measured for windows of 10-bp length tiling at every genomic position centered on the +1 nucleosome dyad).

We then explored global SMAC-seq signal around positioned nucleosomes (Figure 4e-f and Supplementary Figure 16). We find higher accessibility immediately at the dyad point, in contrast to the points on the nucleosome located two DNA helical turns away in each direction (Supplementary Figure 16). The same pattern was observed for all nucleosomes irrespective of the strength of positioning; the difference between strongly positioned nucleosomes is the overall strength of protection from methylation (Figure 4e).

To further explore the limits of SMAC-seq’s resolution, we studied methylation patterns around positioned nucleo somes in more detail (Figure 4e-f and Supplementary Figure 16). We find remarkably higher accessibility immediately at the dyad point, in contrast to the points of contact of the nucleosome with the DNA two helical turns away in each direction (Supplementary Figure 16). The same pattern was observed for all nucleosomes irrespective of the strength of positioning (Figure 4e).

Because SMAC-seq directly maps accessibility independently on individual DNA strands, we next aimed to quantify strand-specificity in DNA accessibility within well-positioned nucleosomes. We observe a striking strand asymmetry in DNA accessibility around the nucleosome particle (Figure 4f), especially within the dyad and at the points two helical turns away from the dyad. The magnitude of these differences in average methylation levels are similar to those observed between nucleosomes and flanking linker regions. Thus SMAC-seq reveals significant heterogeneity in DNA’s accessibility potential within the nucleosomal particle, which has important implications for understanding how transcription factors interact with the genome in vivo, in particular in light of recent studies demonstrating that certain classes of TFs may preferentially occupy nucleosomal DNA ^37^.

### SMAC-seq reveals distal co-accessibility patterns in the genome

We next examined co-accessibility patterns in the yeast genome. We first aimed to measure correlation between the positions of individual nucleosomes. Average NMI profiles centered on positioned nucleosomes reveal detectable correlation between nucleosome positions up to three to four nucleosomes away from an individual positioned nucleosome (Figure 4g; Supplementary Figure 45), with strongly positioned nucleosomal particles exhibiting stronger overall correlation over larger distances. These observations are consistent with a model whereby the restrictions on positioning that nucleosomes impose on each result in correlation between their protection footprints on short distances until random positional fluctuations of individual nucleosomes in the chromatin fiber eventually dephase this correlation signal on longer scales.

We then examined co-accessibility patterns in the vicinity of promoters (Figure 4h). Expressed yeast genes are characterized by a nucleosome-depleted/free region (NFR) upstream of the TSS and a well positioned +1 nucleosome. NMI profiles centered on the +1 nucleosome show significant differences between highly expressed and silent genes. While correlation patterns decay downstream of the TSS similarly for both groups of genes, highly expressed genes exhibit an inverse accessibility correlation pattern upstream of the TSS. The NMI profile for highly expressed genes also exhibits inverse accessibility correlation not just with the NFR but at a distance of at least one nucleosome beyond it.

Actively transcribed yeast genes often exist in a looped conformation, in which the promoter and termination regions are brought in physical proximity, potentially helping to enforce transcriptional directionality^38,39^. Given this physical coupling, we wondered if correlation also exists between chromatin accessibility around gene ends. SMAC-seq data reveals low levels of correlation between the NFR and the accessible region in the 3′ end of genes, and stronger correlation between positioned nucleosomes in these locations (Supplementary Figure 46). The correlation between the accessibility in the NFR and the 3′ end is increased for highly expressed genes and decreased for silent genes, suggesting that transcriptional activity and looping may help more strongly position nucleosomes at the beginning and end of transcribed regions; the decreased correlation between the nucleosome-depleted areas could be explained by transcriptional activity and dynamic regulatory occupancy leading to less stable protection patterns at each end that are accordingly less well correlated with one other.

We next assessed coordinated accessibility between yeast TSSs. To this end, we devised an explicit test of coordinated coaccessibility based on splitting reads into separate pieces, randomly reassembling them, then deriving an empirical coaccessibility distribution (see the Methods for details). Using this approach we identified 1,115 TSS pairs as significantly correlated out of 19,578 pairs covered with ≥100 reads in our initial sample (Supplementary Figure 47). Of these, 560 were located a distance ≥ 1 kb from each other. An example of significantly coordinated accessibility (between the *GAL10* and *GAL1* genes) is shown in (Supplementary Figure 48). As one possible mechanism for generating correlated accessibility is increased frequency of physical association in 3D space, we used publicly available Micro-C ^40^ data to assess whether promoters exhibiting coordinated accessibility are more often physically interacting with each other. Indeed, we observe that significantly coaccessible promoters interact more frequently than noncoaccessible promoters at a similar distance (Supplementary Figure 49).

### SMAC-seq charts dynamic coordinated changes in chromatin accessibility in the course of the yeast stress response

To monitor chromatin accessibility at long-range scales during dynamic changes in gene regulatory activity brought about by environmental stimuli, we carried out SMAC-seq experiments during a time course of diamide treatment. Diamide oxidizes thiols in proteins, resulting in the generation of disulfides and activation of the stress response pathway, leading to changes in the expression of several hundred genes ^41^. Stress response in yeast is mediated through the action of the Hsf1 transcription factor (with the Msn2/4 proteins also playing a key role). Hsf1 binds to its cognate genes when activated upon stress leading to their transcriptional activation^42^.

We performed SMAC-seq at 0, 30 and 60 minutes after diamide treatment of yeast cells, as well as RNA-seq, ATAC-seq, and ChIP-seq for RNA Pol2, the elongating version of Pol2 (Pol2pS2), and a V5-tagged version of HSF1 (RNA-seq and ATAC-seq data was also collected at 15 and 45 minutes; Figure 5a). We observe several hundred genes exhibiting strong changes in gene expression during the time course (Supplementary Figure 50), and strong induction of Hsf1 occupancy at hundreds of sites in the genome (Supplementary Figure 51). SMAC-seq data at 30 minutes post-diamide treatment shows strong footprinting over Hsf1 motifs within induced Hsf1 binding sites.

**Figure 5:**
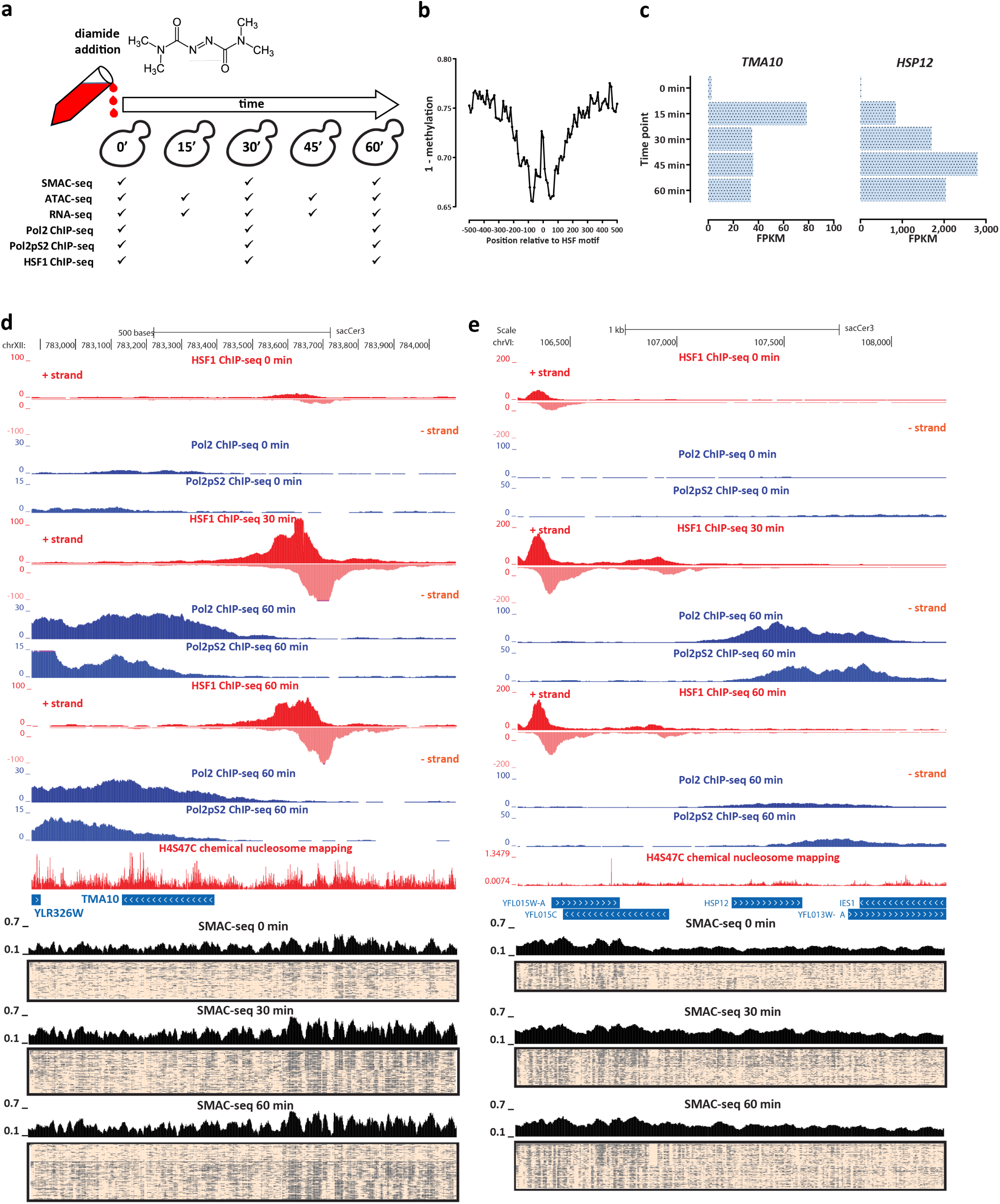
Coordinated changes in chromatin accessibility and nucleosomal occupancy during the yeast stress response. (a) Experimental outline. Yeast cells were treated with diamide, then SMAC-seq and other functional genomic assays where carried out at 15- or 30-minute intervals. (b) Sites occupied by the HSF1 transcription factor upon its activation by the stress response pathway exhibit strong footprints in SMAC-seq data. (c) Changes in the expression of the *TMA10* gene upon diamide treatment (d) Changes in RNA Polymerase and HSF1 occupancy (measured by ChIP-seq), and of chromatin accessibility at the single molecule level in the vicinity of the *TMA10* gene during the diamide time course. (e) Changes in RNA Polymerase and HSF1 occupancy (measured by ChIPseq), and of chromatin accessibility at the single molecule level in the vicinity of the *HSP12* gene during the diamide time course.

We illustrate the dynamic patterns of chromatin accessibility that we observe upon diamide treatment using the *TMA10* and *HSP12* genes as examples in Figure 5d-e, and multiple others in Supplementary Figures 52-65. The *TMA10* and *HSP12* genes are strongly upregulated at 15 minutes after diamide treatment; TMA’s expression subsequently declines somewhat and stabilizes (Figure 5c) while that of *HSP12* continues to increase up to the 45-minute time point. SMAC-seq reveals a relatively modest level of accessibility usptream of these genes before diamide treatment. However, at 30 minutes and upon Hsf1 binding, dramatic changes are evident. Nearby nucleosomes appear to be evicted in many cells, and nucleosome depletion is also observed at increased levels within the gene bodies, where ChIP-seq data for Pol2 and Pol2pS2 shows highly active transcription. At 60 minutes we observe dampening of this response, with the accessible fraction of reads decreasing, more strongly so around the TMA10 gene, whose expression decreases earlier than that of HSP12. Examination of NMI co-accessibility maps (Supplementary Figures 52-65) frequently shows loss of correlation between positioned nucleosomes within and upstream of activated gene bodies as a result of response to diamide treatment, consistent with an increased movement of nucleosomal particles due to the activities of transcribing polymerase molecules and chromatin remodelers.

## Discussion

SMAC-seq is a novel single-molecule method for profiling chromatin accessibility within individual chromatin fibers at a multikilobase resolution that leverages the power of nanopore sequencing to detect DNA modifications and the preference of DNA methyltransferases for chemically modifying accessible DNA. We show that SMAC-seq generates accessibility signals similar to those of widely used methods such as DNase-seq and ATAC-seq while also opening new windows into the long-range structure of eukaryotic chromatin. SMAC-seq enables the simultaneous profiling of nucleosome positioning and accessible chromatin on a truly genome-wide scale (including repetitive regions that are poorly mappable with short reads), the assessment of the absolute distribution of chromatin accessibility states within a population of cells, and the identification of pairs of genomic loci that exhibit significant correlations in co-accessibility.

Our initial work on SMAC-seq focused on applications in *S. cerevisiae* because of its modest genome size. Extending SMAC-seq to larger genomes will require significantly increased sequencing throughput, or a method for selective enrichment of a collection of individual loci. Fortunately, both of these approaches appear feasible in the near term, as nanopore sequencing throughput is increasing rapidly, while methods for selective enrichment of genomic regions for nanopore sequencing are also becoming available ^43^. Read length increases will also be useful, in particular for correlating the activity of distal regulatory elements to their cognate promoters, which can often be located many tens of kilobases apart in mammalian genomes.

Improvements to base calling accuracy constitute another major area of potential future advances. At the time of our analysis, Tombo was the only readily available algorithm for calling m^6^A, but, as discussed above, its base calls exhibit on the order of 20-25% error rates for all three bases combined. Results obtained with the other widely used methylation-aware base caller, Nanopolish, which is currently capable of calling only CpG and GpC methylation, show ~15% error on CpG and GpC calls. The major barrier to base-calling improvements is the lack of sets of ground truth controls that can be used to train base calling algorithms. Pools of DNA templates with individual modifications in well-defined yet highly diverse base pair contexts are the ideal training set for developing accurate such models. Alternatively, the introduction of tags bulkier than a simple methyl group ^44^ may provide a much stronger modulation of electric current through the nanopore than does simple methylation, enabling much more reliable modified base calls and accessibility evaluation.

We also anticipate an increase in the diversity of the DNA modifying enzymes available to carry out variations of SMAC-seq and related assays. In mammalian systems, endogenous 5mC methylation occurs primarily in the CpG context, thus it is not possible to use M.SssI for mapping accessibility, leaving only GpC as an option for traditional NOME-seq, bisulfite-based assays. Fortunately, the bulk of SMAC-seq’s increase in resolution is derived from m^6^A (Supplementary Figure 1), so this will not be a significant obstacle to its widespread application in mammalian systems. However, there are species where m^6^A occurs endogenously and is strongly correlated with patterns of accessibility and nucleosome positioning (e.g. *Chlamydomonas* ^45^ and *Tetrahymena* ^46,47^). Alternative methylation strategies such as 4mC methyltransferases^48^, cytidine deaminases ^49^, or the conversion of thymine to modified bases such as 5hydroxymethyluracil (dhmU)^50^ are areas of potential future exploration.

Finally, we note that there is considerable scope for integration of SMAC-seq-type approaches with other measurements of the physical genome and the epigenome, especially once the improvements in accuracy outlined above are achieved. We anticipate the potential for obtaining simultaneous measurements of accessibility and nucleosomal positioning together with endogenous DNA methylation, general and specific protein occupancy, chromatin interactions, DNA replication, and other features on a multikilobase scale and within single molecules. In principle, similar approaches may also be applicable to individual RNA molecules. We expect long-read single-molecule approaches to provide an important new class of tools for the study of the functional and physical organization of genomes in the coming years.

## Materials and Methods

Except for when explicitly stated otherwise, all analyses were carried out using custom-written Python or R scripts.

### Cell lines and cell culture

The BY4741 *S. cerevisiae* strain (a kind gift from Ji-Ping Wang and Xiaozhong Wang) was used for all experiments except for Hsf1 ChIP-seq experiments where MS143 (H4S47C Hsf1-V5::HphMX6, this study) was used. MS143 was generated by PCR-based C-terminal tagging of Hsf1 with the V5 epitope. Hsf1-V5 tagging was confirmed by colony PCR and western blotting. For all experiments, except the initial one (“Sample 1”), cells were grown in YPD media (30 °C) to OD~0.8 before collection.

### SMAC-seq experiments

#### Enzymatic treatment of chromatin

We developed and optimized SMAC-seq using the equivalent of 1 × 10^6^ human cells, which in the case of *S. cerevisiae* translates in to 2.5 × 10^8^ (the size of the haploid human genome is ~3 × 10^9^bp while that of *S. cerevisiae* is 1.2 × 10^6^bp). As yeast cells have a cell wall, we adapted the spheroplasting protocol previously used for carrying out ATAC-seq in yeast cells ^51^ for our SMAC-seq experiments.

Yeast cells in log phase (OD_660_ ≤ 1.0) were first centrifuged at 13,000 rpm for 1 minute, then washed with 100 *µ*L Sorbitol Buffer(1.4 M Sorbitol, 40 mM HEPES-KOH pH 7.5, 0.5 mM MgCl_2_), and centrifuged again at 13,000 rpm for 1 minute. Cells were then spheroplasted by resuspending in 200 *µ*L Sorbitol Buffer with DTT added at a final concentration of 10 mM and 0.5 mg/mL 100T Zymolase, followed by incubating for 5 minutes at 30 °C at 300 rpm in a Thermomixer. The pellet was centrifuged for 2 minutes at 5,000 rpm, washed in 100 *µ*L Sorbitol Buffer, and centrifuged again at 5,000 rpm for 2 minutes.

Cells were then resuspended in 100 *µ*L ice-cold Nuclei Lysis Buffer (10 mM Tris pH 7.4, 10 mM NaCl, 3 mM MgCl_2_, 0.1 mM EDTA, 0.5% NP-40) and incubated on ice for 10 minutes. Nuclei were then centrifuged at 5000 rpm for 5 min at 4 °C, resuspended in 100 *µ*L cold Nuclei Wash Buffer (10 mM Tris pH 7.4, 10 mM NaCl, 3 mM MgCl_2_, 0.1 mM EDTA), and centrifuged again at 5,000 rpm for 5 min at 4 °C. Finally, nuclei were resuspended in 100 *µ*L M.CviPI Reaction Buffer (50 mM Tris-HCl pH 8.5, 50 mM NaCl, 10 mM DTT).

Nuclei were then first treated with M.CviPI + EcoGII by adding 200 U of M.CviPI (NEB) and 200 units of EcoGII (NEB), SAM at 0.6 mM and sucrose at 300 mM, and incubating at 30 °C for 7.5 min. After this incubation, 128 pmol SAM and another 100 U of enzymes were added, and a further incubation at 30 °C for 7.5 min was carried out. Immediately after, M.SssI treatment followed, by adding 60 U of M.SssI (NEB), 128 pmol SAM, MgCl_2_ at 10 mM and incubation at 30 °C for 7.5 min.

The reaction was stopped by adding an equal volume of Stop Buffer (20 mM Tris-HCl pH 8.5, 600 mM NaCl, 1% SDS, 10 mM EDTA).

#### High-molecular weight DNA isolation

HMW DNA was isolated using the MagAttract HMW DNA Kit (Qiagen; cat # 67563) following the manufacturer’s instructions.

#### Enzymatic treatment of naked DNA

Naked DNA was treated under exactly the same conditions as chromatin except that the reaction volume and enzyme amounts were reduced in half. HMW DNA was purified as described above

### SMAC-seq analysis

#### Nanopore sequencing

HMW DNA was converted into libraries using the Ligation Sequencing Kit 1D (Oxford Nanopore Technologies, SQK-LSK108) following the manufacturer’s instructions. Nanopore sequencing was carried out on R9.4 MinION flow-cells (Oxford Nanopore Technologies) for up to 48 hours.

#### Nanopore base calling

Nanopore events were converted to DNA sequence using Albacore (V2.3.3) using default settings. Reads were resquiggled using Tombo ^26^, version 1.3, using the sacCer3 reference genome. Methylated bases were identified using Tombo in the “de novo model” mode.

#### Aggregation of accessibility information over multibasepair windows

Even with the addition of m^6^A methylation, the resolution of SMAC-seq still does not cover every nucleotide in the genome, and it varies substantially between different locations depending on local sequence content differences. In addition to that, nanopore base calling is still far from being a fully resolved problem, and even more so in methylation-aware mode. For these reasons, for many of the analyses described in this study we aimed at assigning aggregate accessibility scores over windows, taking the totality of the available evidence into account, thus obtaining more reliable, if coarser-grained, views of accessibility patterns along the genome. We used a Bayesian approach to carry out aggregation, as follows.

For a given window of width *w* in the genome, specified by coordinates *c, i, i* + *w* (where *c* denotes the chromosome, and *i* the leftmost coordinate of the window), and for all reads *r ∈ R_c,i,i_*_+*w*_ fully spanning the window, we obtain all Tombo probabilities *p*_*r*,(*c,j*)_ such that *j* ∈ [*i, i* + *w*) for sequence contexts CpG, GpC and A on the corresponding genomic strand. We use a Beta prior *B*(*α*,*β*), with *α* = *β* = 10, which we then updated based on each probability *p*_*r*,(*c,j*)_ for all *j* ∈ [*i, i* + *w*). The final binary accessibility score *p*_*r*,(*c,i,i*+*w*)_ for read *r* and window *c, i, i* + *w* is determined by the final state of the prior.

#### Read filtering

As discussed above, we sometimes observe a population of reads that are fully methylated across their whole length or over large segments of it. There reads most likely derive from dead cells, as our initial experiment, which was carried out on a very dense yeast population containing a substantial number of dead cells, exhibited much higher proportion of such reads compared to subsequent experiments using early log-phase cells. In order to remove such potentially artifactual reads, “filtered” sets of reads were obtained by removing all reads containing a ≥1-kbp stretch that is ≥75% methylated (while also filtering out reads shorter than 1 kb).

#### Read clustering

For most analyses presented in this manuscript, the `tglkmeans` package was used to cluster SMAC-seq reads (implemented in R, https://bitbucket.org/tanaylab/tglkmeans. In addition, the hierarchical clustering implementation in scipy was also used in certain cases.

### Co-accessibility assessment using Normalized Mutual Information

To evaluate co-accessibility patterns along the genome, we applied a Normalized Mutual Information as follows. Each chromosome in the genome *c* was split into windows of size *w*. For each such window (*c, i, i*+*w*), we identified the maximum range to the right of it, (*c, j, j*+*w*) such that the span (*c, i, j* + *w*) is covered by ≥ *M* reads. All reads spanning (*c, i, j* + *w*) were then extracted and subsampled down to *M* reads (usually *M* = 100, unless specified otherwise). Accessibility scores were then aggregated and binarized as described above for all windows located in the span (*c, i, j*+*w*), and for all *M* reads fully spanning it, resulting in a local co-accessibility matrix *LCM* of size *M* × (*j* + *w−i*)*/w*. We then calculated Normalized Mutual Information scores for each pair of columns *LCM_k_* and *LCM_l_* as follows:

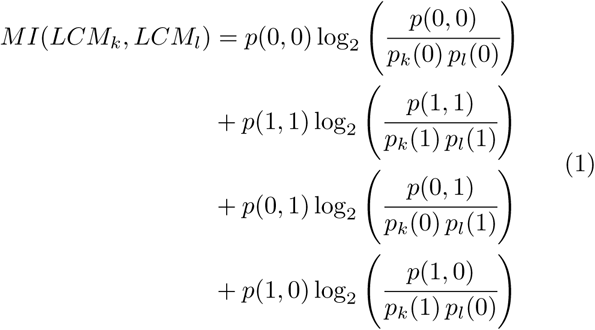

MI scores were then normalized and rescaled in the interval (−1, 1):

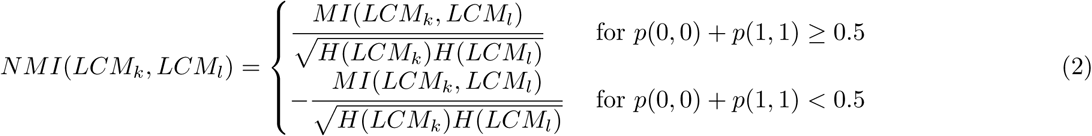

Where *H* refers to the entropy of each individual distribution.

For computational efficiency, local NMI matrices were calculated for even-sized (50kb) evenly spaced (every 10kb) tiles of the genome. The entries of the general genome-wide NMI matrix were then calculated as the average of all local NMI matrices containing each entry.

#### Testing for coordinated accessibility

Coordinated accessibility was evaluated as follows. For each pair of locations (*c, i*_1_*, i*_1_ + *r*_1_) and (*c, i*_2_*, i*_2_ + *r*_2_) (usually *r*_1_ = *r*_2_), a minimum number of reads *N* was required that fully spans the (*c, i*_1_*, i*_2_ + *r*) interval. All such reads were then obtained for each pair, and then subsampled multiple times down to *N* reads (in order not to introduce bias in coordinated accessibility tests arising due to differential read coverage between locations closer/further apart). For each subsampling, the fraction of accessible regions *p*_1_ and *p*_2_ was estimated for each of the two locations using the Bayesian procedure described above, as well as the distribution of joint accessibilities over the four states (0, 0), (1, 0), (0, 1), and (1, 1). The two halves of the reads were then virtually split in half and recombined for a total of 10^3^ random combinations. The empirical distribution 𝒩(*µ, σ*) of the four states was then estimated from these random combinations, where *µ* = *N* ∗ (*p*_(0,0)_ + *p*_(1,1)_) if *p*_(0,0)_ + *p*_(1,1)_ *>* 0.5 and µ = *N* ∗ (*p*_(1,0)_ + *p*_(1,0)_) if *p*_(0,0)_ + *p*_(1,1)_ *≤* 0.5. Empirical coordinated accessibility *p*-values were then estimated based on the observed counts |(0, 0)|+|(1, 1)| if |(0, 0)|+|(1, 1)| *>* 0.5∗*N* or |(0, 1)|+|(1, 0)| if |(0, 0)|+|(1, 1)| ≤ 0.5 ∗ *N*. Bonferroni correction was applied to account for multiple hypothesis testing.

### dSMF and Bisulfite sequencing

Illumina measurements of CpG and GpC methylation levels were carried out using the PBAT^52^ with modifications. HMW DNA (~500 ng) was bisulfite converted using the EZ DNA Methylation-Lightning Kit (Zymo, Cat # D5030) by mixing 20 *µ*L of purified DNA (~500 ng) with 130 *µ*L DNA Methylation Lightning Conversion reagent and incubating at 98 °C for 8 minutes and then at 64 °C for 60 minutes. Bisulfite converted DNA was then cleaned up using the EZ DNA Methylation-Lightning Kit following manufacturer’s instructions.

First strand synthesis was carried out by mixing 20 *µ*L bisulfite converted DNA, 19.75 *µ*L H_2_O, 5 *µ*L 10× Blue Buffer (ThermoFisher), 1.25 *µ*L 10 mM dNTP (NEB), and 4 *µ*L custom-designed biotinylated adapter. Samples were then incubated at 94 °C for 5 minutes, and at 4 °C for 5 minutes, after which 1.5 *µ*L Klenow (3′ → 5′ exo minus; MCLab) were added, and the reaction was incubated at 4 °C for 15 minutes, at 37 °C for 90 minutes, and at 70 °C for 5 minutes. First-strand reaction cleanup was carried out using 50 *µ*L AMPure XP beads (Beckman Coulter); DNA was eluted 50 *µ*L EB buffer.

Biotinylated DNA was captured on streptavidin beads. A total of 20 *µ*L streptavidin Dybaneads M-280 (ThermoFisher) per sample were added to a PCR tube, separated on a magnet and then resuspended in 50 *µ*L 2× BW(Li) buffer (6.3 g LiCl, 0.5 mL Tris-HCL pH 8.0, and 0.1 mL 500 mM EDTA for 50 mL total volume), to which the 50 *µ*L of eluted first-strand reaction DNA was added. Beads were then incubated at room temperature for 30 minutes, washed with 180 *µ*L 2× BW(Li) buffer, twice with 0.1 N NaOH (by resuspending well and incubating at room temperature for 2 minutes), washed again with 180 *µ*L 2× BW(Li) buffer, then with 180 *µ*L 10 mM Tris-HCL pH 7.5.

Second-strand synthesis was carried out by resuspending streptavidin beads in the following reaction mix: 5 *µ*L 10 × Blue Buffer, 1.25 *µ*L 10 mM dNTPs, 39.75 *µ*L H_2_O, 4 *µ*L custom-designed second-strand adapter. Samples were then incubated at 94 °C for 5 minutes, and at 4 °C for 5 minutes, after which 1.5 *µ*L Klenow (3′ → 5′ exo minus) were added, followed by further incubation at 4 °C for 15 minutes, at 37 °C for 30 minutes, and at 70 °C for 5 minutes.

Beads were separated on magnet and the chase reaction was carried out by resuspending in a mix of 5 *µ*L 10× Thermo Pol Buffer, 1.25 *µ*L 10 mM dNTPs, 43.5 *µ*L H_2_O, and 1 *µ*L *Bst* DNA Polymerase Large Fragment (NEB). Samples were incubated at 65 °C for 30 minutes, then again separated on magnet.

PCR was performed on beads in 50 *µ*L reactions composed of 25 *µ*L 2× NEB Next PCR Master Mix, 20 *µ*L H_2_O, 2.5 *µ*L i7 and 2.5 *µ*L i5 primers (both custom-designed), with initial extension at 72 °C for 3 min, denaturation at 98 °C for 30 sec, 15 cycles of 98 °C for 10 sec, 63 °C for 30 sec, and 72 °C for 30 sec, and final extension at 72 °C for 5 min. PCR reactions were cleaned up and size-selected using AMPure XP beads.

Libraries were sequenced on Illumina NextSeq or MiSeq instruments, as 2×75mers or 2×300mers, respectively.

### dSMF data processing

Bisulfite reads were trimmed using cutadapt (version 0.16) and Trim Galore (version 0.4.4), using the following settings (taking into account that the bisulfite sequencing libraries are generated with the PBAT protocol): `–clip_R1 9 –clip_R2 9 –three_prime_clip_r1 6 –three_prime_clip_r2 6 –paired`. Trimmed reads were the mapped to the sacCer3 version of the yeast genome using Bismark^53^ (version 0.19.0) with the following settings: `–bowtie2 –pbat`. Methylation calls were extract using the `bismark_methylation_extractor` program within Bismark and the following settings: `-s –no_overlap –comprehensive –merge_non_CpG –cytosine_report –CX`.

### ATAC-seq

ATAC-seq was carried out on the same nuclei isolated for SMAC-seq as described above (before resuspension in M.CviPI Reaction Buffer), by resuspending nuclei with 25 *µ*L 2× TD buffer (20 mM Tris-HCl pH 7.6, 10 mM MgCl_2_, 20% Dimethyl Formamide), 2.5 *µ*L transposase (custom produced) and 22.5 *µ*L nuclease-free H_2_0, and incubating at 37 °C for 30 min in a Thermomixer at 1000 RPM. Transposed DNA was isolated using the DNA Clean & Concentrator Kit (Zymo, cat # D4014) and PCR amplified as described before ^54^. Libraries were then sequenced on a Illumina NextSeq instrument as 2×36mers or as 2×75mers.

### ATAC-seq data processing

Demultipexed fastq files were mapped to the `sacCer3` assembly of the `S. cerevisiae` genome as 2×36mers using Bowtie ^55^ with the following settings: `-v 2 -k 2 -m 1 –best –strata`. Duplicate reads were removed using `picard-tools` (version 1.99).

### ChIP-seq experiments

Cell lysis and ChIP reactions were performed as previously described ^56^ with minor modifications. Cells were fixed with 1% formaldehyde for 20 minutes (Rpb1-CTD and Rbp1-CTD-S2P ChIP) or 30 minutes (Hsf1-V5 ChIP) and quenched with 0.125 M glycine for 5 minutes. A total of ~50 ODs of cells were used per Rpb1-CTD or Rpb1-CTDS2P ChIP and ~300 ODs per Hsf1-V5 ChIP. Fixed cell were washed 2× in cold 1× PBS, pelleted and stored at −80 °C. Pellets were lysed in 300 *µ*L FA lysis buffer (50 mM HEPES–KOH pH 8.0, 150 mM NaCl, 1 mM EDTA, 1% Triton X-100, 0.1% sodium deoxycholate, 1 mM PMSF, Roche protease inhibitor) with ~1 mL ceramic beads on a Fastprep-24 (MP Biomedicals). The entire lysate was then collected and adjusted to 1 mL with FA lysis buffer before sonication with a 1/8’ microtip on a Q500 sonicator (Qsonica) for 14 minutes (10 seconds on, 20 seconds off). The sample tube was held in a −20 °C 80% ethanol bath throughout sonication to prevent sample heating. After sonication, cell debris was pelleted and the supernatant was retained for ChIP. For each ChIP reaction, 30 *µ*L Protein G Dynabeads (Invitrogen) were blocked (PBS + 0.5% BSA), prebound with 5-10 *µ*L antibody (8wG16 Rpb1-CTD, Abcam cat # ab817); 3E10 Rpb1-CTD-S2P, Milipore cat # 04-1571-1) or SV5-Pk1 (anti-V5, BioRad cat # MCA1360G)) and washed 1× with PBS before incubation with supernatant (4 °C, overnight). Dynabeads were then washed (5 minutes per wash) 3× in FA lysis buffer, 3× in high-salt FA lysis buffer (50 mM HepesKOH pH 8.0, 500 mM NaCl, 1 mM EDTA, 1% Triton X-100, 0.1% sodium deoxycholate, 1 mM PMSF), 1× in ChIP wash buffer (10 mM TrisHCl pH 7.5, 0.25 M LiCl, 0.5% NP-40, 0.5% sodium deoxycholate, 1 mM EDTA, 1 mM PMSF) and 1× in TE wash buffer (10 mM TrisHCl pH 7.5, 1 mM EDTA, 50 mM NaCl). DNA was eluted from the beads in ChIP elution buffer (50 mM TrisHCl pH 7.5; 10 mM EDTA; 1% SDS) at 65 °C for 20 min. Eluted DNA was incubated at 65 °C overnight to reverse crosslinks, before treatment with RNAse A (37 °C, 1 hour) and then Proteinase K (65 °C, 2 hours). DNA was purified using the ChIP DNA Clean & Concentrator kit (Zymo Research). Sequencing libraries were generated using the NEBNext Ultra II DNA Library Prep kit (NEB Cat # E7645) and sequenced on a Illumina NextSeq instrument as 2×36mers or as 2×75mers.

### ChIP-seq data processing

Demultipexed fastq files were mapped to the sacCer3 assembly of the `S. cerevisiae` genome as 2×36mers using Bowtie ^55^ with the following settings: `-v 2 -k 2 -m 1 –best –strata`. Duplicate reads were removed using `picard-tools` (version 1.99). Hsf1 peaks were called using MACS2 ^57^ (version 2.1.0) with the following settings: `-g 12000000 -f BAMPE`.

### Multiread-preserving alignment and normalization

Multiread-preserving alignment and track generation was carried out by mapping reads to the *sacCer3* assembly of the `S. cerevisiae` genome using Bowtie ^55^ with the following settings: `-v 2 -a –best –strata`. Each alignment was then given a weight inversely proportional to the number of locations that the read maps to i.e. each position’s score was normalized to RPMs as follows:

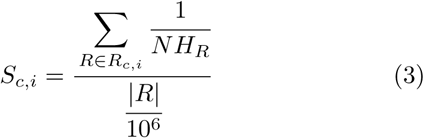

Where *NH_R_* is the number of locations in the genome a read maps to.

### RNA-seq experiments

Cells (1 mL) were pelleted and flash frozen in liquid N_2_. Pellets were resuspended in 300 *µ*L TRIzol and lysed with ~1 mL ceramic beads on a Fastprep-24 (MP Biomedicals). Cell debris were pelleted and RNA was extracted from the supernatant using the Direct-Zol RNA Microprep Kit (Zymo Research). RNA-seq libraries were generated using the NEBNext Ultra Directional RNA Library Prep Kit (NEB Cat # E7420)

### RNA-seq processing and gene expression quantification

RNA-seq reads were mapped to the yeast genome as 1×50mers (external datasets) or 2×75mers (diamide experiments) using TopHat version 2.0.8 ^58^. Gene-level quantifications in FPKMs (Fragments Per Kilobase per Million mapped reads) were generated using Cufflinks version 2.0.2 ^58^. The mean from all replicates was taken as the expression level for each gene for subsequent analyses.

### External sequencing datasets

A number of previously published *S. cerevisiae* genomics datasets were used in this study. ChIP-exo reads and called peaks for Abf1, Cbf1, Rap1 and Reb1 were downloaded from GEO accessions GSE93662 and GSE72106. ChIP-seq data for centromeric proteins was downloaded from GEO accessions GSE31466 and GSE51949. PRO-seq and PRO-CAP data was obtained from GEO accession GSE76142. ORC ChIP-seq data was downloaded from GEO accession GSE16926. DNase-seq was downloaded from GEO accession GSE69651 while DGF data was downloaded from SRA accession SRP000620. MNaseseq data was obtained from GEO accessions GSE26493 and GSE29292, TBP ChIP-seq from GSE44200, Rpb1 ChIP-seq from GSE93190, Rpb3 ChIP-seq from GSE74787, RPC128 ChIP-seq from GSE39566, and Mediator subunits ChIP-seq from GSE95051. RNA-seq data from accession GSE85590 was also used. Except where otherwise stated, raw reads were aligned using Bowtie ^59^ with the following settings: `-v 2 –k 2 -m 1 –best –strata`, with the addition of -X 1000 for paired-end reads. Paired-end reads were aligned as 2×25mers), while single-end reads were aligned as 1×36mers. PRO-seq and PRO-CAP data was aligned as 1×16mers.

### Micro-C data and processing

Micro-C data was downloaded from GEO accession GSE68016 and processed as described in the original publication^40^.

### Transcription factor motif mapping

Transcription factor motif recognition sequences were mapped genome-wide using FIMO ^60^ (version 4.11.2 of the MEME-Suite^61^ using the CIS-BP database^62^ as a reference set of position weight matrices.

### Gene annotation update

Publicly available gene models for *S. cerevisiae* do not contain TSS and TTS information for a major fraction of genes in the genome, only including the coding (“CDS”) portions instead. As the omission of UTRs presents a problem for TSS- and TTS-centered analyses, we updated the existing gene models following the approach described previously ^51^ and the *S. cerevisiae* TIF-seq dataset from GEO accession GSE39128^63^. New TSS and TTS positions were assigned to each gene for which such information was available based on the median UTR length as measured by TIF-seq.

### Nucleosome positioning information

H4S47C ^25,64^ chemical mapping data was downloaded form GEO accessions GSE59523 and GSE36063. H3Q85C ^65^ chemical mapping data was downloaded from GEO accession GSE97290. We used the nucleosome positioning calls obtained from the original 2012 Brogaard et al. study for our analyses, after transforming them from coordinates in the `sacCer2` version of the *S. cerevisiae* genome assembly to `sacCer3` using the `liftOver` function in the UCSC Genome Browser utilities toolkit.

### Mappability tracks generation

To evaluate unique read mappability, the whole genome was tiled with reads of given length at every position. The reads were then mapped back to the genome using the same settings used to map single-end ChIP-seq reads. For every position coverage by mapped reads was calculated, and mappability was scored as the ratio between read coverage and the read length used to tile the genome.

## Author contributions

Z.S, G.K.M and N.A.S.A. conceived and designed the study. Z.S, G.K.M and N.A.S.A. performed initial experiments. Z.S., M.P.S. and G.K.M. performed diamide time course experiments. G.K.M. and Z.S. analyzed data. W.J.G. and A.K. supervised the study. G.K.M., Z.S., W.J.G. and A.K. wrote the manuscript with input from all authors.

## Acknowledgments

This work was supported by NIH grants (P50HG007735, RO1 HG008140, U19AI057266 and UM1HG009442 to W.J.G., 1UM1HG009436 to W.J.G. and A.K., 1DP2OD022870-01 and 1U01HG009431 to A.K.), the Rita Allen Foundation (to W.J.G.), the Baxter Foundation Faculty Scholar Grant, and the Human Frontiers Science Program grant RGY006S (to W.J.G). W.J.G is a Chan Zuckerberg Biohub investigator and acknowledges grants 2017-174468 and 2018-182817 from the Chan Zuckerberg Initiative. Z.S. is supported by EMBO Long-Term Fellowship EMBO ALTF 1119-2016 and by Human Frontier Science Program Long-Term Fellowship HFSP LT 000835/2017-L G.K.M. is supported by the Stanford School of Medicine Dean’s Fellowship. N.A.S.A. is funded by the Department of Defense through a National Defense Science and Engineering Grant and by a Stanford Graduate Fellowship. The authors would also like to thank members of the Greenleaf and Kundaje labs for their helpful suggestions and discussions on the subject over the course of the study.

