## Supplemental Tables and Figures for "Long-range single-molecule mapping of chromatin accessibility in eukaryotes"

### Supplementary Materials

#### Supplementary Tables

**Supplementary Table 1:** Mapping statistics for SMAC-seq datasets in this study

| sample | number reads | total bases | mean read length | median read length |
|---|---|---|---|---|
| Sample 1 | 847,262 | 3,475,258,237 | 4,102 | 1,474 |
| diamide 0 min rep1 | 2,462,311 | 4,018,221,647 | 1,632 | 722 |
| diamide 30 min rep1 | 1,127,937 | 3,376,196,280 | 2,993 | 1,254 |
| diamide 60 min rep1 | 1,543,217 | 4,187,296,821 | 2,713 | 1,171 |
| diamide 0 min rep2 | 1,375,008 | 2,232,750,288 | 1,624 | 678 |
| diamide 30 min rep2 | 2,266,191 | 6,046,929,720 | 2,668 | 977 |

**Supplementary Table 2:** Mapping and QC statistics for ATAC-seq datasets used in this study

| Dataset | Complexity | TSS ratio | Read Length | uniquely mapped deduplicated reads | Raw fragments |
|---|---|---|---|---|---|
| L464 Diamide 0 min rep1 | 0.79 | 1.38 | $2 \times 36$ | 2,284,992 | 2,363,608 |
| L465 Diamide 0 min rep2 | 0.79 | 1.36 | $2 \times 36$ | 2,383,094 | 2,409,446 |
| L466 Diamide 15 min rep1 | 0.77 | 1.49 | $2 \times 36$ | 1,907,268 | 1,760,891 |
| L467 Diamide 15 min rep2 | 0.75 | 1.42 | $2 \times 36$ | 3,415,058 | 3,058,834 |
| L468 Diamide 30 min rep1 | 0.72 | 1.49 | $2 \times 36$ | 3,223,114 | 3,414,835 |
| L469 Diamide 30 min rep2 | 0.74 | 1.41 | $2 \times 36$ | 3,081,970 | 2,846,193 |
| L470 Diamide 45 min rep1 | 0.79 | 1.37 | $2 \times 36$ | 2,411,938 | 2,457,100 |
| L471 Diamide 45 min rep2 | 0.75 | 1.40 | $2 \times 36$ | 3,135,316 | 2,885,651 |
| L472 Diamide 60 min rep1 | 0.74 | 1.45 | $2 \times 36$ | 3,205,726 | 3,120,294 |
| L473 Diamide 60 min rep2 | 0.76 | 1.43 | $2 \times 36$ | 2,387,398 | 2,244,141 |

**Supplementary Table 3:** Mapping and QC statistics for ChIP-seq datasets used in this study

| Dataset | Complexity | Read Length | Uniquely mapped deduplicated reads | Raw fragments |
|---|---|---|---|---|
| L482 Diamide 0 min Input | 0.88 | $2 \times 36$ | 8,776,410 | 5,488,322 |
| L483 Diamide 0 min Pol2 | 0.87 | $2 \times 36$ | 4,944,766 | 2,894,414 |
| L484 Diamide 0 min Pol2pS2 | 0.80 | $2 \times 36$ | 5,834,864 | 3,434,723 |
| L485 Diamide 0 min HSF1-V5 Input | 0.90 | $2 \times 36$ | 6,089,572 | 3,850,484 |
| L486 Diamide 0 min HSF1-V5 | 0.93 | $2 \times 36$ | 2,540,876 | 1,726,223 |
| L487 Diamide 30 min Input | 0.86 | $2 \times 36$ | 10,052,178 | 6,212,240 |
| L488 Diamide 30 min Pol2 | 0.84 | $2 \times 36$ | 6,763,128 | 3,961,384 |
| L489 Diamide 30 min Pol2pS2 | 0.84 | $2 \times 36$ | 6,332,462 | 3,901,689 |
| L490 Diamide 30 min HSF1-V5 Input | 0.92 | $2 \times 36$ | 4,587,466 | 2,831,871 |
| L491 Diamide 30 min HSF1-V5 | 0.89 | $2 \times 36$ | 3,324,498 | 2,054,869 |
| L492 Diamide 60 min Input | 0.90 | $2 \times 36$ | 5,812,736 | 3,539,630 |
| L493 Diamide 60 min Pol2 | 0.90 | $2 \times 36$ | 3,774,106 | 2,204,126 |
| L494 Diamide 60 min Pol2pS2 | 0.85 | $2 \times 36$ | 4,873,244 | 2,924,345 |
| L495 Diamide 60 min HSF1-V5 Input | 0.87 | $2 \times 36$ | 7,683,586 | 4,679,564 |
| L496 Diamide 60 min HSF1-V5 | 0.91 | $2 \times 36$ | 1,094,048 | 698,664 |

**Supplementary Table 4:** Mapping and QC statistics for RNA-seq datasets used in this study

| Dataset | Complexity | Read Length | Unique | Unique Splices | Multi | Multi Splices | Raw fragments |
|---|---|---|---|---|---|---|---|
| Diamide 0 min | 0.20 | $2 \times 75$ | 30,692,672 | 461,804 | 3,974,211 | 38,859 | 18,719,790 |
| Diamide 15 min | 0.21 | $2 \times 75$ | 26,991,043 | 182,208 | 3,217,924 | 18,788 | 15,960,277 |
| Diamide 30 min | 0.19 | $2 \times 75$ | 34,711,193 | 201,334 | 3,716,196 | 21,450 | 21,717,800 |
| Diamide 45 min | 0.22 | $2 \times 75$ | 28,668,901 | 116,773 | 3,103,601 | 19,301 | 17,832,222 |
| Diamide 60 min | 0.31 | $2 \times 75$ | 14,991,437 | 71,198 | 1,759,048 | 10,455 | 13,548,619 |
| Diamide Hsf-V5 0min | 0.19 | $2 \times 75$ | 32,981,363 | 519,649 | 4,320,132 | 35,513 | 21,535,487 |
| Diamide Hsf-V5 30min | 0.19 | $2 \times 75$ | 33,505,595 | 211,992 | 3,973,770 | 23,016 | 22,974,020 |
| Diamide Hsf-V5 60min | 0.19 | $2 \times 75$ | 40,290,524 | 196,082 | 4,698,437 | 29,628 | 28,894,301 |

### Supplementary Figures

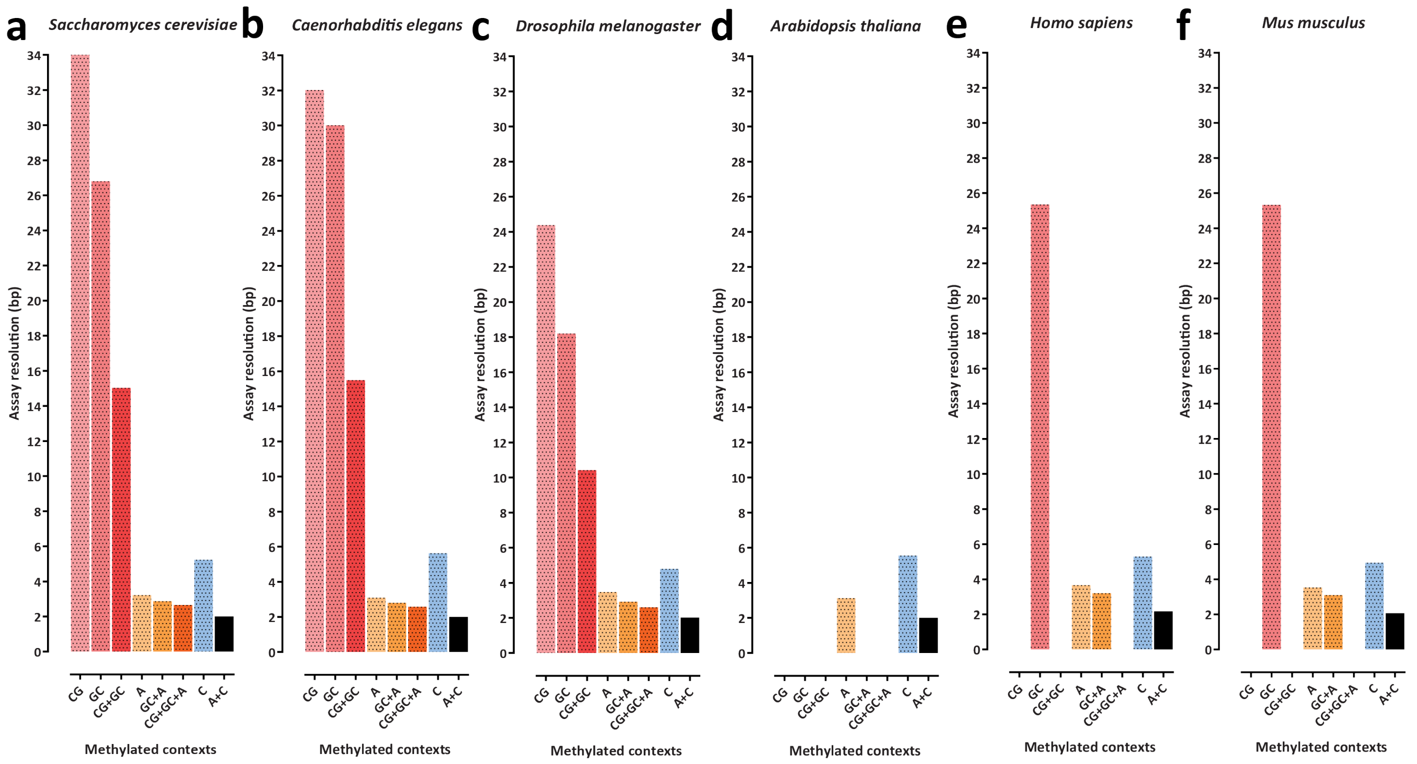


**Supplementary Figure 1: Resolution of the SMAC-seq assay in its current form and some potential other versions of it in the main model organism systems**. Note that where endogenous methylation confounds readout of accessibility, the corresponding combination of sequence contexts has been omitted from the plot. Also note that while $m^6A$ has been reported to be present in the genomes of *Arabidopsis*[66] and *C. elegans*[67], it is generally found at low levels ($\leq 1\%$), and is not strongly as strongly correlated with open chromatin and nucleosome positioning as it is in some other eukaryotes such as *Chlamydomonas*[45], thus its utility for accessibility profiling is not altered significantly. Nevertheless, a universally applicable version of SMAC-seq that is minimally confounded by endogenous methylation status in all eukaryotes will probably require the use of different methyltransferases (once they become available as efficient recombinant enzymes), for example, ones depositing the 4mC mark, which is what the “C” sequence context shown here corresponds to. (a) *Saccharomyces cerevisiae* (complete absence of endogenous methylation); (b) *Caenorhabditis elegans* (no endogenous 5mC, small amounts of endogenous $m^6A$); (c) *Drosophila melanogaster* (no significant endogenous 5mC or $m^6A$ methylation); (d) *Arabidopsis thaliana* (endogenous 5mC in CpG, CHG and CHH contexts, small amounts of endogenous $m^6A$); (e) *Homo sapiens* (endogenous 5mC in CpG contexts, small amounts of endogenous 5mC in CHG and CHH contexts); (f) *Mus musculus* (endogenous 5mC in CpG contexts, small amounts of endogenous 5mC in CHG and CHH contexts).

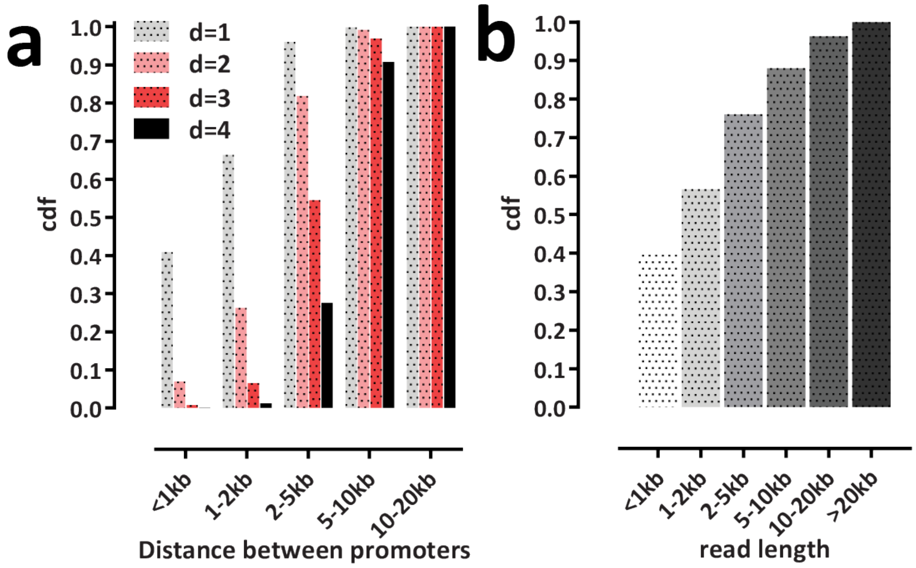


**Supplementary Figure 2: Distance between regulatory elements (i.e. promoters in the case of *S. cerevisiae*) and distribution of obtained nanopore read lengths**. (a) Distance between annotated promoters in *Saccharomyces cerevisiae*; (b) Distribution of nanopore read lengths (in the "Sample 1" experiment).

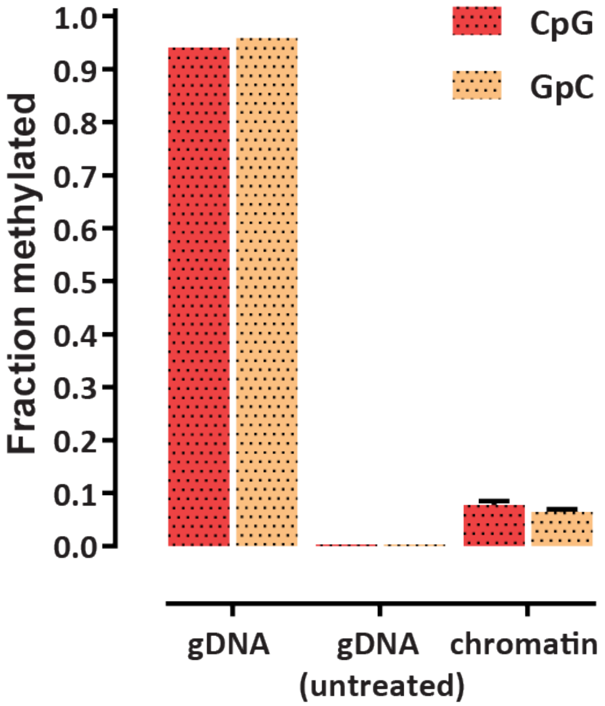


**Supplementary Figure 3: Global methylation levels in yeast dSMF experiments and in positive and negative controls (measured by bisulfite sequencing)**. DNA from SMAC-seq experiments was subjected to Illumina bisulfite sequencing using the PBAT protocol. In parallel, naked genomic DNA (gDNA) was either treated with all three enzymes under the same conditions as in the SMAC-seq protocol or it was left untreated. These samples were also subjected to Illumina PBAT bisulfite sequencing. Shown are the global methylation levels for each sample.

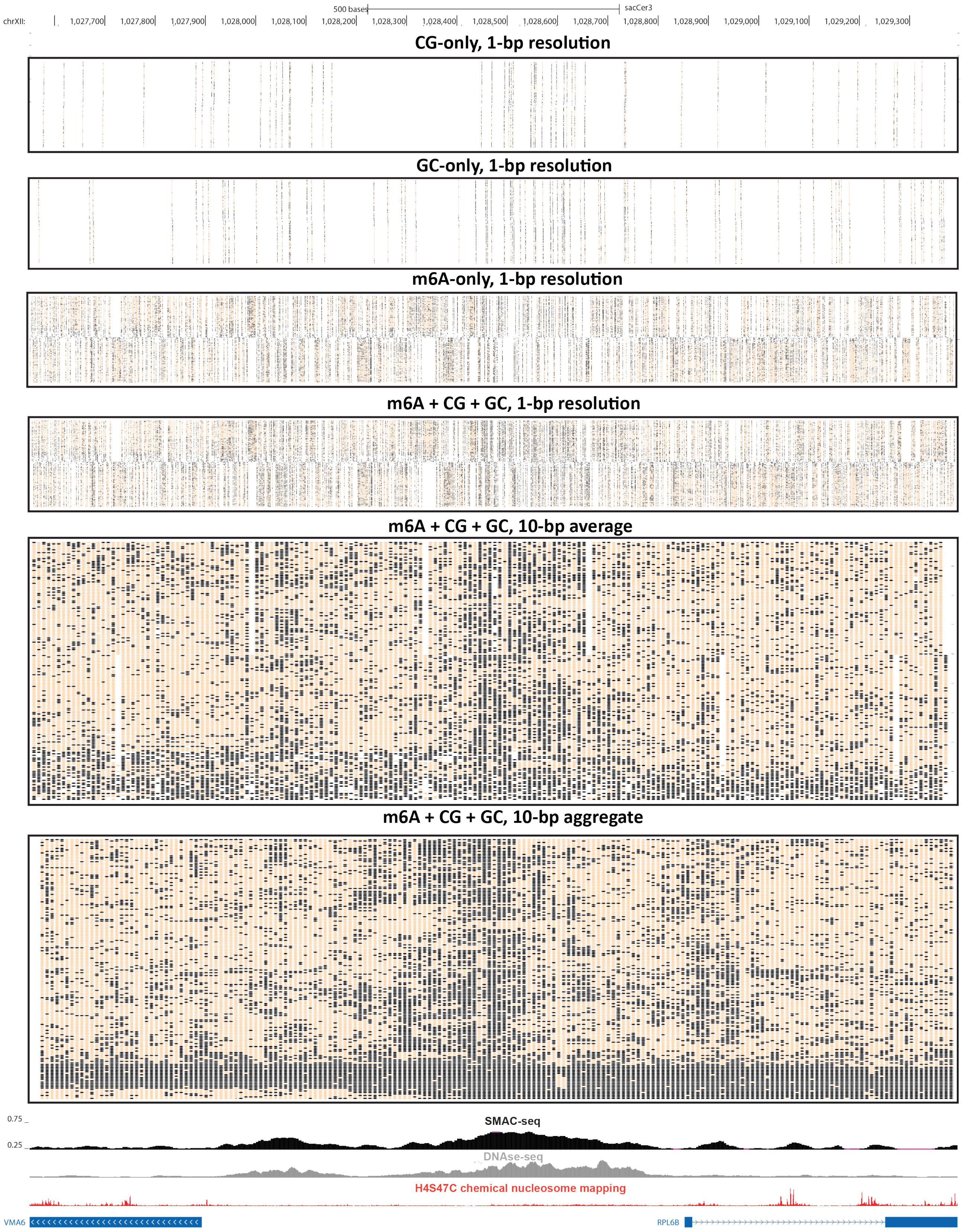


**Supplementary Figure 4: Impact of the addition of $m^6A$ on assay resolution**. Shown is the raw unfiltered nanopore read coverage around a divergent promoter region on chrXII, considering only CG, only GC, only $m^6A$, and all bases at 1-bp resolution as well as all bases at averaged and aggregated 10-bp resolution.

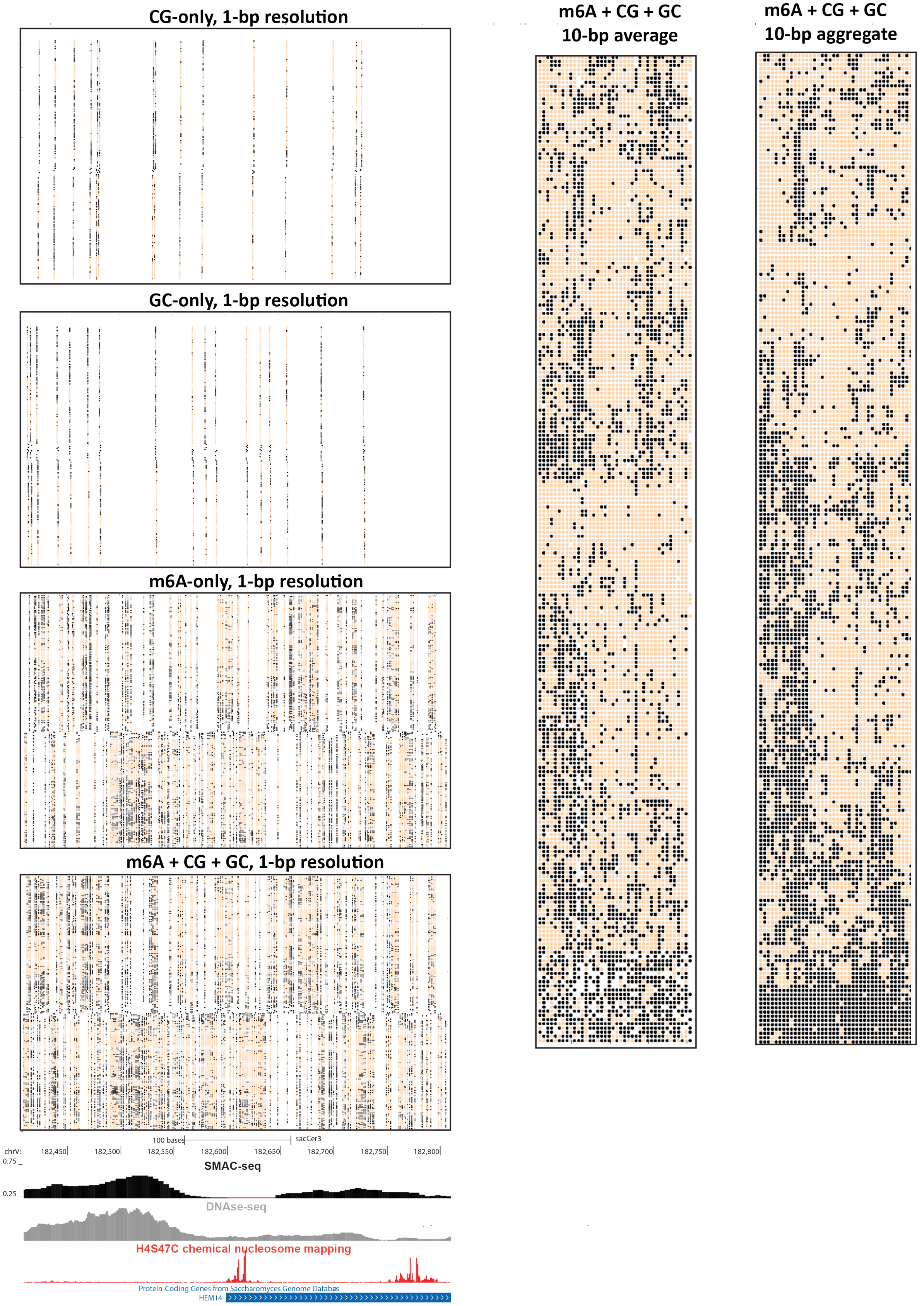


**Supplementary Figure 5: Impact of the addition of m$^6$A on assay resolution**. Shown is the raw unfiltered nanopore read coverage around a strongly positioned +1 nucleosome, considering only CG, only GC, only m$^6$A, and all bases at 1-bp resolution as well as all bases at averaged and aggregated 10-bp resolution.

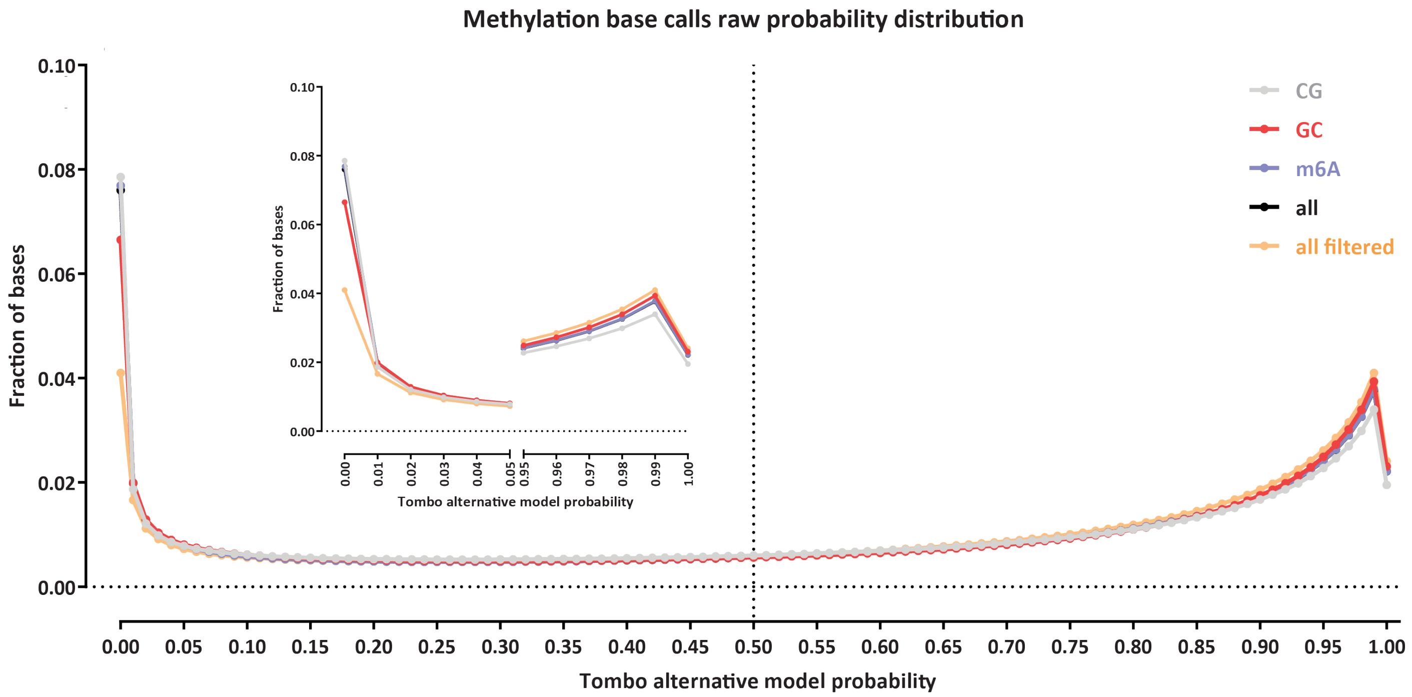


**Supplementary Figure 6: Distribution of methylation call probabilities**. Shown is the distribution of Tombo "alternative model" probabilities for all positions, and each of the three sequence contexts, as well as the distribution after filtering potential poor quality/non-chromatinized reads (see further below for more details).

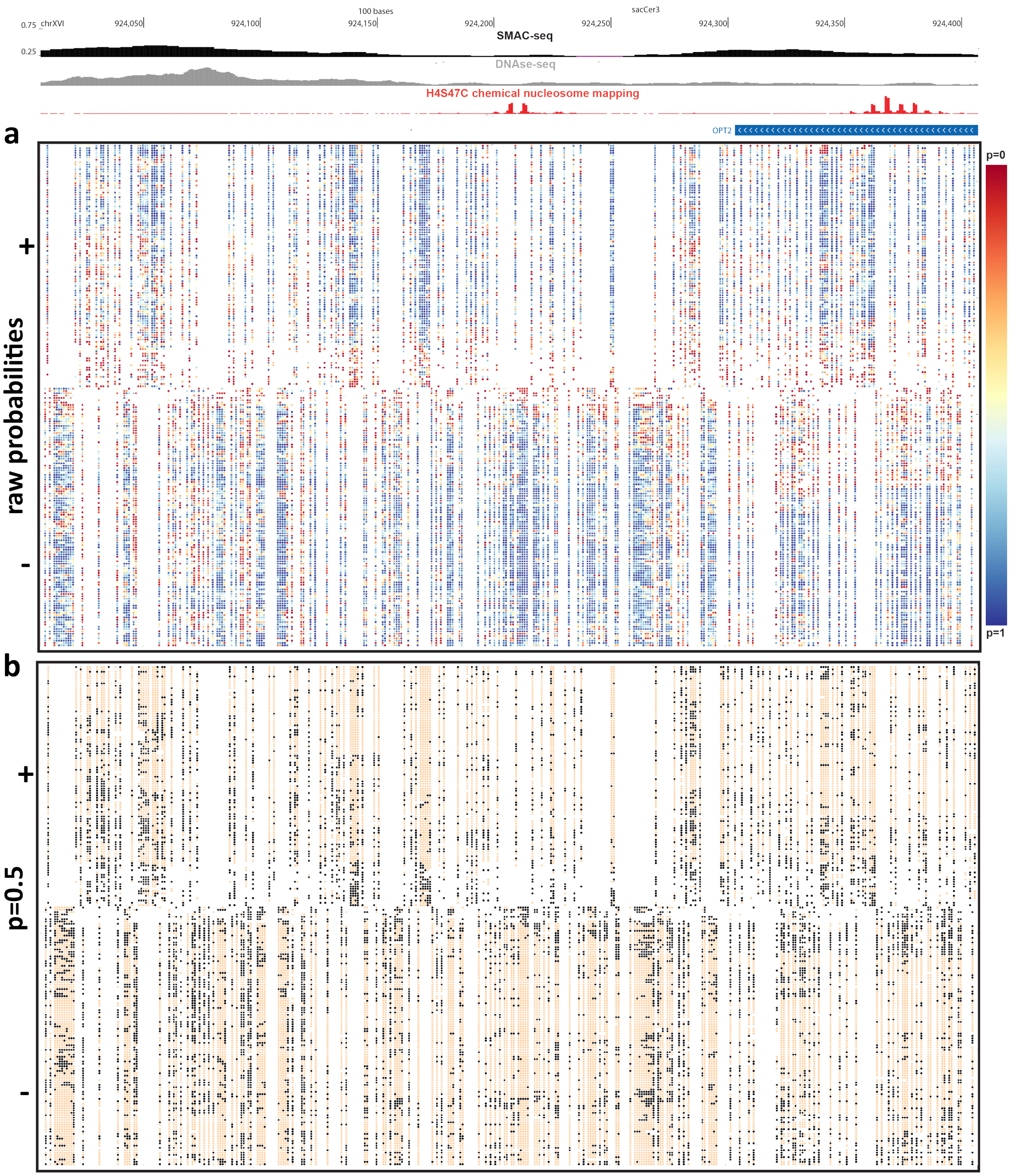

100 bases
sacCer3
chrXVI
924,050
924,100
924,150
924,200
924,250
924,300
924,350
924,400
0.75
0.25
SMAC-seq
DNAse-seq
H4S47C chemical nucleosome mapping
OPT2
a
raw probabilities
+
-
p=0
p=1
b
p=0.5
+
-

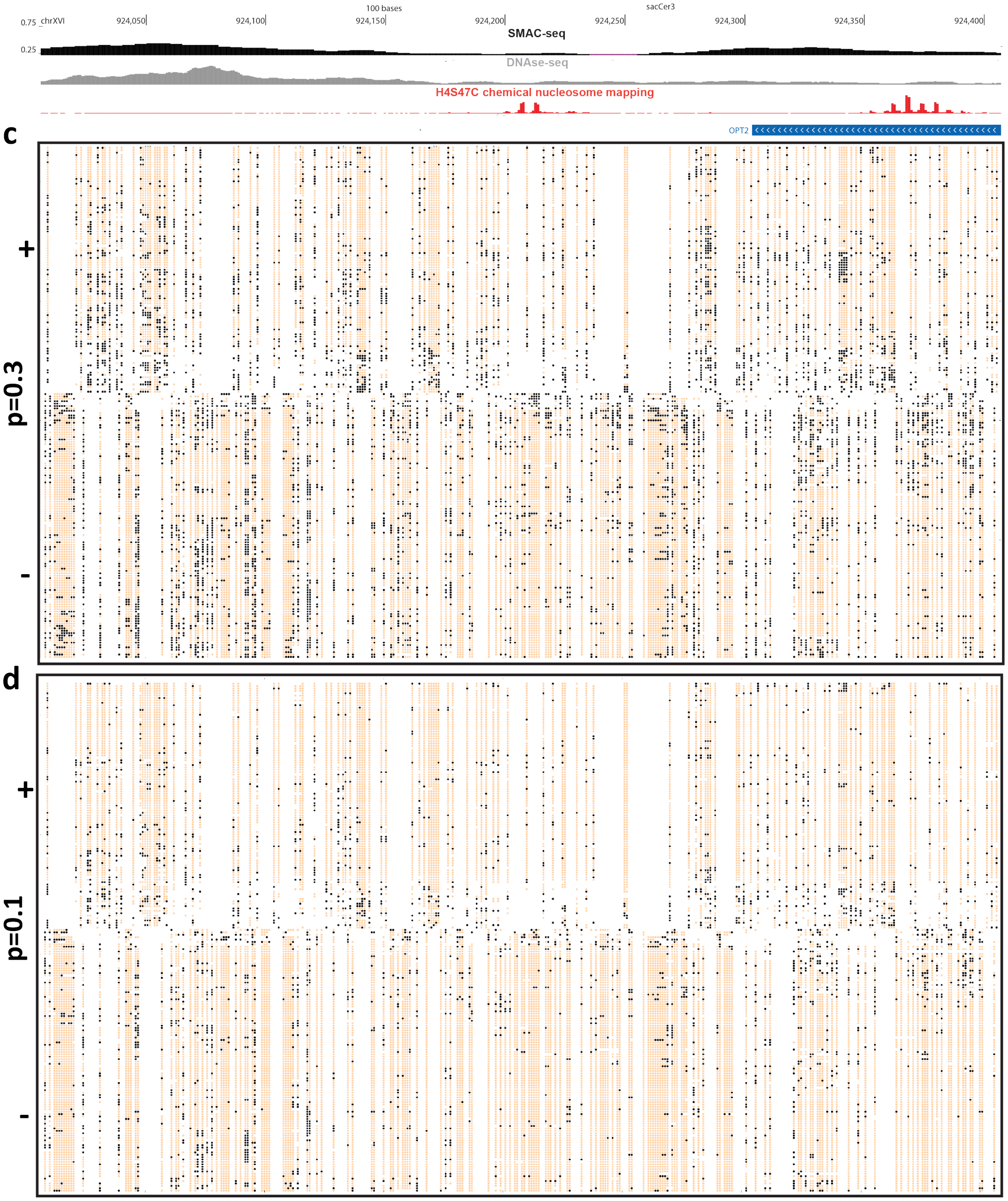


**Supplementary Figure 7: Transformation of raw methylation probabilities into binary methylation calls**. Shown is raw unfiltered SMAC-seq single-molecule data over a strongly positioned nucleosome on chrXVI (1-bp resolution). (a) raw Tombo alternative model methylation probabilities; (b) $p < 0.5$ thresholding; (d) $p < 0.3$ thresholding; (d) $p < 0.1$ thresholding.

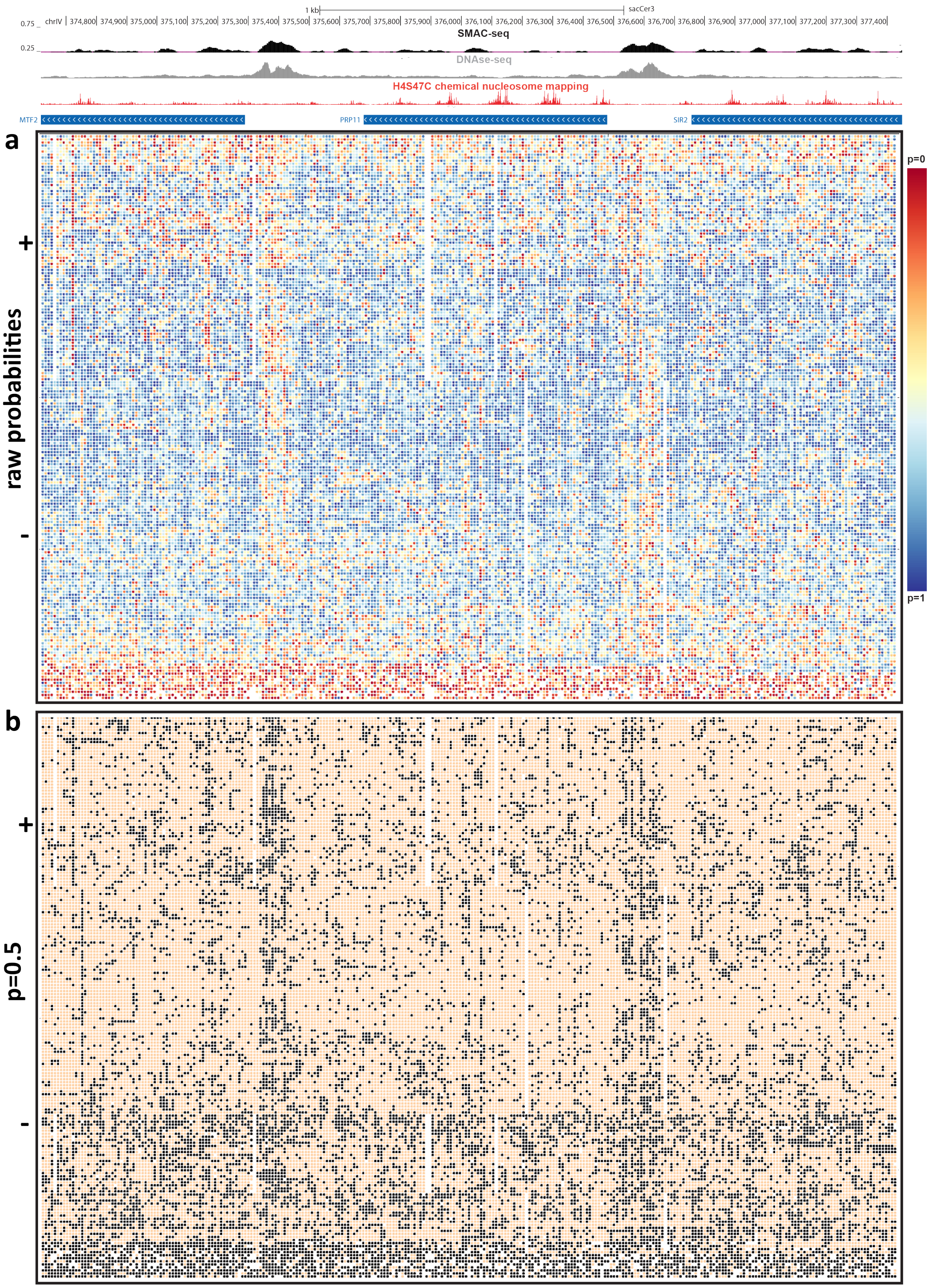
1 kb
sacCer3
chrIV
374,800 374,900 375,000 375,100 375,200 375,300 375,400 375,500 375,600 375,700 375,800 375,900 376,000 376,100 376,200 376,300 376,400 376,500 376,600 376,700 376,800 376,900 377,000 377,100 377,200 377,300 377,400
0.75
0.25
SMAC-seq
DNAse-seq
H4S47C chemical nucleosome mapping
MTF2
PRP11
SIR2
a
+
raw probabilities
-
p=0
p=1
b
+
p=0.5
-

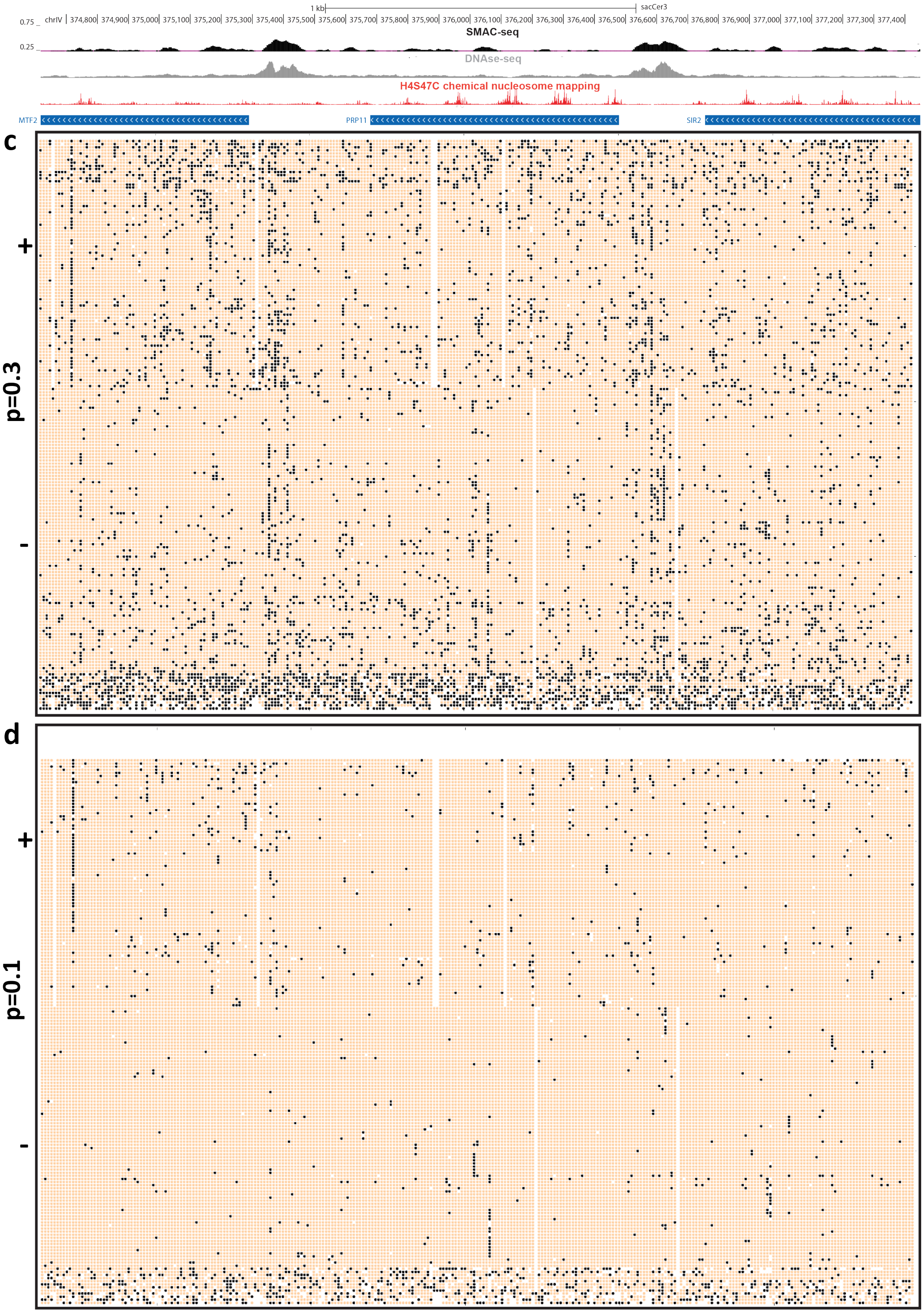


**Supplementary Figure 8: Transformation of raw methylation probabilities into binary methylation calls**. Shown is raw unfiltered SMAC-seq single-molecule data over a ∼2.8kb locus on chrIV (10-bp average for all panels). (a) raw Tombo alternative model methylation probabilities; (b) $p < 0.5$ thresholding; (d) $p < 0.3$ thresholding; (d) $p < 0.1$ thresholding.

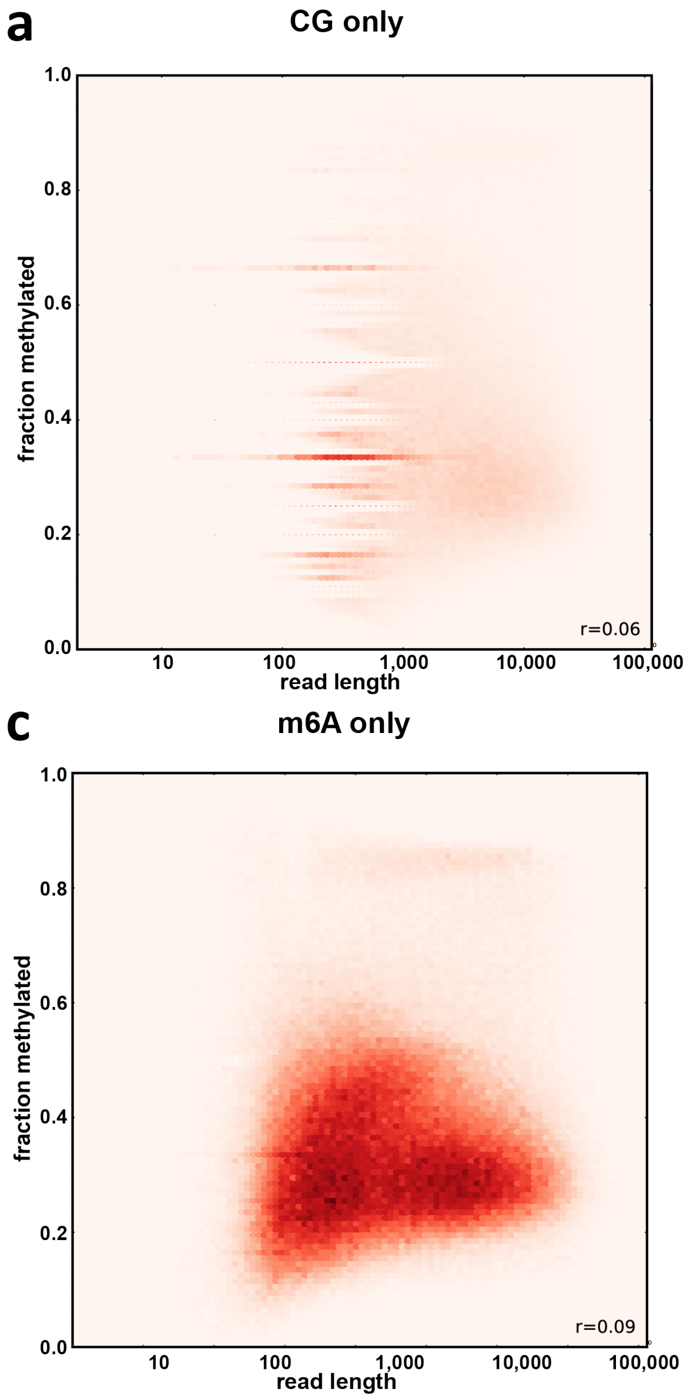


**Supplementary Figure 9: Absence of strong correlation between nanopore sequencing read length and methylation status**. Shown is the fraction of bases within each read that is scored as methylated. (a) CG positions only. (b) GC positions only. (c) $m^6A$ positions only. (d) All positions.

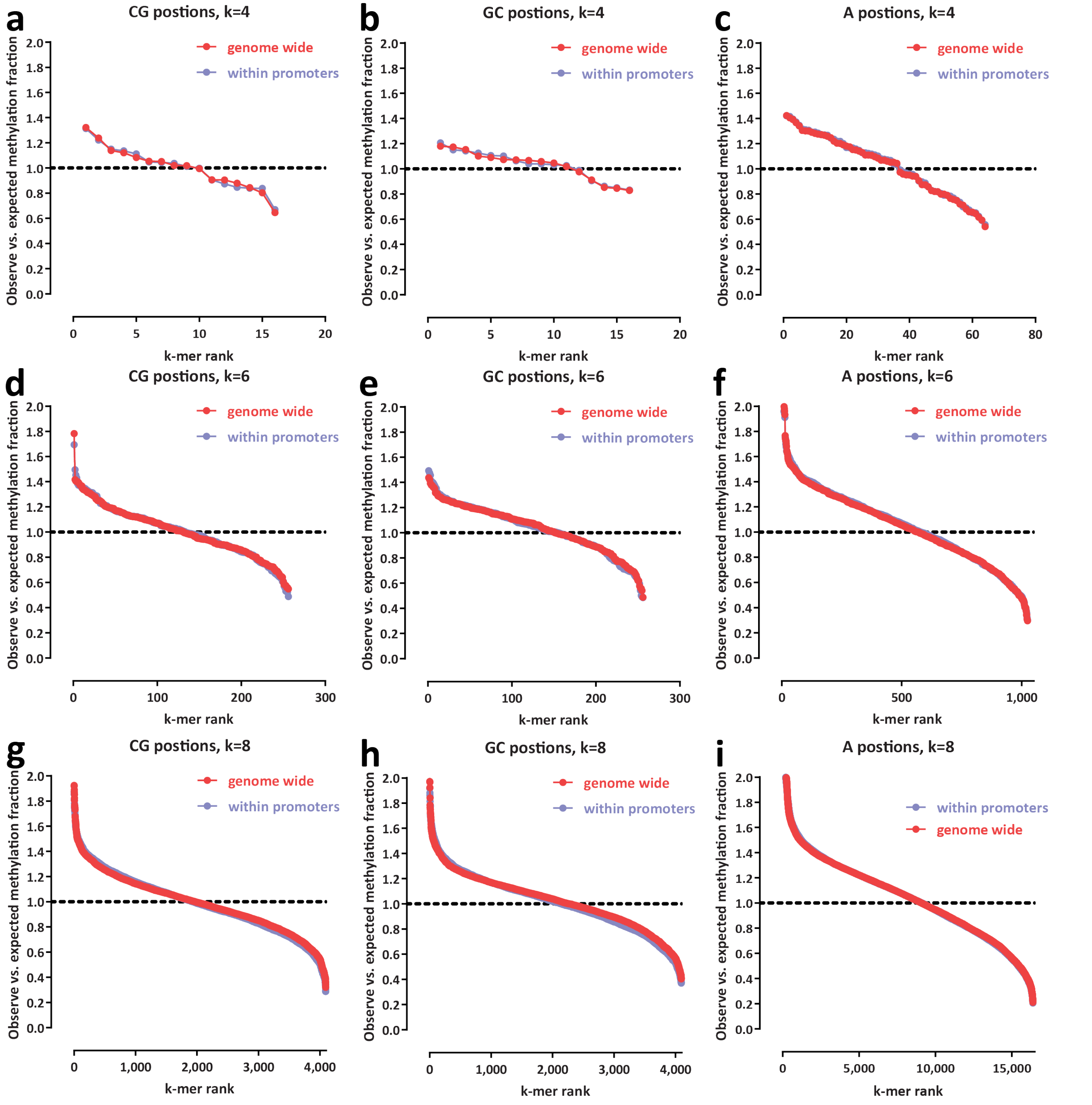


**Supplementary Figure 10: Examination of enzymatic/methylated base calling bias**. Shown is the ratio of observed versus expected fraction of methylated bases for each sequence context of size $k$, calculated as follows:

$$f_{\text{obs/exp},k} = \frac{k_m/k_u}{\sum_k k_m / \sum_k k_u}$$

where $k_m$ refers to the number of bases called as methylated across all reads and $k_m$ refers to the number of bases called as unmethylated . (a) CG positions only, $k = 4$. (b) GC positions only, $k = 4$. (c) A positions only, $k = 4$. (d) CG positions only, $k = 6$. (e) GC positions only, $k = 6$. (f) A positions only, $k = 6$. (g) CG positions only, $k = 8$. (h) GC positions only, $k = 8$. (i) A positions only, $k = 8$.

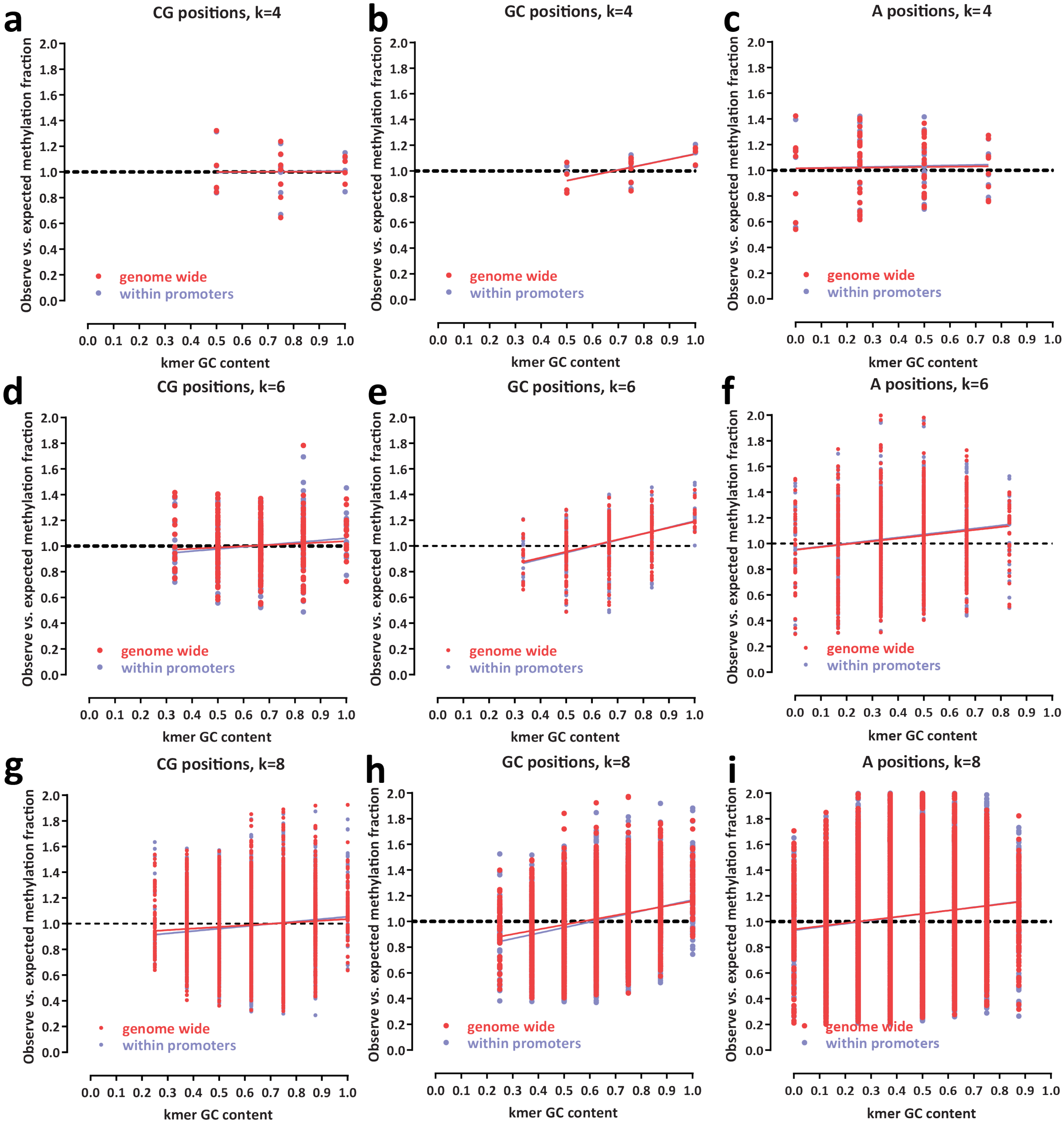


**Supplementary Figure 11: Relationship between local GC content and enzymatic/methylated base calling bias**. Shown is the ratio of observed versus expected fraction of methylated bases for each sequence context of size $k$, calculated as follows:

$$f_{\text{obs/exp},k} = \frac{k_m/k_u}{\sum_k k_m / \sum_k k_u}$$

where $k_m$ refers to the number of bases called as methylated across all reads and $k_m$ refers to the number of bases called as unmethylated . (a) CG positions only, $k = 4$. (b) GC positions only, $k = 4$. (c) A positions only, $k = 4$. (d) CG positions only, $k = 6$. (e) GC positions only, $k = 6$. (f) A positions only, $k = 6$. (g) CG positions only, $k = 8$. (h) GC positions only, $k = 8$. (i) A positions only, $k = 8$.

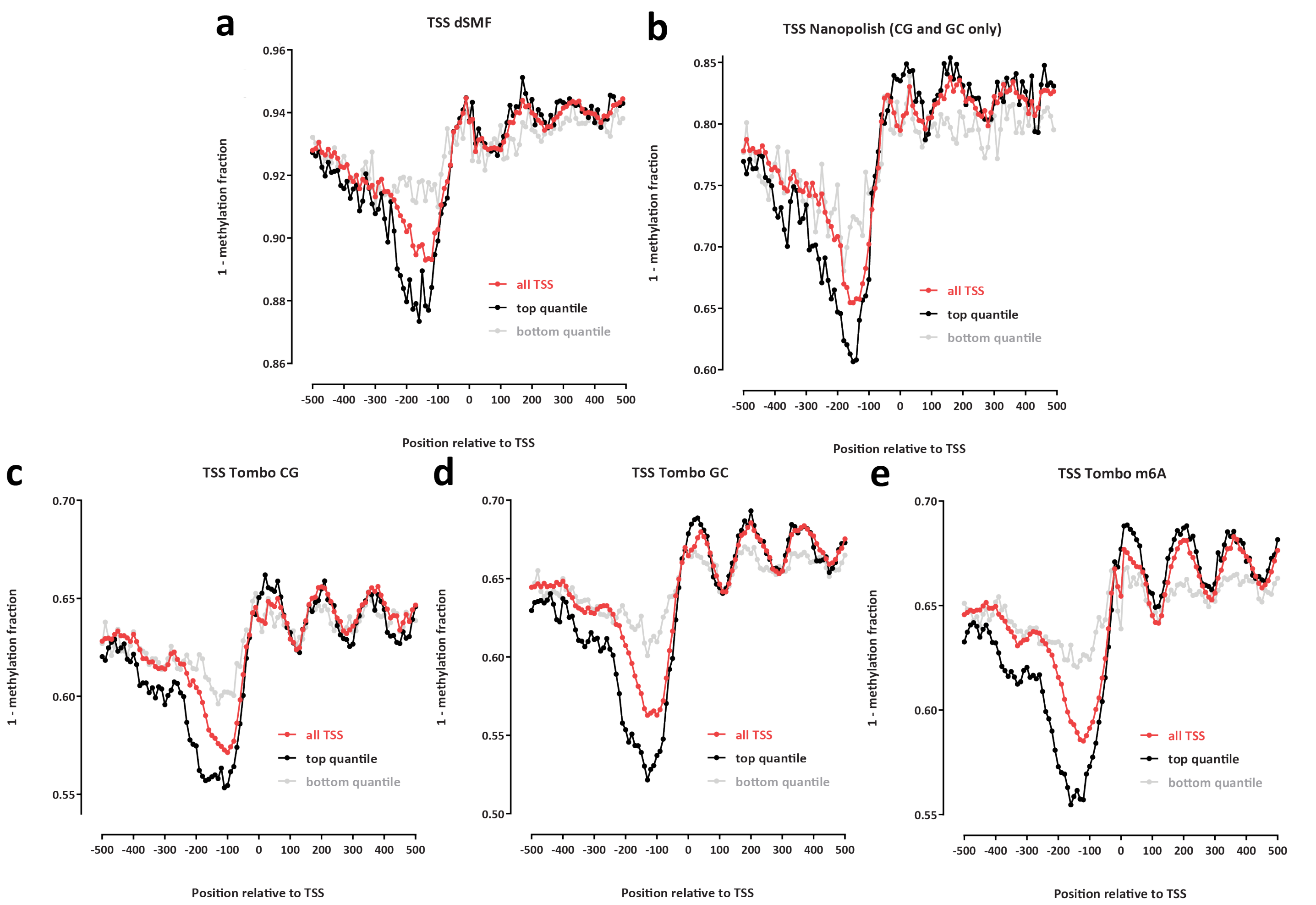


**Supplementary Figure 12: Comparison of dSMF results and different approaches to methylation-aware base calling on SMAC-seq data**. Shown is the inverse of the methylated fraction of nucleotides around TSSs of all, highly express (top quantile) and low expression-level (bottom quantiles) yeast genes (unfiltered "Sample 1" dataset). Note that the different panels are not drawn to the same scale. (a) dSMF; (b) SMAC-seq data using Nanopolish methylation base-calling on CG and GC nucleotides; (c) SMAC-seq data using Tombo methylation base-calling, CG positions only; (d) SMAC-seq data using Tombo methylation base-calling, GC positions only; (e) SMAC-seq data using Tombo methylation base-calling, m6A positions only.

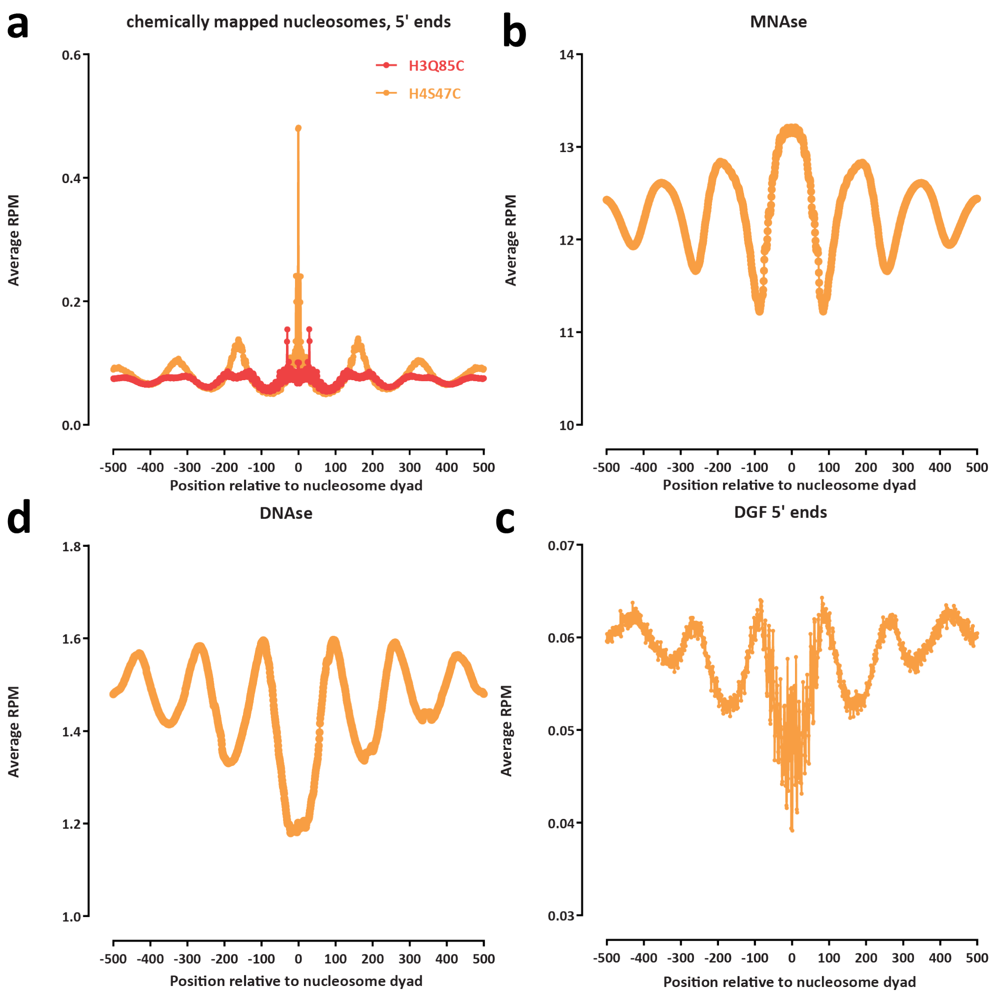


**Supplementary Figure 13: Correlation of SMAC-seq (Figure 1e) with other measures of chromatin structure around the dyad centers of positioned nucleosomes in the *S. cerevisiae* genome**. (a) H4S47C and H3Q85C nucleosome chemical mapping; (b) MNAseq-seq; (c) DNAse-seq; (d) Digital Genomic Footprinting (DGF, 5′ ends of deeply sequenced DNAseq-seq data).

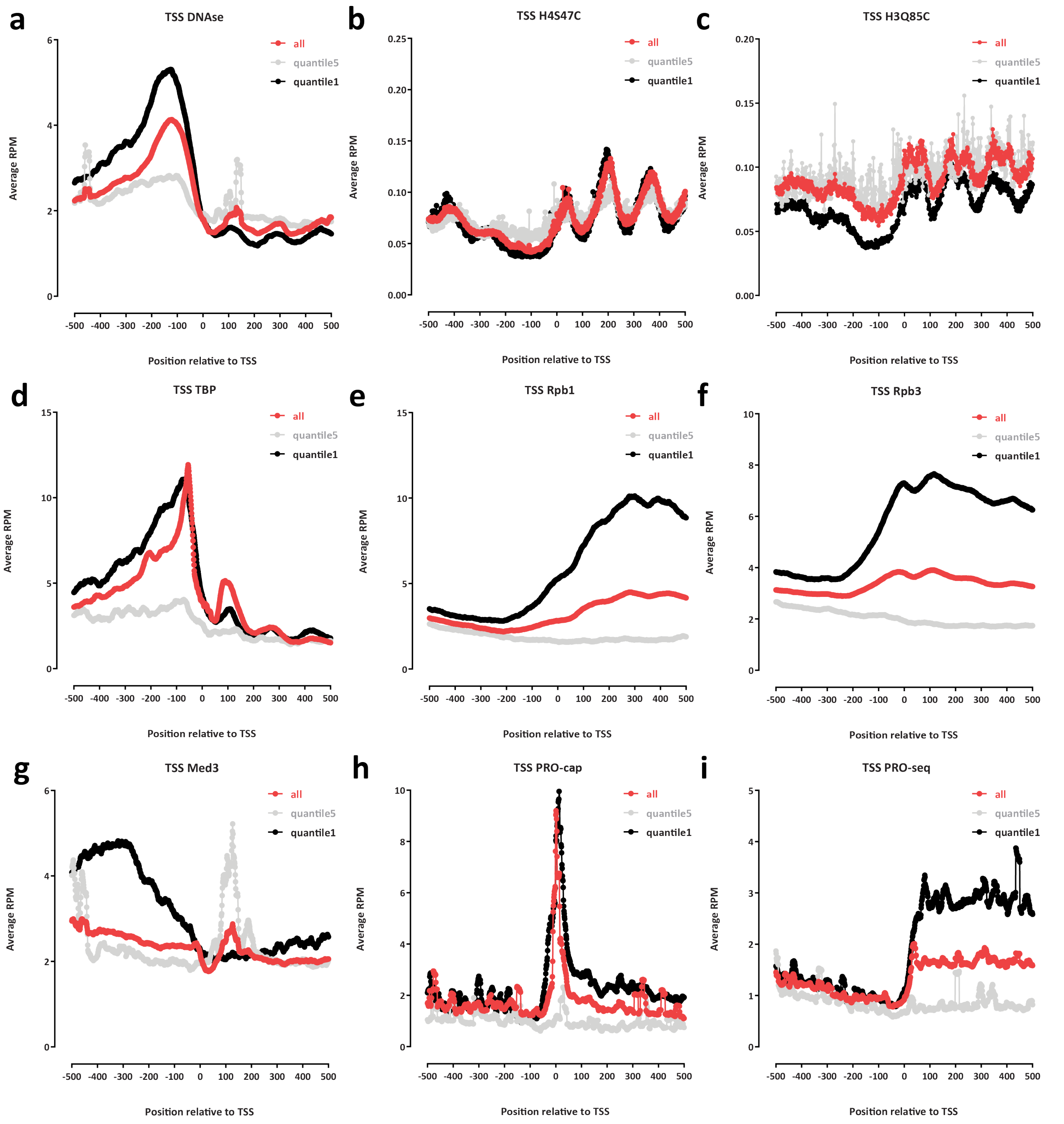


**Supplementary Figure 14: Correlation of SMAC-seq (Figure 1f and g) with other types of functional genomic measurements of chromatin structure, protein occupancy and transcriptional activity around TSSs**. Shown is average coverage over all *S. cerevisiae* genes, for the most highly expressed 20% of genes ("quantile1"), and for the bottom 20% of genes ("quantile5"). (a) DNAse-seq; (b) H4S47C nucleosome chemical mapping; (c) H3Q85C nucleosome chemical mapping; (d) TBP ChIP-seq; (e) Rpb1 ChIP-seq; (f) Rpb3 ChIP-seq; (g) Med3 ChIP-seq; (h) PRO-cap; (i) PRO-seq.

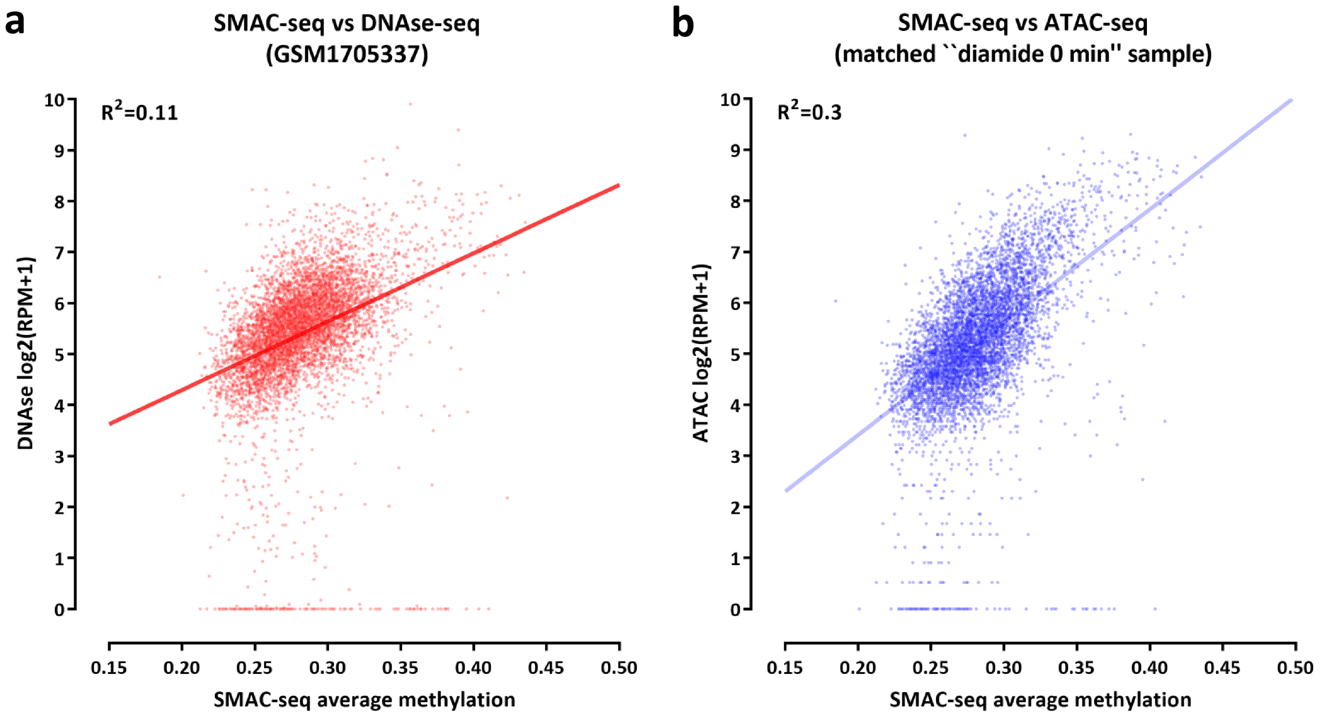


**Supplementary Figure 15: Correlation of SMAC-seq signal with ATAC-seq and DNA-seq signal over yeast promoters**. Shown is the average methylation over the TSS $\pm$ 200 bp for SMAC-seq and RPM (Reads Per Million mapped reads) values for DNAse-seq (a) and ATAC-seq (b). Note that the DNAse-seq dataset is obtained from an external study while the SMAC-seq and ATAC-seq ones are from the same sample ("diamide 0 min rep1").

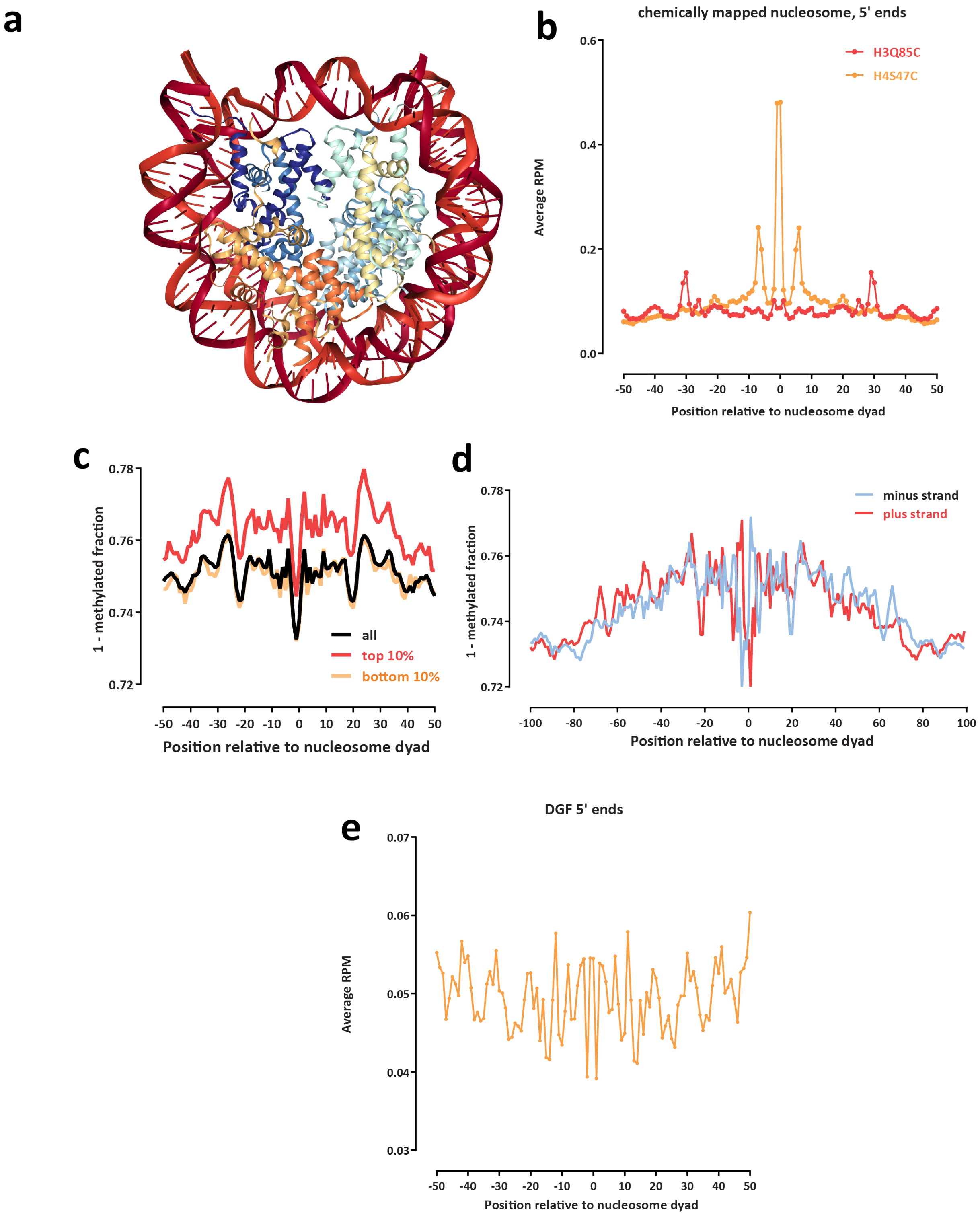


**Supplementary Figure 16: SMAC-seq data displays higher methylation propensity in more exposed parts of the nucleosomal particle**. (a) Structure of the eukaryotic nucleosome; (b) High-resolution (50-bp radius) view of chemical nucleosome mapping data relative to nucleosome dyads; (c) High-resolution (50-bp radius) view of SMAC-seq data relative to nucleosome dyads; (d) Strand-specific (100-bp radius) view of SMAC-seq data relative to nucleosome dyads; (e) High-resolution (50-bp radius) view of DGF cleavage profiles relative to nucleosome dyads.

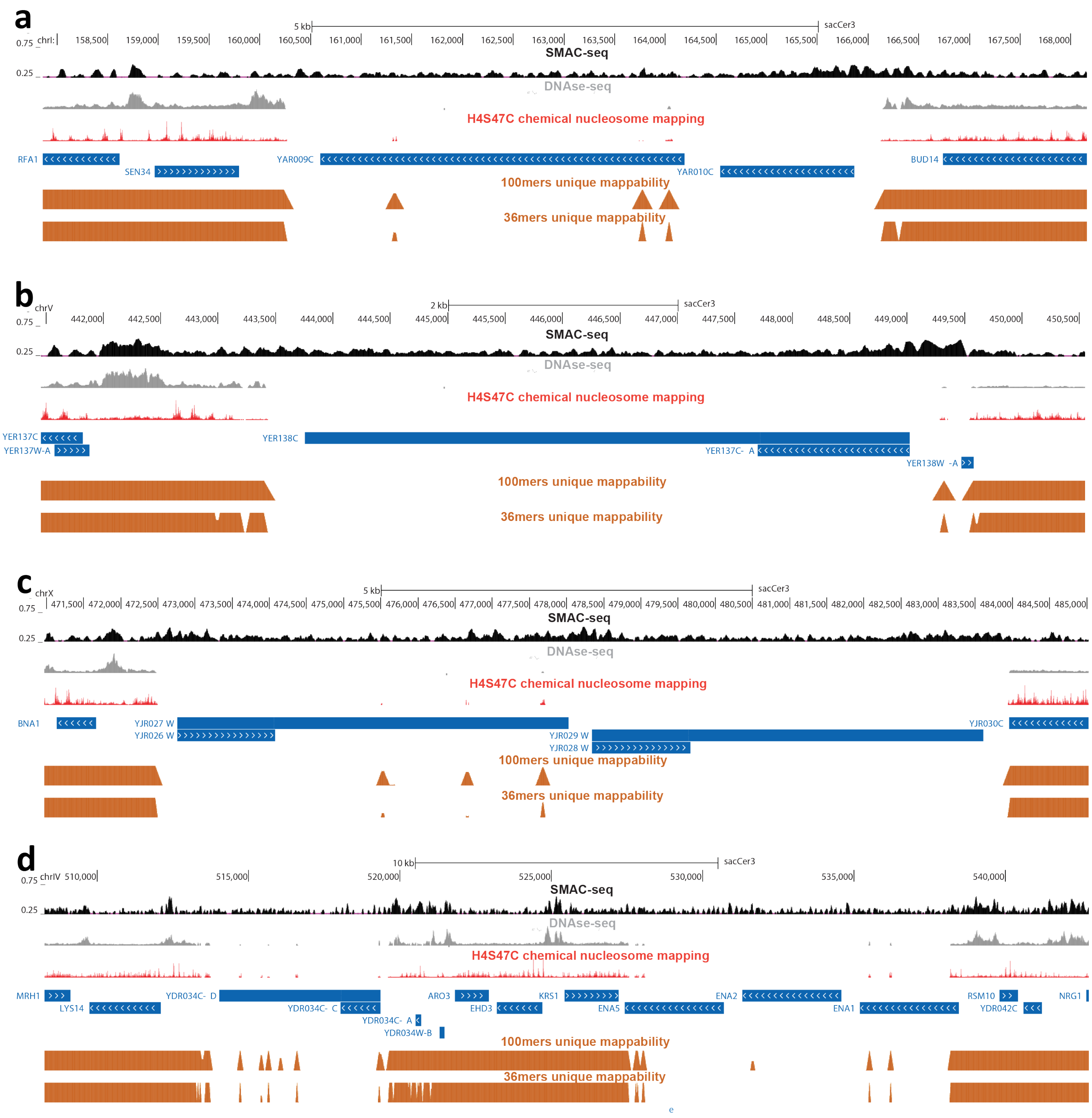


**Supplementary Figure 17: SMAC-seq provides coverage of areas of the genome that cannot be uniquely mapped using short reads.** Four different repetitive regions are shown (a, b, c and d) together with short read coverage for DNAse-seq and chemical nucleosome mapping data and unique mappability tracks for 36mers and 100mers.

**a**

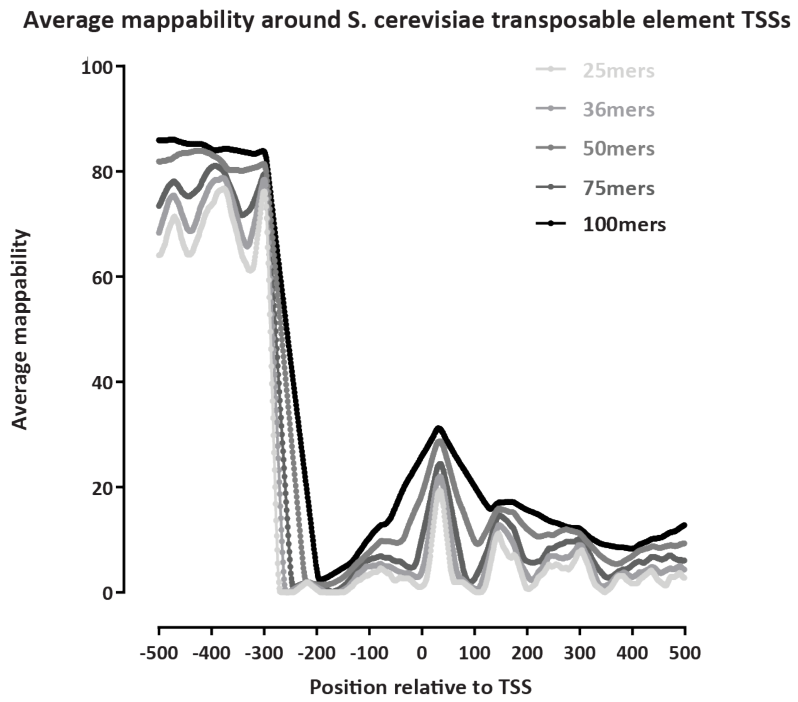


**b**

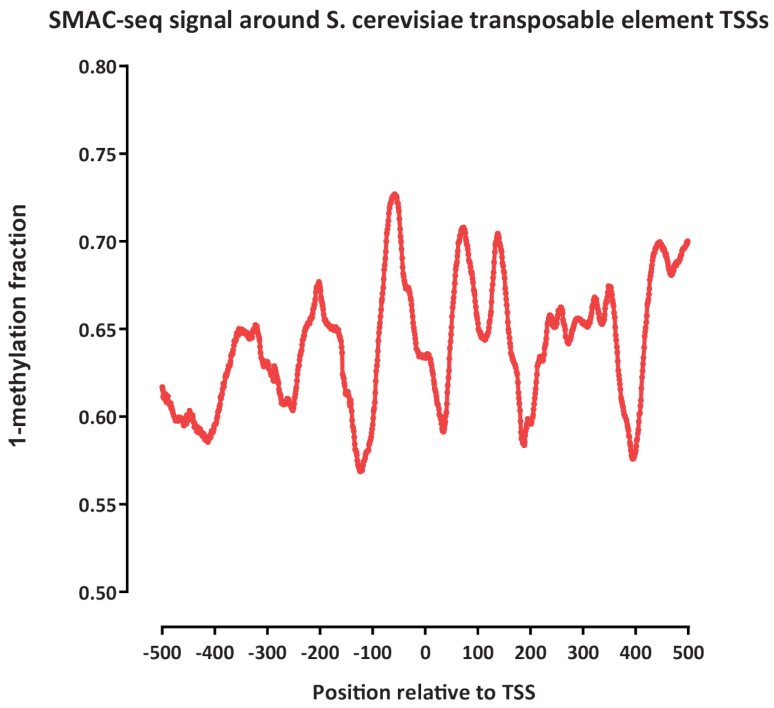


**Supplementary Figure 18: SMAC-seq provides coverage of areas of the genome that cannot be uniquely mapped using short reads**. (a) Average short-read mappability around TSSs of annotated transposable elements in the *S. cerevisiae* genome; (b) SMAC-seq signal around TSSs of annotated transposable elements in the *S. cerevisiae* genome.

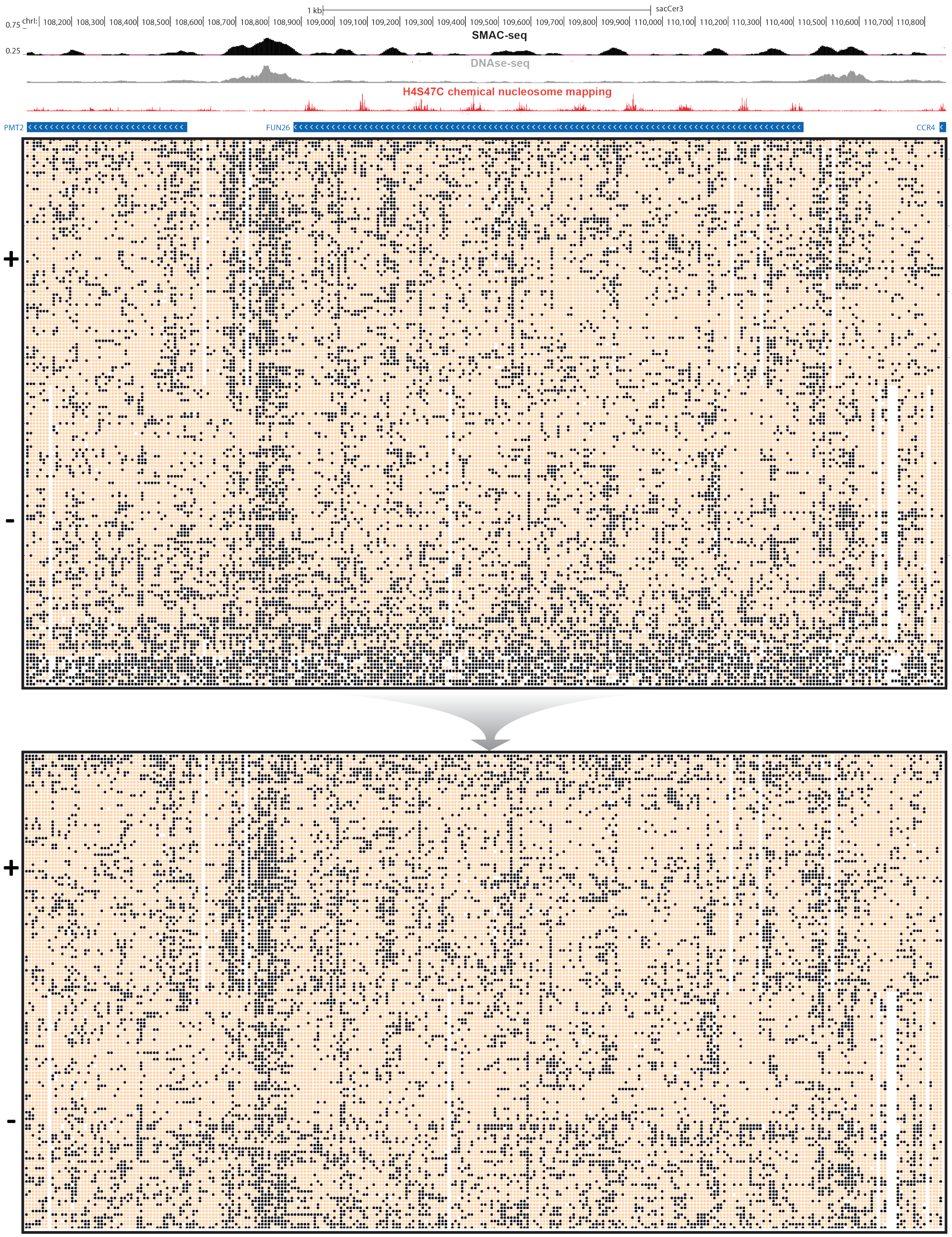


**Supplementary Figure 19: Removal of potentially artifactual high-methylation reads**. Shown is unfiltered SMAC-seq data and the same locus after removal of all reads containing a 1-kb ≥75% methylated stretch (average accessibility over 10 bp).

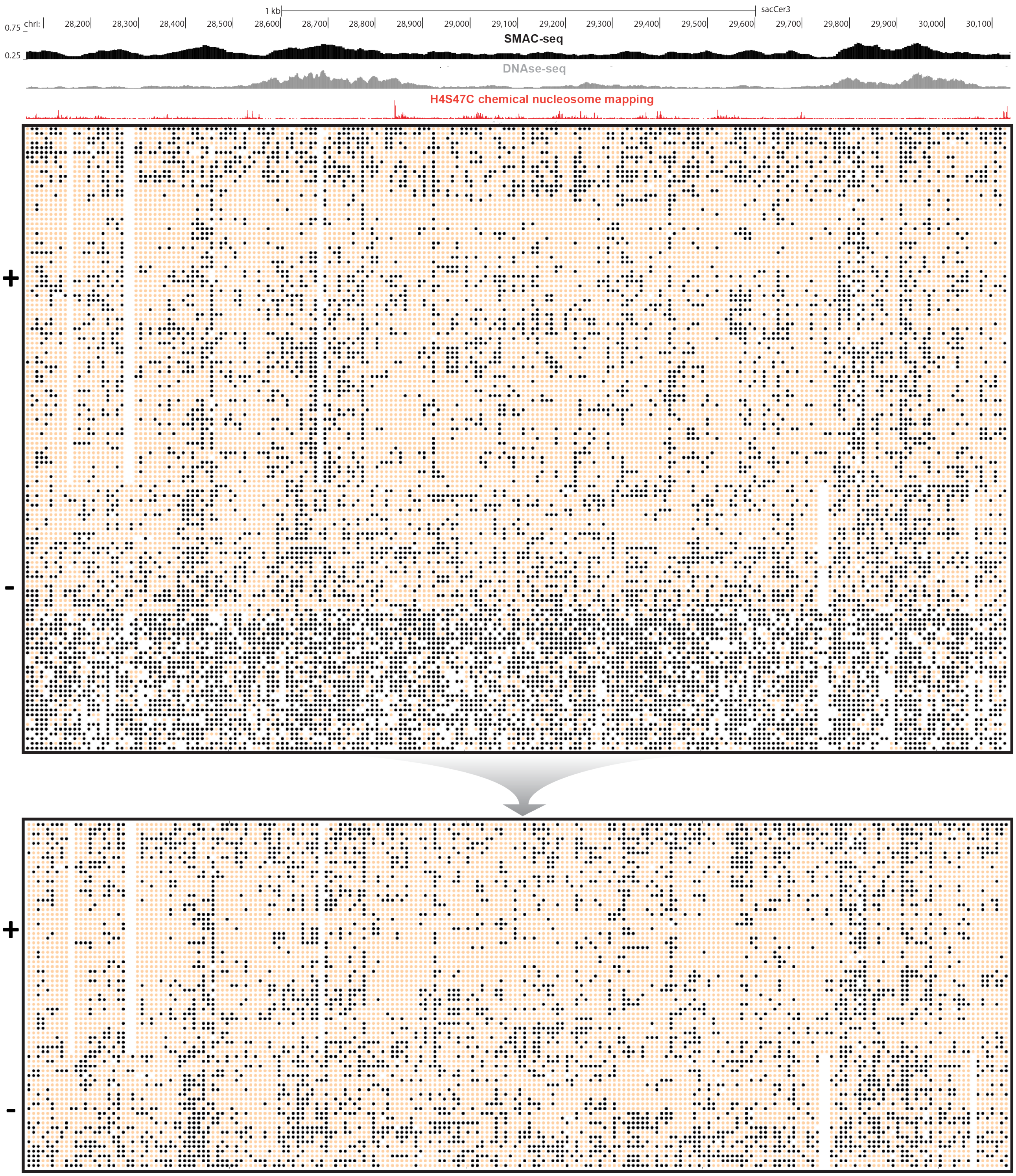


**Supplementary Figure 20: Removal of potentially artifactual high-methylation reads**. Shown is unfiltered SMAC-seq data and the same locus after removal of all reads containing a 1-kb ≥75% methylated stretch (average accessibility over 10 bp).

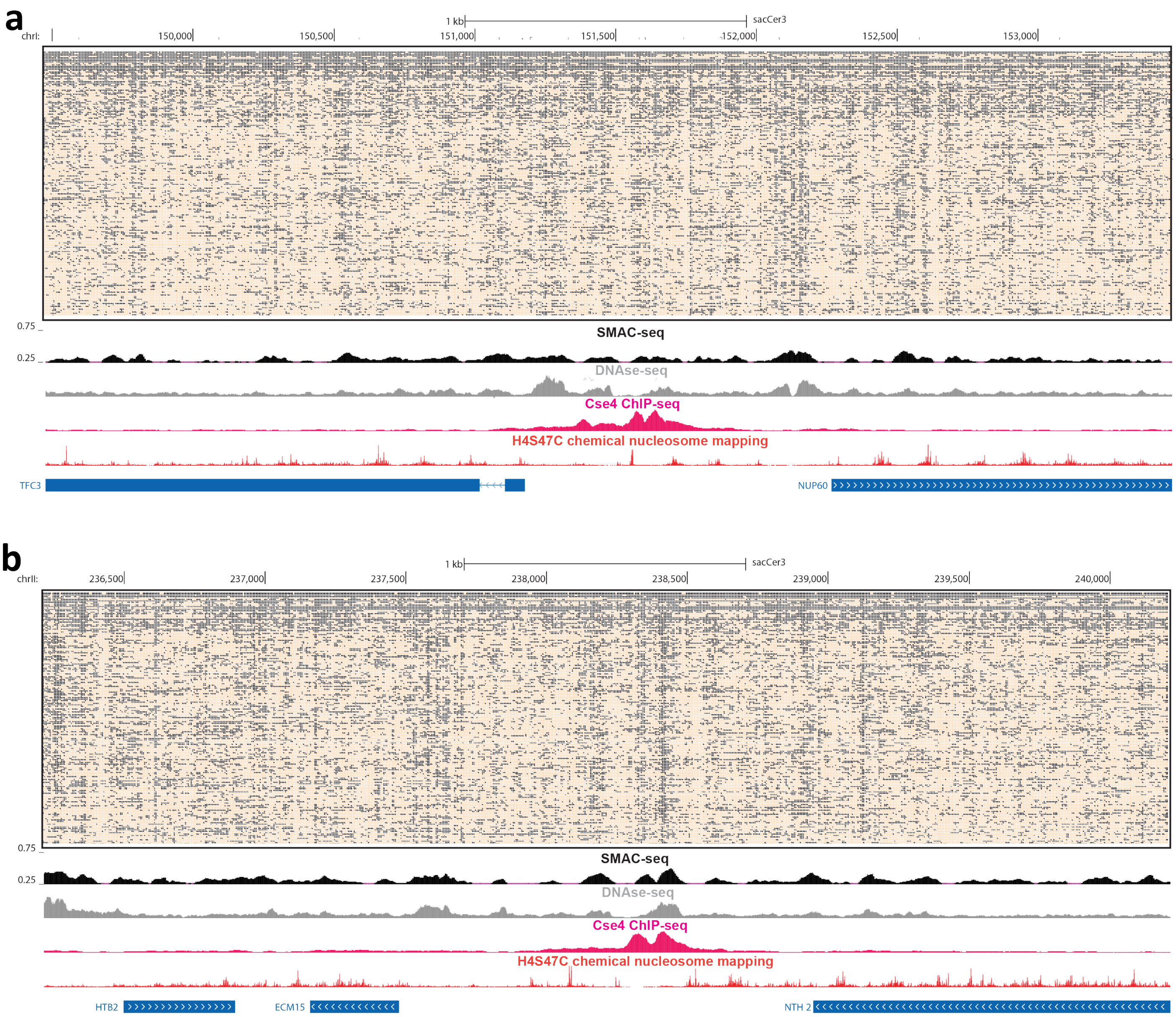


**Supplementary Figure 21: Single-molecule long-read accessibility around well positioned centromeres**. (a) Raw unfiltered nanopore reads fully spanning the 4-kilobase neighborhood of the centromere of *S. cerevisiae* chrI; (b) Raw unfiltered nanopore reads fully spanning the 4-kilobase neighborhood of the centromere of *S. cerevisiae* chrII. In both cases, accessibility is shown at aggregated 10-bp resolution (see Methods section for details) for the single-molecule display, and aggregated over sliding (every 5 bases) 50-bp windows for the genome browser SMAC-seq track.

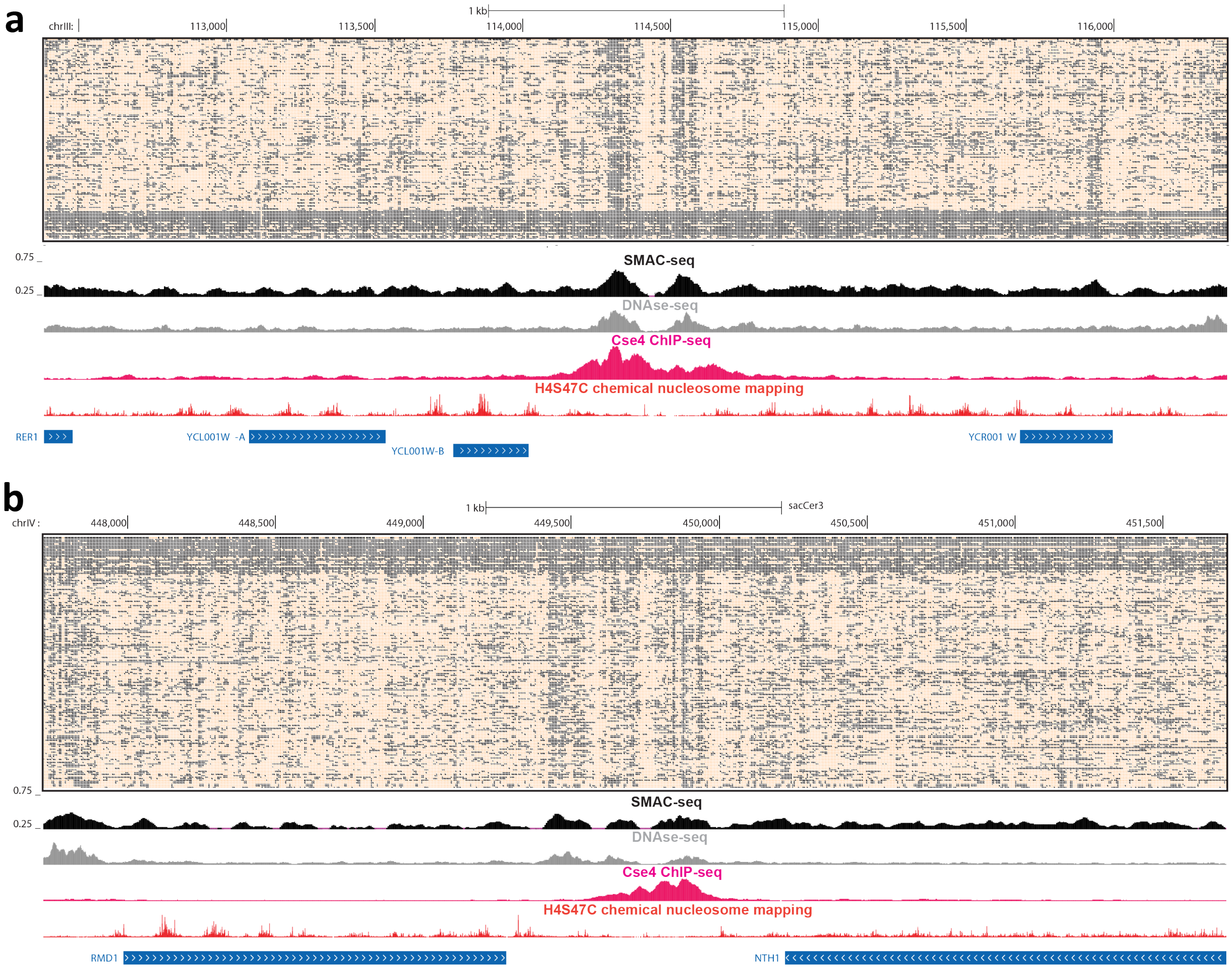


**Supplementary Figure 22: Single-molecule long-read accessibility around well positioned centromeres**. (a) Raw unfiltered nanopore reads fully spanning the 4-kilobase neighborhood of the centromere of *S. cerevisiae* chrIII; (b) Raw unfiltered nanopore reads fully spanning the 4-kilobase neighborhood of the centromere of *S. cerevisiae* chrIV. In both cases, accessibility is shown at aggregated 10-bp resolution (see Methods section for details) for the single-molecule display, and aggregated over sliding (every 5 bases) 50-bp windows for the genome browser SMAC-seq track.

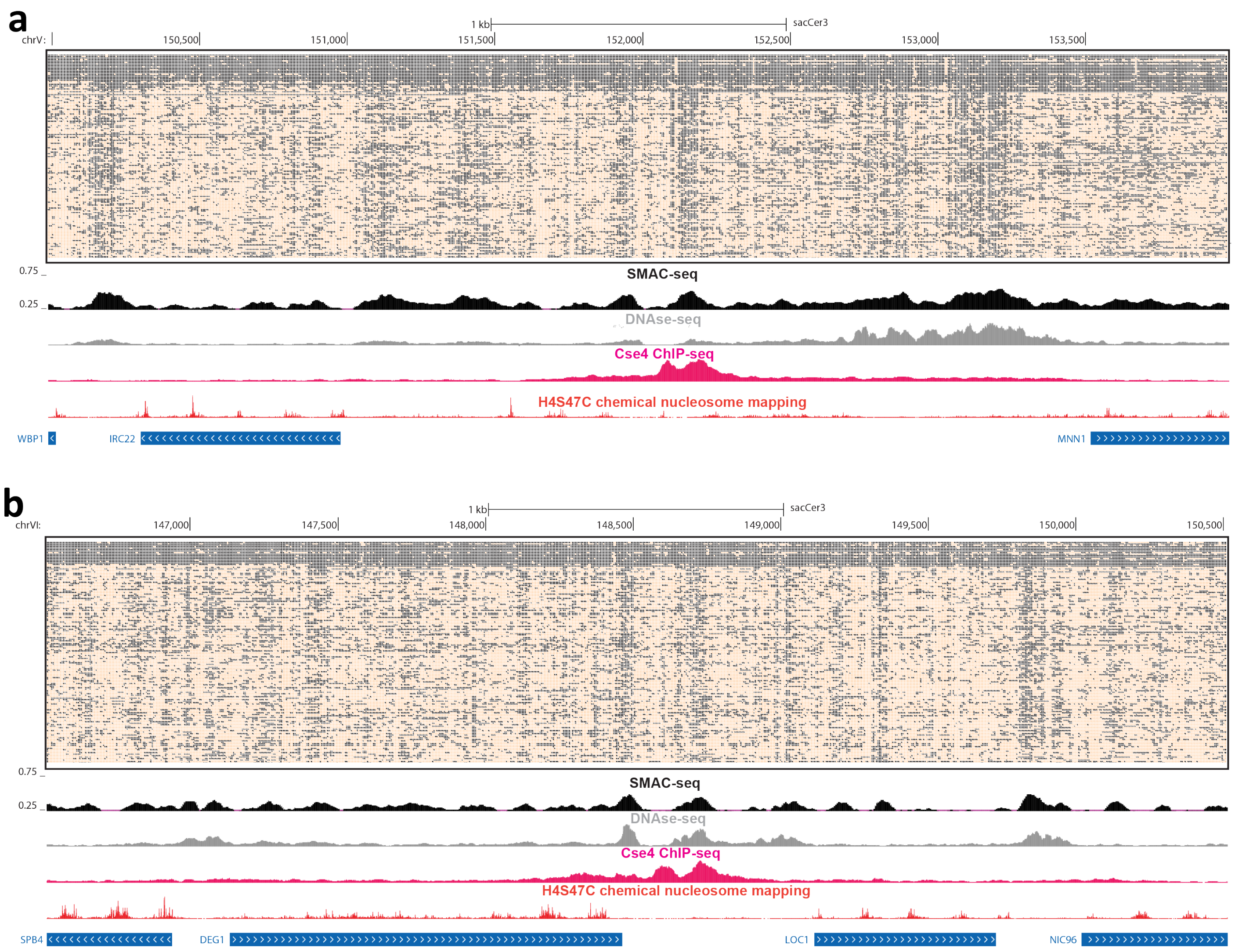


**Supplementary Figure 23: Single-molecule long-read accessibility around well positioned centromeres**. (a) Raw unfiltered nanopore reads fully spanning the 4-kilobase neighborhood of the centromere of *S. cerevisiae* chrV; (b) Raw unfiltered nanopore reads fully spanning the 4-kilobase neighborhood of the centromere of *S. cerevisiae* chrVI. In both cases, accessibility is shown at aggregated 10-bp resolution (see Methods section for details) for the single-molecule display, and aggregated over sliding (every 5 bases) 50-bp windows for the genome browser SMAC-seq track.

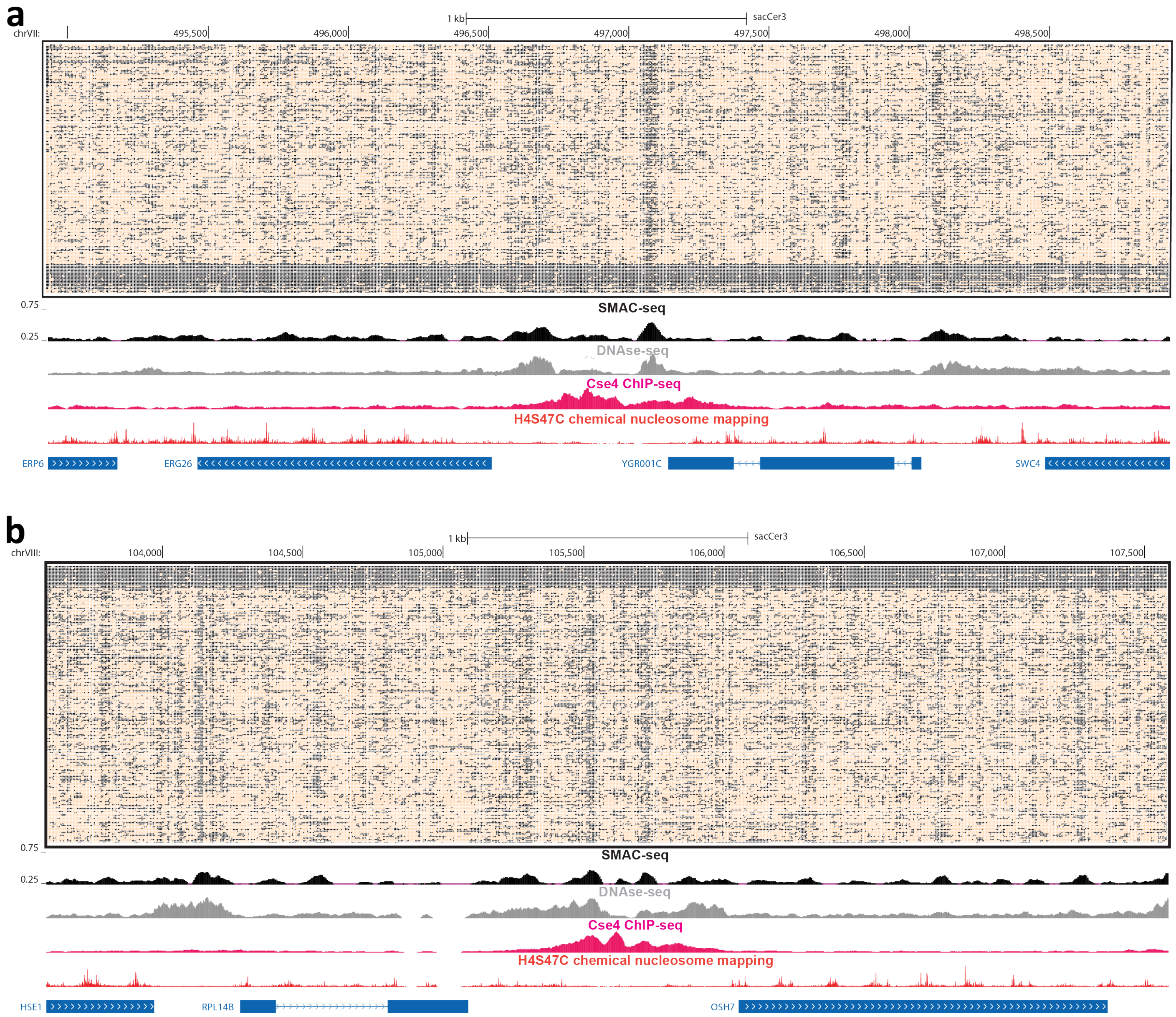


**Supplementary Figure 24: Single-molecule long-read accessibility around well positioned centromeres**. (a) Raw unfiltered nanopore reads fully spanning the 4-kilobase neighborhood of the centromere of *S. cerevisiae* chrVII; (b) Raw unfiltered nanopore reads fully spanning the 4-kilobase neighborhood of the centromere of *S. cerevisiae* chrVIII. In both cases, accessibility is shown at aggregated 10-bp resolution (see Methods section for details) for the single-molecule display, and aggregated over sliding (every 5 bases) 50-bp windows for the genome browser SMAC-seq track.

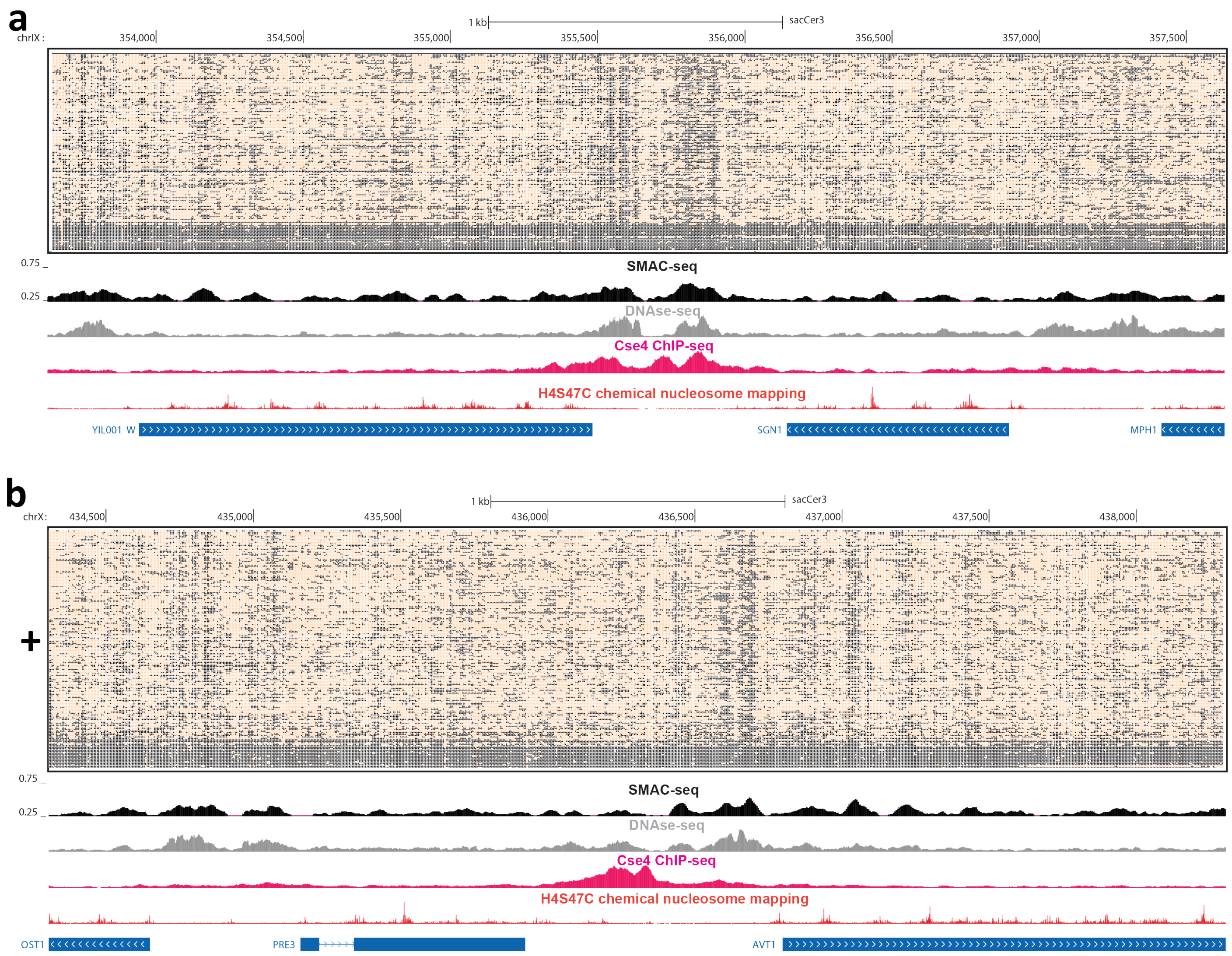


**Supplementary Figure 25: Single-molecule long-read accessibility around well positioned centromeres**. (a) Raw unfiltered nanopore reads fully spanning the 4-kilobase neighborhood of the centromere of *S. cerevisiae* chrIX; (b) Raw unfiltered nanopore reads fully spanning the 4-kilobase neighborhood of the centromere of *S. cerevisiae* chrX. In both cases, accessibility is shown at aggregated 10-bp resolution (see Methods section for details) for the single-molecule display, and aggregated over sliding (every 5 bases) 50-bp windows for the genome browser SMAC-seq track.

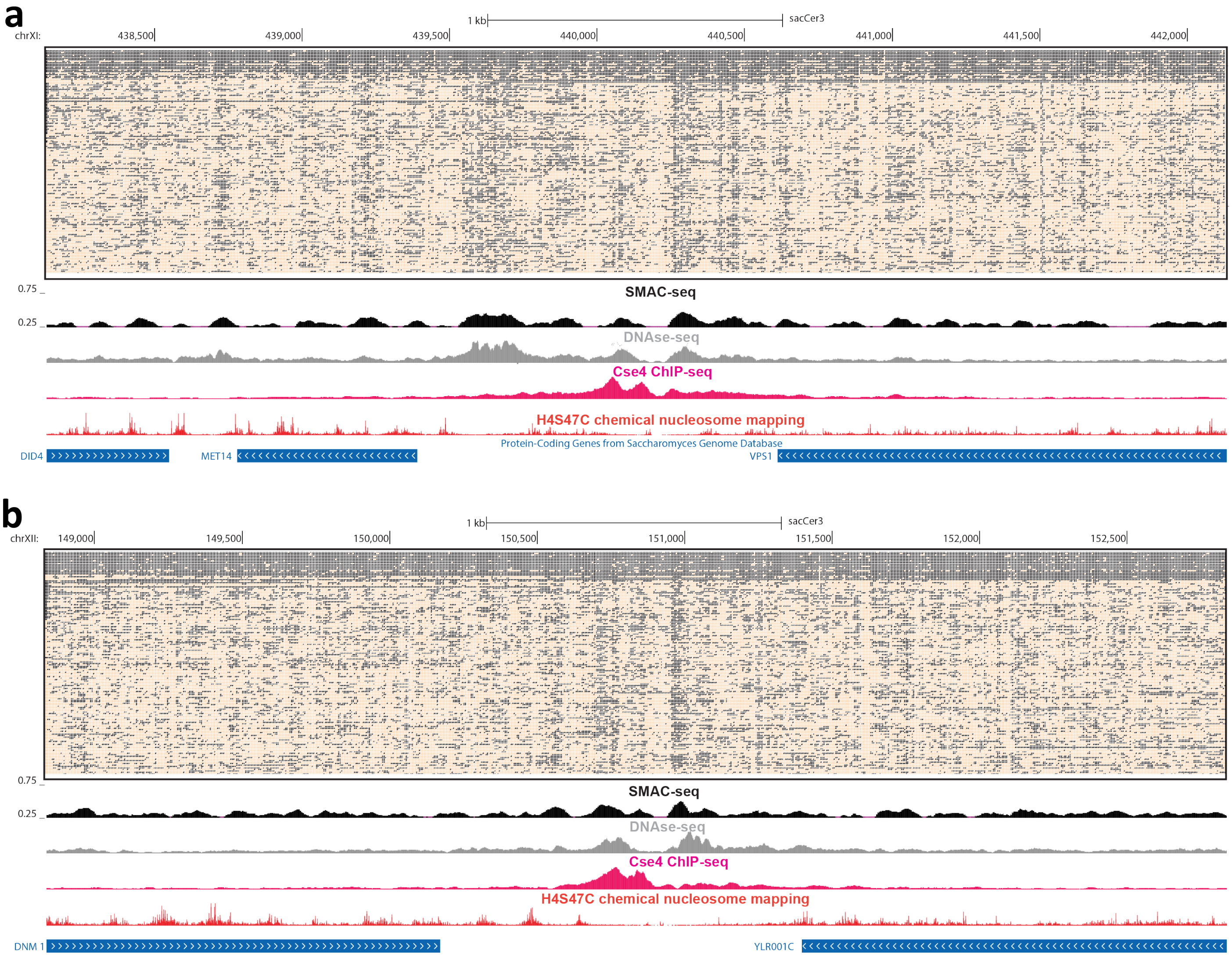


**Supplementary Figure 26: Single-molecule long-read accessibility around well positioned centromeres**. (a) Raw unfiltered nanopore reads fully spanning the 4-kilobase neighborhood of the centromere of *S. cerevisiae* chrXI; (b) Raw unfiltered nanopore reads fully spanning the 4-kilobase neighborhood of the centromere of *S. cerevisiae* chrXII. In both cases, accessibility is shown at aggregated 10-bp resolution (see Methods section for details) for the single-molecule display, and aggregated over sliding (every 5 bases) 50-bp windows for the genome browser SMAC-seq track.

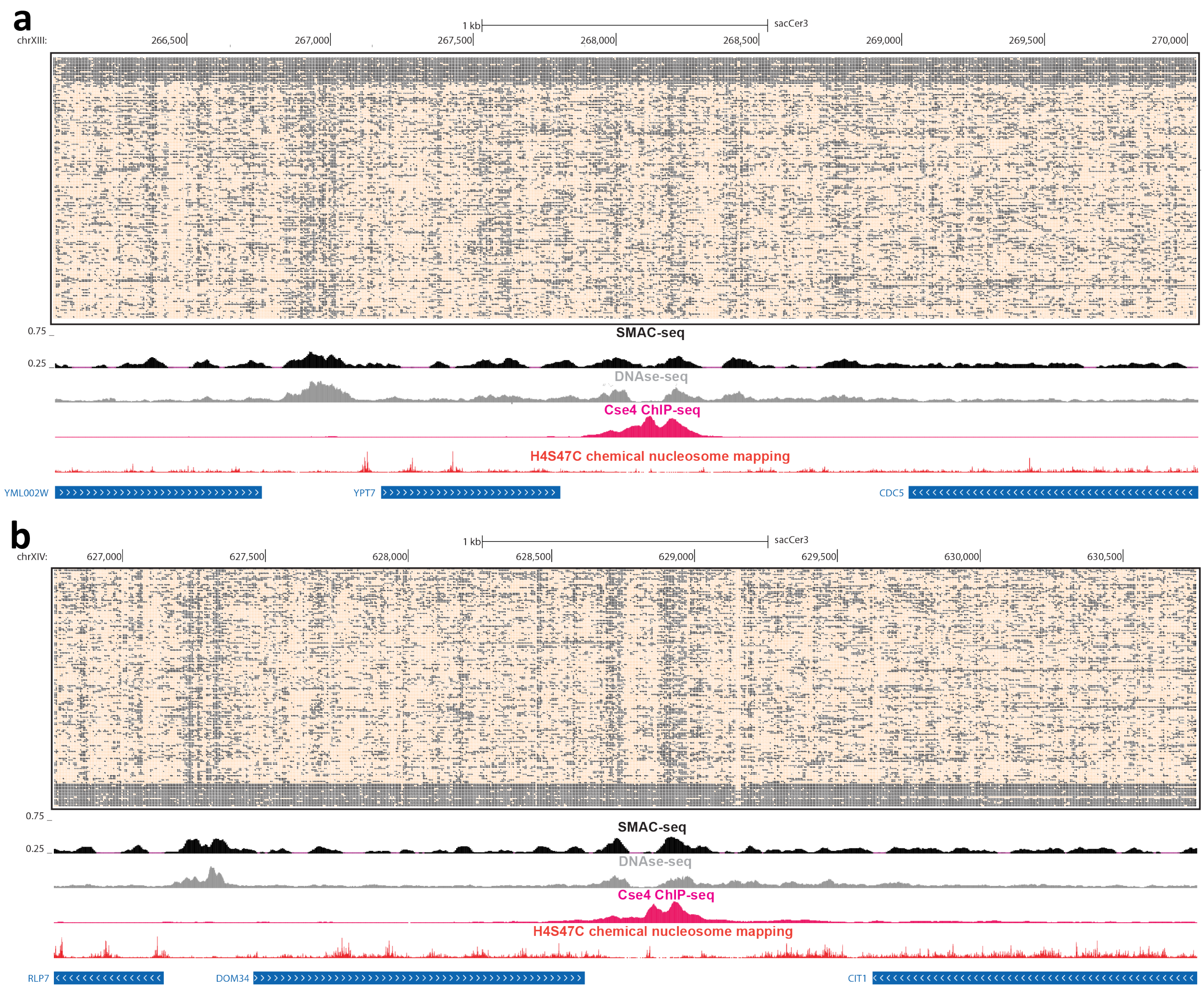


**Supplementary Figure 27: Single-molecule long-read accessibility around well positioned centromeres**. (a) Raw unfiltered nanopore reads fully spanning the 4-kilobase neighborhood of the centromere of *S. cerevisiae* chrXIII; (b) Raw unfiltered nanopore reads fully spanning the 4-kilobase neighborhood of the centromere of *S. cerevisiae* chrXIV. In both cases, accessibility is shown at aggregated 10-bp resolution (see Methods section for details) for the single-molecule display, and aggregated over sliding (every 5 bases) 50-bp windows for the genome browser SMAC-seq track.

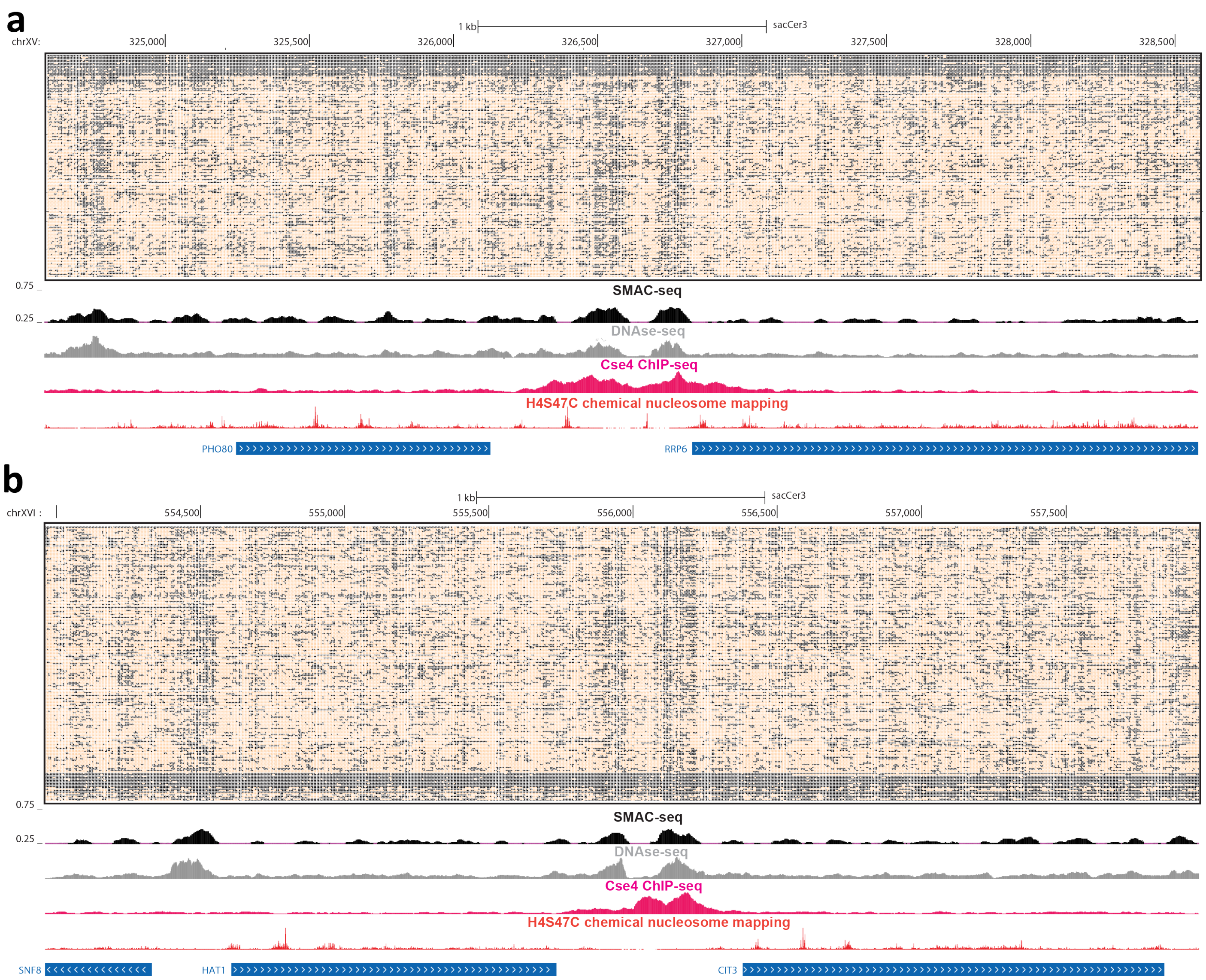


**Supplementary Figure 28: Single-molecule long-read accessibility around well positioned centromeres**. (a) Raw unfiltered nanopore reads fully spanning the 4-kilobase neighborhood of the centromere of *S. cerevisiae* chrXV; (b) Raw unfiltered nanopore reads fully spanning the 4-kilobase neighborhood of the centromere of *S. cerevisiae* chrXVI. In both cases, accessibility is shown at aggregated 10-bp resolution (see Methods section for details) for the single-molecule display, and aggregated over sliding (every 5 bases) 50-bp windows for the genome browser SMAC-seq track.

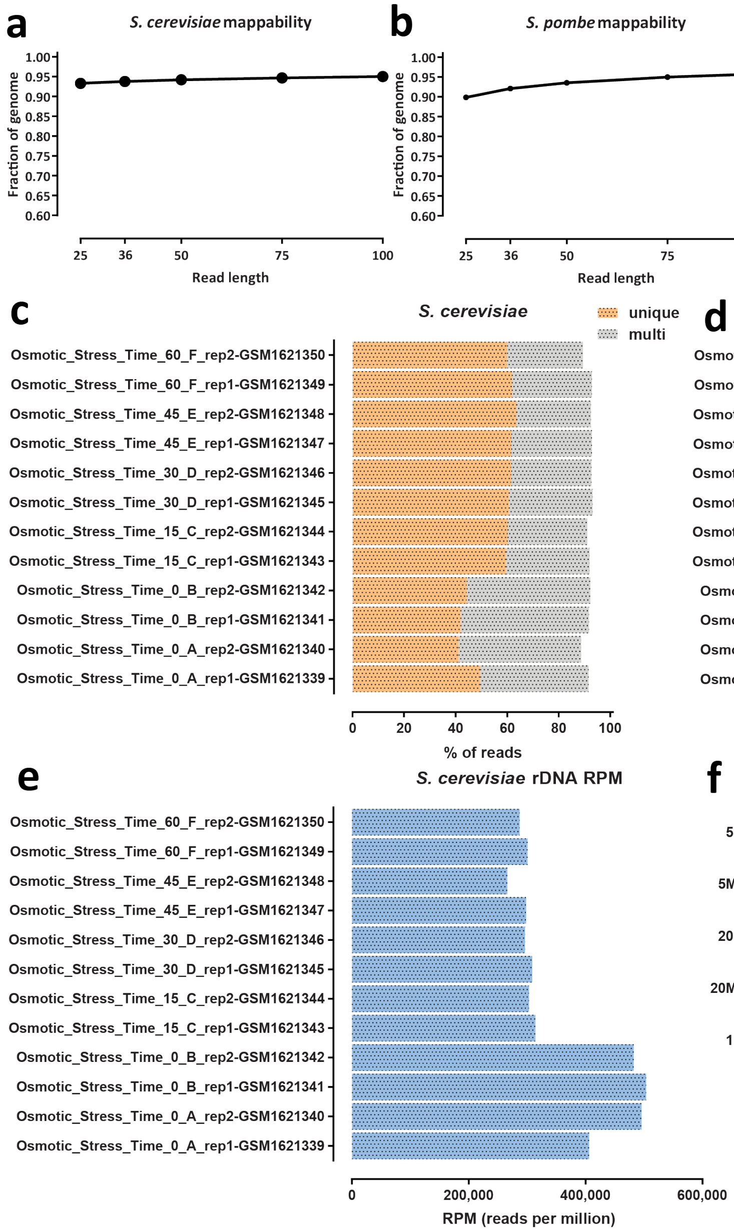


**Supplementary Figure 29: Ribosomal DNA arrays are highly enriched for chromatin accessibility as measured by ATAC-seq**. (a) Unique read mappability of the *Saccharomyces cerevisiae* genome as a function of read length (b) Unique read mappability of the *Schizosaccharomyces pombe* genome as a function of read length (c and d) Enrichment of multimapping reads in *Saccharomyces cerevisiae* ATAC-seq datasets (e) ATAC-seq multireads are highly enriched for rDNA-mapping reads (f) Enrichment of multimapping reads in *Schizosaccharomyces pombe* ATAC-seq datasets ATAC-seq datasets were obtained from Schep et al. 2015[51]

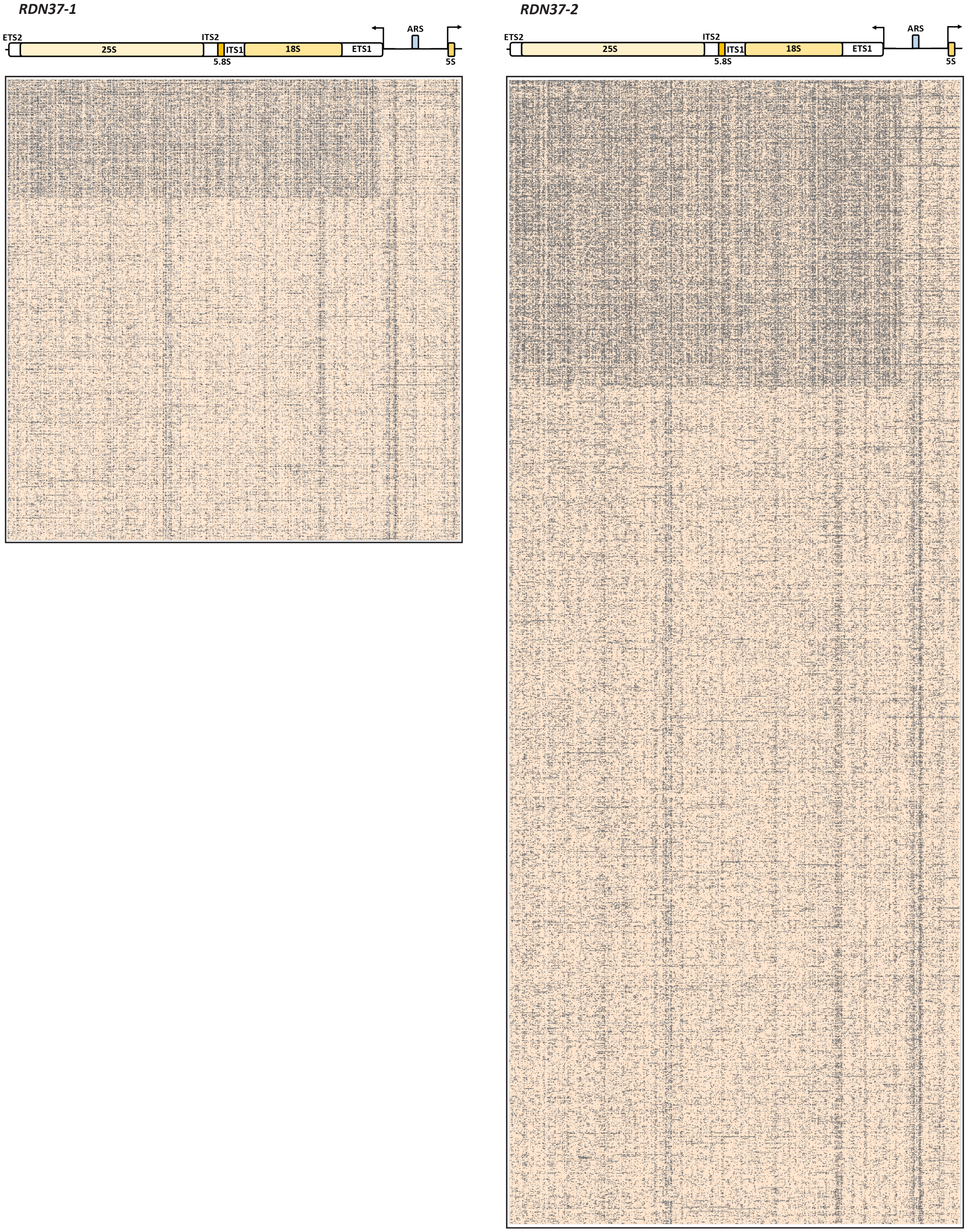


**Supplementary Figure 30: SMAC-seq reveals the distribution of alternative chromatin states of rDNA arrays. Shown are all reads covering the *RDN37-1* and *RDN37-2* arrays in the *RDN1* locus in the "diamide 30 min rep1" experiment (unfiltered reads, "aggregate" signal).**

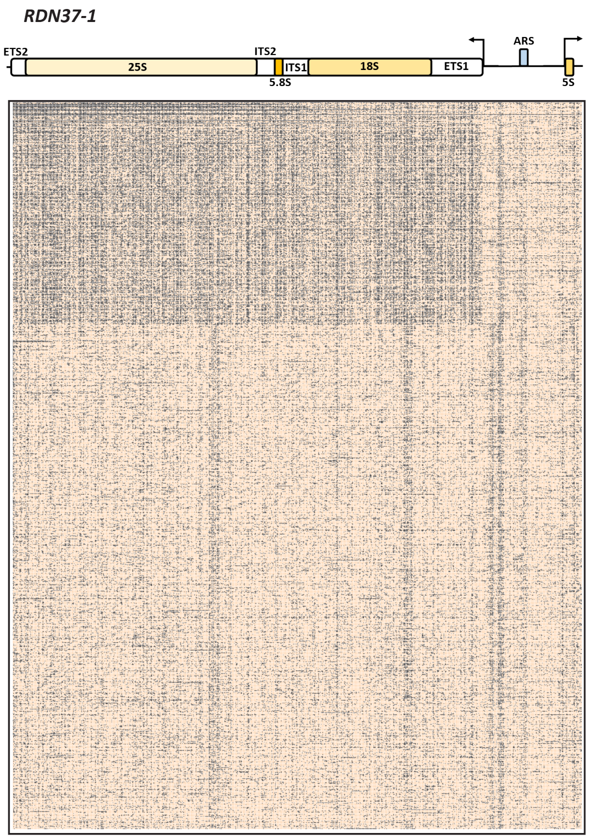
RDN37-1
ETS2
25S
ITS2
ITS1
5.8S
18S
ETS1
ARS
5S

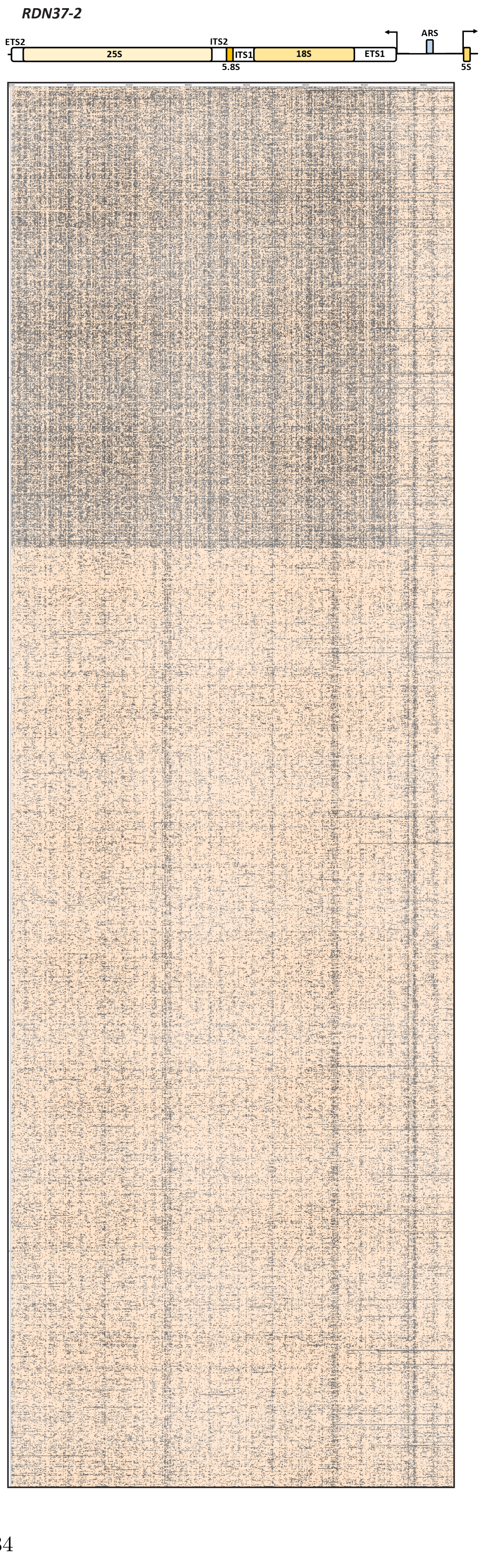
RDN37-2
ETS2
25S
ITS2
ITS1
5.8S
18S
ETS1
ARS
5S

**Supplementary Figure 31** ***(preceding page)*****: SMAC-seq reveals the distribution of alternative chromatin states of rDNA arrays. Shown are all reads covering the** ***RDN37-1*** **and** ***RDN37-2*** **arrays in the** ***RDN1*** **locus in the "diamide 60 min rep1" experiment (unfiltered reads, "aggregate" signal).**

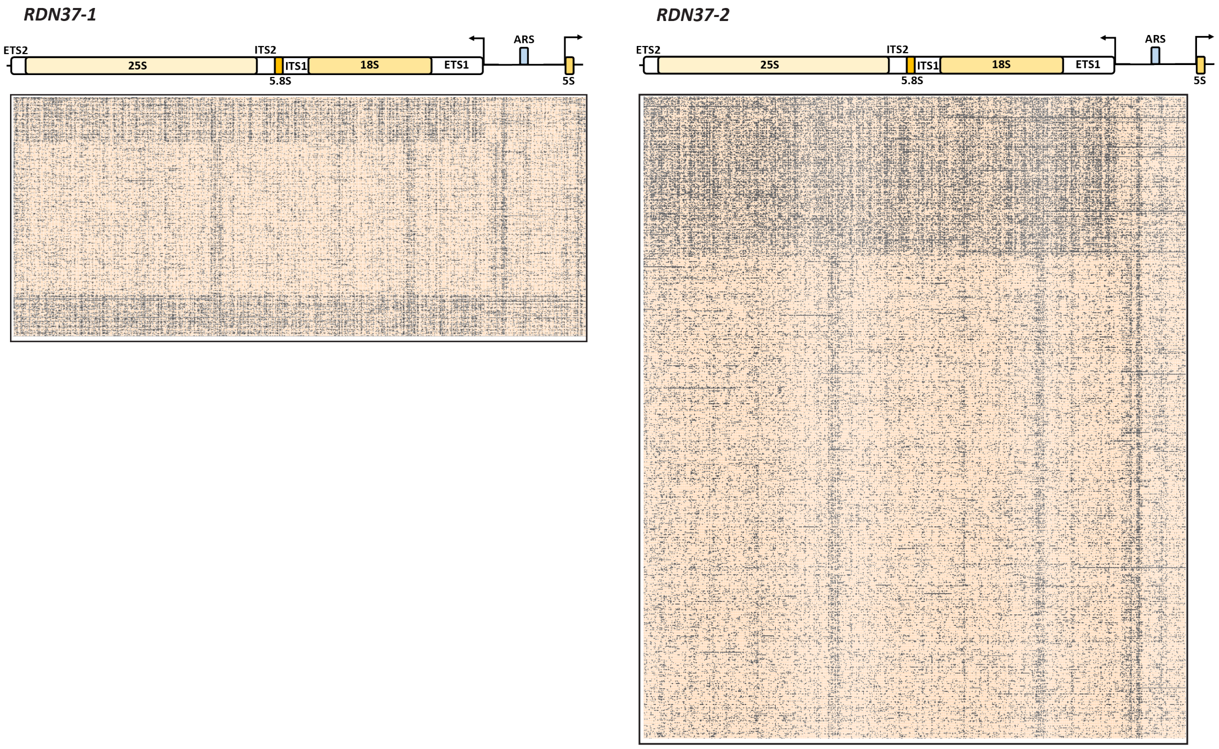


**Supplementary Figure 32: SMAC-seq reveals the distribution of alternative chromatin states of rDNA arrays. Shown are all reads covering the** ***RDN37-1*** **and** ***RDN37-2*** **arrays in the** ***RDN1*** **locus in the "diamide 0 min rep1" experiment (unfiltered reads, "aggregate" signal).**

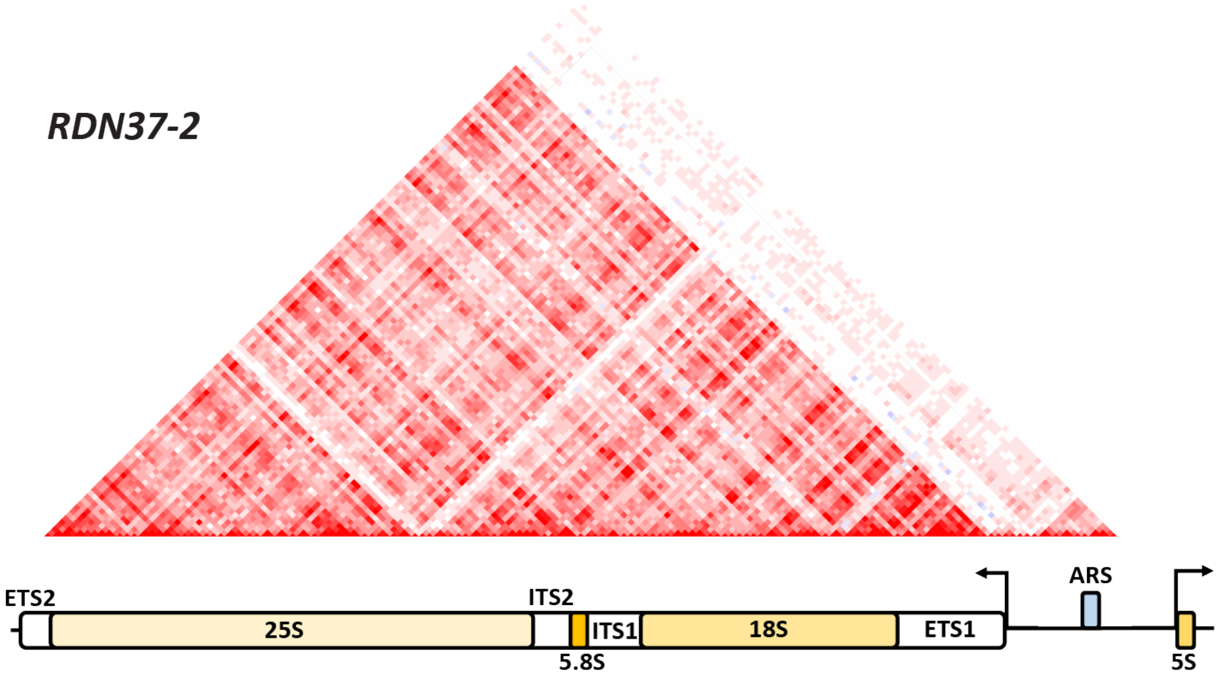


**Supplementary Figure 33: NMI profile for the *RDN37-2* array, as in Figure 3b.**

| # informative positions: | GC | | | | | | | m6A | | | | | | | GC+m6A | | | | | | |
|---|---|---|---|---|---|---|---|---|---|---|---|---|---|---|---|---|---|---|---|---|---|
| | 0 | 1 | 2 | 3 | 4-5 | 6-10 | >10 | 0 | 1 | 2 | 3 | 4-5 | 6-10 | >10 | 0 | 1 | 2 | 3 | 4-5 | 6-10 | >10 |
| ABF1 | 0.56 | 0.33 | 0.10 | 0.01 | 0.00 | 0.00 | 0.00 | 0.00 | 0.02 | 0.08 | 0.17 | 0.48 | 0.25 | 0.00 | 0.00 | 0.01 | 0.03 | 0.12 | 0.46 | 0.38 | 0.00 |
| ABF2 | 0.78 | 0.22 | 0.00 | 0.00 | 0.00 | 0.00 | 0.00 | 0.00 | 0.00 | 0.48 | 0.42 | 0.09 | 0.00 | 0.00 | 0.00 | 0.00 | 0.33 | 0.51 | 0.16 | 0.00 | 0.00 |
| ACA1 | 1.00 | 0.00 | 0.00 | 0.00 | 0.00 | 0.00 | 0.00 | 0.00 | 0.00 | 0.36 | 0.50 | 0.14 | 0.00 | 0.00 | 0.00 | 0.00 | 0.36 | 0.50 | 0.14 | 0.00 | 0.00 |
| ACE2 | 0.00 | 0.79 | 0.21 | 0.00 | 0.00 | 0.00 | 0.00 | 0.31 | 0.16 | 0.17 | 0.17 | 0.20 | 0.00 | 0.00 | 0.00 | 0.23 | 0.20 | 0.14 | 0.37 | 0.06 | 0.00 |
| ADR1 | 0.92 | 0.08 | 0.00 | 0.00 | 0.00 | 0.00 | 0.00 | 0.28 | 0.29 | 0.27 | 0.14 | 0.03 | 0.00 | 0.00 | 0.24 | 0.29 | 0.30 | 0.14 | 0.03 | 0.00 | 0.00 |
| AFT1 | 0.00 | 1.00 | 0.00 | 0.00 | 0.00 | 0.00 | 0.00 | 0.00 | 0.35 | 0.41 | 0.20 | 0.04 | 0.00 | 0.00 | 0.00 | 0.00 | 0.35 | 0.41 | 0.24 | 0.00 | 0.00 |
| ARG5,6 | 0.12 | 0.61 | 0.25 | 0.02 | 0.00 | 0.00 | 0.00 | 0.16 | 0.24 | 0.22 | 0.15 | 0.23 | 0.00 | 0.00 | 0.01 | 0.09 | 0.17 | 0.29 | 0.39 | 0.04 | 0.00 |
| ARG80 | 0.31 | 0.43 | 0.20 | 0.06 | 0.01 | 0.00 | 0.00 | 0.04 | 0.12 | 0.17 | 0.17 | 0.34 | 0.16 | 0.00 | 0.01 | 0.04 | 0.09 | 0.16 | 0.39 | 0.31 | 0.00 |
| ARG81 | 0.44 | 0.40 | 0.16 | 0.00 | 0.00 | 0.00 | 0.00 | 0.15 | 0.51 | 0.29 | 0.05 | 0.00 | 0.00 | 0.00 | 0.03 | 0.24 | 0.47 | 0.26 | 0.00 | 0.00 | 0.00 |
| ARO80 | 0.61 | 0.32 | 0.06 | 0.00 | 0.00 | 0.00 | 0.00 | 0.05 | 0.17 | 0.27 | 0.22 | 0.25 | 0.03 | 0.00 | 0.02 | 0.10 | 0.23 | 0.26 | 0.35 | 0.04 | 0.00 |
| ARR1 | 1.00 | 0.00 | 0.00 | 0.00 | 0.00 | 0.00 | 0.00 | 0.00 | 0.00 | 0.00 | 0.28 | 0.72 | 0.00 | 0.00 | 0.00 | 0.00 | 0.00 | 0.28 | 0.72 | 0.00 | 0.00 |
| AZF1 | 0.96 | 0.04 | 0.00 | 0.00 | 0.00 | 0.00 | 0.00 | 0.43 | 0.06 | 0.00 | 0.00 | 0.00 | 0.51 | 0.00 | 0.41 | 0.08 | 0.00 | 0.00 | 0.00 | 0.51 | 0.00 |
| CAD1 | 1.00 | 0.00 | 0.00 | 0.00 | 0.00 | 0.00 | 0.00 | 0.00 | 0.00 | 0.07 | 0.53 | 0.41 | 0.00 | 0.00 | 0.00 | 0.00 | 0.07 | 0.53 | 0.41 | 0.00 | 0.00 |
| CAT8 | 0.39 | 0.41 | 0.16 | 0.04 | 0.00 | 0.00 | 0.00 | 0.12 | 0.24 | 0.24 | 0.20 | 0.19 | 0.01 | 0.00 | 0.02 | 0.10 | 0.23 | 0.27 | 0.34 | 0.03 | 0.00 |
| CBF1 | 0.60 | 0.39 | 0.01 | 0.00 | 0.00 | 0.00 | 0.00 | 0.00 | 0.37 | 0.40 | 0.21 | 0.02 | 0.00 | 0.00 | 0.00 | 0.16 | 0.45 | 0.32 | 0.07 | 0.00 | 0.00 |
| CEP3 | 0.93 | 0.07 | 0.00 | 0.00 | 0.00 | 0.00 | 0.00 | 0.00 | 0.12 | 0.47 | 0.34 | 0.06 | 0.00 | 0.00 | 0.00 | 0.10 | 0.45 | 0.37 | 0.08 | 0.00 | 0.00 |
| CHA4 | 0.04 | 0.47 | 0.36 | 0.11 | 0.02 | 0.00 | 0.00 | 0.02 | 0.04 | 0.13 | 0.14 | 0.34 | 0.31 | 0.01 | 0.00 | 0.01 | 0.03 | 0.06 | 0.29 | 0.58 | 0.03 |
| CIN5 | 0.97 | 0.03 | 0.00 | 0.00 | 0.00 | 0.00 | 0.00 | 0.00 | 0.00 | 0.06 | 0.52 | 0.42 | 0.00 | 0.00 | 0.00 | 0.00 | 0.05 | 0.52 | 0.44 | 0.00 | 0.00 |
| CRZ1 | 0.00 | 0.93 | 0.07 | 0.00 | 0.00 | 0.00 | 0.00 | 0.35 | 0.26 | 0.22 | 0.14 | 0.04 | 0.00 | 0.00 | 0.00 | 0.33 | 0.24 | 0.24 | 0.19 | 0.00 | 0.00 |
| CST6 | 1.00 | 0.00 | 0.00 | 0.00 | 0.00 | 0.00 | 0.00 | 0.00 | 0.02 | 0.47 | 0.48 | 0.03 | 0.00 | 0.00 | 0.00 | 0.02 | 0.47 | 0.48 | 0.03 | 0.00 | 0.00 |
| CUP2 | 0.15 | 0.74 | 0.10 | 0.00 | 0.00 | 0.00 | 0.00 | 0.32 | 0.18 | 0.00 | 0.00 | 0.35 | 0.15 | 0.00 | 0.06 | 0.23 | 0.21 | 0.00 | 0.20 | 0.31 | 0.00 |
| CUP9 | 0.96 | 0.04 | 0.00 | 0.00 | 0.00 | 0.00 | 0.00 | 0.00 | 0.08 | 0.22 | 0.39 | 0.31 | 0.00 | 0.00 | 0.00 | 0.08 | 0.20 | 0.39 | 0.33 | 0.00 | 0.00 |
| DAL80 | 0.35 | 0.61 | 0.04 | 0.00 | 0.00 | 0.00 | 0.00 | 0.00 | 0.16 | 0.49 | 0.29 | 0.06 | 0.00 | 0.00 | 0.00 | 0.07 | 0.23 | 0.44 | 0.26 | 0.00 | 0.00 |
| DAL81 | 0.01 | 0.22 | 0.43 | 0.24 | 0.10 | 0.00 | 0.00 | 0.05 | 0.15 | 0.22 | 0.26 | 0.27 | 0.05 | 0.00 | 0.00 | 0.00 | 0.05 | 0.12 | 0.45 | 0.38 | 0.00 |
| DAL82 | 0.34 | 0.53 | 0.14 | 0.00 | 0.00 | 0.00 | 0.00 | 0.00 | 0.05 | 0.18 | 0.29 | 0.44 | 0.03 | 0.00 | 0.00 | 0.00 | 0.07 | 0.22 | 0.57 | 0.14 | 0.00 |
| DAT1 | 0.99 | 0.01 | 0.00 | 0.00 | 0.00 | 0.00 | 0.00 | 0.32 | 0.13 | 0.04 | 0.00 | 0.00 | 0.50 | 0.00 | 0.32 | 0.14 | 0.04 | 0.00 | 0.00 | 0.50 | 0.00 |
| DOT6 | 0.24 | 0.43 | 0.24 | 0.08 | 0.01 | 0.00 | 0.00 | 0.00 | 0.03 | 0.05 | 0.13 | 0.36 | 0.42 | 0.01 | 0.00 | 0.01 | 0.02 | 0.04 | 0.26 | 0.65 | 0.02 |
| ECM22 | 0.36 | 0.36 | 0.22 | 0.05 | 0.00 | 0.00 | 0.00 | 0.03 | 0.13 | 0.20 | 0.21 | 0.32 | 0.10 | 0.00 | 0.00 | 0.03 | 0.08 | 0.23 | 0.47 | 0.19 | 0.00 |
| ECM23 | 0.86 | 0.14 | 0.00 | 0.00 | 0.00 | 0.00 | 0.00 | 0.00 | 0.00 | 0.35 | 0.35 | 0.29 | 0.01 | 0.00 | 0.00 | 0.00 | 0.29 | 0.35 | 0.35 | 0.01 | 0.00 |
| EDS1 | 1.00 | 0.00 | 0.00 | 0.00 | 0.00 | 0.00 | 0.00 | 0.09 | 0.25 | 0.15 | 0.01 | 0.40 | 0.10 | 0.00 | 0.09 | 0.25 | 0.15 | 0.01 | 0.40 | 0.10 | 0.00 |
| ERT1 | 0.87 | 0.13 | 0.00 | 0.00 | 0.00 | 0.00 | 0.00 | 0.11 | 0.28 | 0.21 | 0.31 | 0.09 | 0.00 | 0.00 | 0.07 | 0.30 | 0.20 | 0.30 | 0.13 | 0.00 | 0.00 |
| FHL1 | 0.08 | 0.68 | 0.20 | 0.03 | 0.00 | 0.00 | 0.00 | 0.01 | 0.03 | 0.07 | 0.13 | 0.28 | 0.46 | 0.02 | 0.00 | 0.01 | 0.03 | 0.06 | 0.24 | 0.62 | 0.05 |
| FKH1 | 0.89 | 0.11 | 0.00 | 0.00 | 0.00 | 0.00 | 0.00 | 0.01 | 0.20 | 0.20 | 0.11 | 0.27 | 0.21 | 0.00 | 0.01 | 0.18 | 0.21 | 0.11 | 0.25 | 0.24 | 0.00 |
| FKH2 | 0.74 | 0.23 | 0.03 | 0.00 | 0.00 | 0.00 | 0.00 | 0.01 | 0.04 | 0.08 | 0.11 | 0.19 | 0.33 | 0.24 | 0.01 | 0.03 | 0.07 | 0.10 | 0.21 | 0.31 | 0.27 |
| FYV5 | 0.65 | 0.29 | 0.05 | 0.01 | 0.00 | 0.00 | 0.00 | 0.00 | 0.00 | 0.01 | 0.04 | 0.25 | 0.67 | 0.03 | 0.00 | 0.00 | 0.00 | 0.02 | 0.20 | 0.73 | 0.04 |
| FZF1 | 0.38 | 0.45 | 0.13 | 0.04 | 0.00 | 0.00 | 0.00 | 0.00 | 0.01 | 0.03 | 0.05 | 0.21 | 0.59 | 0.10 | 0.00 | 0.00 | 0.01 | 0.03 | 0.15 | 0.66 | 0.15 |
| GAL4 | 0.22 | 0.42 | 0.23 | 0.10 | 0.02 | 0.00 | 0.00 | 0.04 | 0.17 | 0.24 | 0.21 | 0.29 | 0.05 | 0.00 | 0.00 | 0.03 | 0.13 | 0.20 | 0.46 | 0.19 | 0.00 |
| GAT1 | 1.00 | 0.00 | 0.00 | 0.00 | 0.00 | 0.00 | 0.00 | 0.00 | 0.00 | 0.09 | 0.27 | 0.59 | 0.05 | 0.00 | 0.00 | 0.00 | 0.09 | 0.27 | 0.59 | 0.05 | 0.00 |
| GAT3 | 0.86 | 0.14 | 0.00 | 0.00 | 0.00 | 0.00 | 0.00 | 0.00 | 0.04 | 0.32 | 0.40 | 0.24 | 0.00 | 0.00 | 0.00 | 0.04 | 0.24 | 0.43 | 0.29 | 0.00 | 0.00 |
| GAT4 | 0.86 | 0.14 | 0.00 | 0.00 | 0.00 | 0.00 | 0.00 | 0.00 | 0.04 | 0.29 | 0.30 | 0.35 | 0.02 | 0.00 | 0.00 | 0.03 | 0.24 | 0.29 | 0.41 | 0.02 | 0.00 |
| GCN4 | 0.96 | 0.04 | 0.00 | 0.00 | 0.00 | 0.00 | 0.00 | 0.00 | 0.00 | 0.55 | 0.36 | 0.09 | 0.00 | 0.00 | 0.00 | 0.00 | 0.51 | 0.40 | 0.09 | 0.00 | 0.00 |
| GCR1 | 0.00 | 1.00 | 0.00 | 0.00 | 0.00 | 0.00 | 0.00 | 0.27 | 0.22 | 0.02 | 0.21 | 0.29 | 0.00 | 0.00 | 0.00 | 0.27 | 0.22 | 0.02 | 0.44 | 0.06 | 0.00 |
| GCR2 | 0.00 | 1.00 | 0.00 | 0.00 | 0.00 | 0.00 | 0.00 | 0.25 | 0.22 | 0.12 | 0.21 | 0.20 | 0.00 | 0.00 | 0.00 | 0.25 | 0.22 | 0.12 | 0.37 | 0.04 | 0.00 |
| GIS1 | 1.00 | 0.00 | 0.00 | 0.00 | 0.00 | 0.00 | 0.00 | 0.00 | 0.13 | 0.50 | 0.35 | 0.03 | 0.00 | 0.00 | 0.00 | 0.13 | 0.50 | 0.35 | 0.03 | 0.00 | 0.00 |
| GSM1 | 0.57 | 0.36 | 0.07 | 0.01 | 0.00 | 0.00 | 0.00 | 0.00 | 0.01 | 0.06 | 0.09 | 0.36 | 0.48 | 0.01 | 0.00 | 0.00 | 0.03 | 0.07 | 0.29 | 0.60 | 0.01 |
| HAC1 | 0.66 | 0.31 | 0.03 | 0.00 | 0.00 | 0.00 | 0.00 | 0.03 | 0.14 | 0.29 | 0.31 | 0.22 | 0.01 | 0.00 | 0.02 | 0.07 | 0.22 | 0.36 | 0.31 | 0.02 | 0.00 |
| HAL9 | 0.20 | 0.36 | 0.28 | 0.12 | 0.04 | 0.00 | 0.00 | 0.04 | 0.09 | 0.16 | 0.23 | 0.37 | 0.11 | 0.00 | 0.00 | 0.01 | 0.05 | 0.15 | 0.44 | 0.35 | 0.00 |
| HAP1 | 0.60 | 0.34 | 0.06 | 0.00 | 0.00 | 0.00 | 0.00 | 0.04 | 0.13 | 0.28 | 0.26 | 0.25 | 0.02 | 0.00 | 0.02 | 0.11 | 0.20 | 0.26 | 0.35 | 0.06 | 0.00 |
| HAP3 | 0.47 | 0.44 | 0.08 | 0.00 | 0.00 | 0.00 | 0.00 | 0.00 | 0.05 | 0.13 | 0.18 | 0.40 | 0.24 | 0.00 | 0.00 | 0.02 | 0.07 | 0.14 | 0.40 | 0.36 | 0.00 |
| HAP4 | 0.70 | 0.27 | 0.03 | 0.00 | 0.00 | 0.00 | 0.00 | 0.00 | 0.07 | 0.13 | 0.15 | 0.35 | 0.29 | 0.00 | 0.00 | 0.04 | 0.10 | 0.16 | 0.35 | 0.35 | 0.00 |
| HAP5 | 0.41 | 0.41 | 0.16 | 0.02 | 0.00 | 0.00 | 0.00 | 0.01 | 0.14 | 0.21 | 0.17 | 0.28 | 0.19 | 0.00 | 0.00 | 0.04 | 0.14 | 0.19 | 0.32 | 0.30 | 0.00 |
| HCM1 | 1.00 | 0.00 | 0.00 | 0.00 | 0.00 | 0.00 | 0.00 | 0.16 | 0.26 | 0.08 | 0.00 | 0.13 | 0.36 | 0.00 | 0.16 | 0.26 | 0.08 | 0.00 | 0.13 | 0.36 | 0.00 |
| HMRA1 | 0.00 | 1.00 | 0.00 | 0.00 | 0.00 | 0.00 | 0.00 | 0.00 | 0.21 | 0.21 | 0.31 | 0.27 | 0.00 | 0.00 | 0.00 | 0.00 | 0.21 | 0.21 | 0.53 | 0.05 | 0.00 |
| HMRA2 | 0.91 | 0.09 | 0.00 | 0.00 | 0.00 | 0.00 | 0.00 | 0.00 | 0.00 | 0.24 | 0.44 | 0.32 | 0.00 | 0.00 | 0.00 | 0.00 | 0.22 | 0.41 | 0.37 | 0.00 | 0.00 |
| HMS1 | 0.90 | 0.10 | 0.00 | 0.00 | 0.00 | 0.00 | 0.00 | 0.00 | 0.38 | 0.43 | 0.16 | 0.03 | 0.00 | 0.00 | 0.00 | 0.32 | 0.44 | 0.21 | 0.03 | 0.00 | 0.00 |
| HMS2 | 0.93 | 0.07 | 0.00 | 0.00 | 0.00 | 0.00 | 0.00 | 0.09 | 0.17 | 0.18 | 0.06 | 0.35 | 0.15 | 0.00 | 0.09 | 0.16 | 0.18 | 0.08 | 0.33 | 0.17 | 0.00 |
| HSF1 | 0.99 | 0.01 | 0.00 | 0.00 | 0.00 | 0.00 | 0.00 | 0.00 | 0.00 | 0.15 | 0.51 | 0.34 | 0.00 | 0.00 | 0.00 | 0.00 | 0.15 | 0.50 | 0.35 | 0.00 | 0.00 |
| IME1 | 0.00 | 0.76 | 0.23 | 0.01 | 0.00 | 0.00 | 0.00 | 0.32 | 0.40 | 0.23 | 0.05 | 0.00 | 0.00 | 0.00 | 0.00 | 0.24 | 0.35 | 0.30 | 0.11 | 0.00 | 0.00 |
| IME4 | 0.81 | 0.19 | 0.00 | 0.00 | 0.00 | 0.00 | 0.00 | 0.19 | 0.23 | 0.06 | 0.00 | 0.04 | 0.47 | 0.00 | 0.12 | 0.28 | 0.09 | 0.00 | 0.04 | 0.47 | 0.00 |
| INO2 | 0.20 | 0.75 | 0.05 | 0.00 | 0.00 | 0.00 | 0.00 | 0.00 | 0.11 | 0.35 | 0.47 | 0.08 | 0.00 | 0.00 | 0.00 | 0.00 | 0.16 | 0.39 | 0.46 | 0.00 | 0.00 |
| INO4 | 0.42 | 0.54 | 0.04 | 0.00 | 0.00 | 0.00 | 0.00 | 0.00 | 0.06 | 0.54 | 0.30 | 0.10 | 0.00 | 0.00 | 0.00 | 0.00 | 0.27 | 0.48 | 0.25 | 0.00 | 0.00 |
| IXR1 | 0.22 | 0.55 | 0.21 | 0.02 | 0.00 | 0.00 | 0.00 | 0.07 | 0.15 | 0.22 | 0.17 | 0.26 | 0.12 | 0.00 | 0.01 | 0.07 | 0.14 | 0.19 | 0.34 | 0.24 | 0.00 |
| KAR4 | 0.47 | 0.40 | 0.12 | 0.01 | 0.00 | 0.00 | 0.00 | 0.03 | 0.10 | 0.15 | 0.14 | 0.25 | 0.33 | 0.00 | 0.01 | 0.05 | 0.12 | 0.17 | 0.23 | 0.42 | 0.01 |
| LEU3 | 0.39 | 0.39 | 0.16 | 0.06 | 0.00 | 0.00 | 0.00 | 0.21 | 0.43 | 0.27 | 0.09 | 0.00 | 0.00 | 0.00 | 0.06 | 0.22 | 0.35 | 0.26 | 0.11 | 0.00 | 0.00 |
| LYS14 | 0.55 | 0.36 | 0.09 | 0.01 | 0.00 | 0.00 | 0.00 | 0.00 | 0.04 | 0.11 | 0.22 | 0.48 | 0.15 | 0.00 | 0.00 | 0.02 | 0.07 | 0.16 | 0.50 | 0.25 | 0.00 |
| MAC1 | 0.00 | 0.98 | 0.02 | 0.00 | 0.00 | 0.00 | 0.00 | 0.17 | 0.18 | 0.11 | 0.16 | 0.32 | 0.06 | 0.00 | 0.00 | 0.17 | 0.18 | 0.11 | 0.33 | 0.21 | 0.00 |
| MATALPHA1 | 0.57 | 0.34 | 0.08 | 0.01 | 0.00 | 0.00 | 0.00 | 0.00 | 0.00 | 0.02 | 0.05 | 0.20 | 0.63 | 0.10 | 0.00 | 0.00 | 0.01 | 0.03 | 0.15 | 0.68 | 0.13 |
| MBP1 | 0.00 | 1.00 | 0.00 | 0.00 | 0.00 | 0.00 | 0.00 | 0.15 | 0.31 | 0.36 | 0.15 | 0.03 | 0.00 | 0.00 | 0.00 | 0.15 | 0.31 | 0.36 | 0.18 | 0.00 | 0.00 |
| MCM1 | 0.69 | 0.27 | 0.03 | 0.00 | 0.00 | 0.00 | 0.00 | 0.00 | 0.01 | 0.02 | 0.07 | 0.29 | 0.59 | 0.01 | 0.00 | 0.00 | 0.02 | 0.05 | 0.27 | 0.65 | 0.02 |
| MET31 | 0.35 | 0.56 | 0.09 | 0.00 | 0.00 | 0.00 | 0.00 | 0.14 | 0.35 | 0.28 | 0.20 | 0.03 | 0.00 | 0.00 | 0.01 | 0.22 | 0.34 | 0.29 | 0.15 | 0.00 | 0.00 |
| MET32 | 0.19 | 0.72 | 0.09 | 0.00 | 0.00 | 0.00 | 0.00 | 0.13 | 0.22 | 0.34 | 0.26 | 0.06 | 0.00 | 0.00 | 0.00 | 0.10 | 0.37 | 0.22 | 0.31 | 0.00 | 0.00 |
| MET4 | 0.79 | 0.19 | 0.01 | 0.00 | 0.00 | 0.00 | 0.00 | 0.00 | 0.17 | 0.39 | 0.35 | 0.09 | 0.00 | 0.00 | 0.00 | 0.14 | 0.33 | 0.35 | 0.18 | 0.00 | 0.00 |
| MGA1 | 0.68 | 0.27 | 0.05 | 0.00 | 0.00 | 0.00 | 0.00 | 0.01 | 0.03 | 0.06 | 0.09 | 0.21 | 0.49 | 0.11 | 0.00 | 0.02 | 0.04 | 0.08 | 0.21 | 0.51 | 0.14 |
| MIG1 | 0.58 | 0.34 | 0.07 | 0.00 | 0.00 | 0.00 | 0.00 | 0.36 | 0.41 | 0.20 | 0.03 | 0.00 | 0.00 | 0.00 | 0.14 | 0.41 | 0.37 | 0.08 | 0.00 | 0.00 | 0.00 |
| MIG2 | 0.00 | 0.75 | 0.24 | 0.01 | 0.00 | 0.00 | 0.00 | 0.44 | 0.41 | 0.14 | 0.01 | 0.00 | 0.00 | 0.00 | 0.00 | 0.28 | 0.47 | 0.22 | 0.03 | 0.00 | 0.00 |
| MIG3 | 0.00 | 0.75 | 0.24 | 0.01 | 0.00 | 0.00 | 0.00 | 0.44 | 0.41 | 0.14 | 0.01 | 0.00 | 0.00 | 0.00 | 0.00 | 0.28 | 0.47 | 0.22 | 0.03 | 0.00 | 0.00 |
| MSN1 | 1.00 | 0.00 | 0.00 | 0.00 | 0.00 | 0.00 | 0.00 | 0.00 | 0.08 | 0.38 | 0.36 | 0.19 | 0.00 | 0.00 | 0.00 | 0.08 | 0.38 | 0.36 | 0.19 | 0.00 | 0.00 |
| MSN2 | 0.57 | 0.43 | 0.00 | 0.00 | 0.00 | 0.00 | 0.00 | 0.15 | 0.22 | 0.34 | 0.22 | 0.07 | 0.00 | 0.00 | 0.00 | 0.30 | 0.32 | 0.23 | 0.16 | 0.00 | 0.00 |
| NDT80 | 0.92 | 0.08 | 0.00 | 0.00 | 0.00 | 0.00 | 0.00 | 0.26 | 0.21 | 0.03 | 0.01 | 0.35 | 0.14 | 0.00 | 0.22 | 0.25 | 0.03 | 0.01 | 0.34 | 0.15 | 0.00 |
| NHP10 | 0.58 | 0.35 | 0.07 | 0.00 | 0.00 | 0.00 | 0.00 | 0.36 | 0.40 | 0.20 | 0.05 | 0.00 | 0.00 | 0.00 | 0.13 | 0.39 | 0.39 | 0.09 | 0.00 | 0.00 | 0.00 |
| NHP6A | 0.91 | 0.09 | 0.01 | 0.00 | 0.00 | 0.00 | 0.00 | 0.00 | 0.00 | 0.00 | 0.00 | 0.05 | 0.75 | 0.20 | 0.00 | 0.00 | 0.00 | 0.00 | 0.04 | 0.75 | 0.21 |
| NHP6B | 0.93 | 0.07 | 0.00 | 0.00 | 0.00 | 0.00 | 0.00 | 0.00 | 0.00 | 0.00 | 0.01 | 0.06 | 0.75 | 0.18 | 0.00 | 0.00 | 0.00 | 0.01 | 0.05 | 0.75 | 0.19 |
| NRG1 | 1.00 | 0.00 | 0.00 | 0.00 | 0.00 | 0.00 | 0.00 | 0.00 | 0.34 | 0.44 | 0.20 | 0.02 | 0.00 | 0.00 | 0.00 | 0.34 | 0.44 | 0.20 | 0.02 | 0.00 | 0.00 |
| NRG2 | 0.94 | 0.06 | 0.00 | 0.00 | 0.00 | 0.00 | 0.00 | 0.00 | 0.36 | 0.47 | 0.17 | 0.00 | 0.00 | 0.00 | 0.00 | 0.31 | 0.51 | 0.18 | 0.00 | 0.00 | 0.00 |
| OAF1 | 0.05 | 0.16 | 0.27 | 0.34 | 0.18 | 0.01 | 0.00 | 0.08 | 0.11 | 0.21 | 0.13 | 0.27 | 0.19 | 0.00 | 0.00 | 0.00 | 0.02 | 0.10 | 0.36 | 0.50 | 0.02 |
| OPI1 | 0.80 | 0.19 | 0.01 | 0.00 | 0.00 | 0.00 | 0.00 | 0.22 | 0.23 | 0.29 | 0.20 | 0.05 | 0.00 | 0.00 | 0.15 | 0.27 | 0.25 | 0.24 | 0.09 | 0.00 | 0.00 |
| ORC1 | 0.96 | 0.04 | 0.00 | 0.00 | 0.00 | 0.00 | 0.00 | 0.10 | 0.11 | 0.10 | 0.08 | 0.08 | 0.14 | 0.39 | 0.09 | 0.11 | 0.11 | 0.08 | 0.08 | 0.13 | 0.40 |
| PDC2 | 0.99 | 0.01 | 0.00 | 0.00 | 0.00 | 0.00 | 0.00 | 0.00 | 0.00 | 0.04 | 0.11 | 0.76 | 0.09 | 0.00 | 0.00 | 0.00 | 0.04 | 0.10 | 0.77 | 0.09 | 0.00 |
| PDR1 | 0.24 | 0.49 | 0.26 | 0.01 | 0.00 | 0.00 | 0.00 | 0.31 | 0.39 | 0.22 | 0.08 | 0.00 | 0.00 | 0.00 | 0.00 | 0.28 | 0.41 | 0.23 | 0.07 | 0.00 | 0.00 |
| PDR3 | 0.00 | 0.67 | 0.33 | 0.00 | 0.00 | 0.00 | 0.00 | 0.33 | 0.44 | 0.15 | 0.08 | 0.00 | 0.00 | 0.00 | 0.00 | 0.20 | 0.40 | 0.28 | 0.12 | 0.00 | 0.00 |
| PDR8 | 0.47 | 0.35 | 0.18 | 0.00 | 0.00 | 0.00 | 0.00 | 0.17 | 0.44 | 0.29 | 0.10 | 0.00 | 0.00 | 0.00 | 0.00 | 0.31 | 0.42 | 0.21 | 0.07 | 0.00 | 0.00 |
| PHD1 | 0.00 | 0.34 | 0.59 | 0.07 | 0.00 | 0.00 | 0.00 | 0.29 | 0.45 | 0.22 | 0.04 | 0.00 | 0.00 | 0.00 | 0.00 | 0.12 | 0.28 | 0.39 | 0.21 | 0.00 | 0.00 |
| PHO4 | 0.32 | 0.56 | 0.12 | 0.00 | 0.00 | 0.00 | 0.00 | 0.00 | 0.52 | 0.41 | 0.08 | 0.00 | 0.00 | 0.00 | 0.00 | 0.12 | 0.49 | 0.31 | 0.08 | 0.00 | 0.00 |
| PPR1 | 0.37 | 0.43 | 0.17 | 0.02 | 0.00 | 0.00 | 0.00 | 0.02 | 0.11 | 0.23 | 0.27 | 0.30 | 0.07 | 0.00 | 0.01 | 0.03 | 0.12 | 0.23 | 0.47 | 0.14 | 0.00 |
| PUT3 | 0.24 | 0.45 | 0.23 | 0.08 | 0.01 | 0.00 | 0.00 | 0.13 | 0.20 | 0.20 | 0.27 | 0.17 | 0.02 | 0.00 | 0.02 | 0.09 | 0.16 | 0.22 | 0.42 | 0.08 | 0.00 |
| PZF1 | 0.64 | 0.28 | 0.07 | 0.00 | 0.00 | 0.00 | 0.00 | 0.02 | 0.09 | 0.24 | 0.30 | 0.30 | 0.04 | 0.00 | 0.01 | 0.06 | 0.18 | 0.27 | 0.43 | 0.06 | 0.00 |
| RAP1 | 0.82 | 0.16 | 0.02 | 0.00 | 0.00 | 0.00 | 0.00 | 0.16 | 0.14 | 0.13 | 0.15 | 0.28 | 0.13 | 0.00 | 0.14 | 0.11 | 0.14 | 0.15 | 0.31 | 0.15 | 0.00 |
| RCO1 | 1.00 | 0.00 | 0.00 | 0.00 | 0.00 | 0.00 | 0.00 | 0.00 | 0.00 | 0.26 | 0.21 | 0.49 | 0.04 | 0.00 | 0.00 | 0.00 | 0.26 | 0.21 | 0.49 | 0.04 | 0.00 |
| RDR1 | 0.54 | 0.38 | 0.08 | 0.00 | 0.00 | 0.00 | 0.00 | 0.07 | 0.20 | 0.41 | 0.28 | 0.04 | 0.00 | 0.00 | 0.02 | 0.11 | 0.32 | 0.38 | 0.17 | 0.00 | 0.00 |
| RDS1 | 0.00 | 0.42 | 0.43 | 0.15 | 0.00 | 0.00 | 0.00 | 0.44 | 0.43 | 0.10 | 0.03 | 0.00 | 0.00 | 0.00 | 0.00 | 0.14 | 0.39 | 0.36 | 0.11 | 0.00 | 0.00 |
| RDS2 | 0.48 | 0.42 | 0.09 | 0.01 | 0.00 | 0.00 | 0.00 | 0.02 | 0.14 | 0.22 | 0.22 | 0.32 | 0.08 | 0.00 | 0.01 | 0.05 | 0.17 | 0.21 | 0.42 | 0.14 | 0.00 |
| REB1 | 0.58 | 0.38 | 0.04 | 0.00 | 0.00 | 0.00 | 0.00 | 0.01 | 0.13 | 0.28 | 0.31 | 0.25 | 0.02 | 0.00 | 0.00 | 0.05 | 0.21 | 0.33 | 0.37 | 0.03 | 0.00 |
| REI1 | 0.71 | 0.27 | 0.02 | 0.00 | 0.00 | 0.00 | 0.00 | 0.11 | 0.44 | 0.37 | 0.08 | 0.00 | 0.00 | 0.00 | 0.06 | 0.32 | 0.46 | 0.16 | 0.00 | 0.00 | 0.00 |
| RFX1 | 0.23 | 0.47 | 0.23 | 0.06 | 0.01 | 0.00 | 0.00 | 0.01 | 0.04 | 0.14 | 0.24 | 0.48 | 0.09 | 0.00 | 0.00 | 0.01 | 0.03 | 0.12 | 0.52 | 0.32 | 0.00 |
| RGT1 | 0.59 | 0.33 | 0.07 | 0.00 | 0.00 | 0.00 | 0.00 | 0.00 | 0.01 | 0.03 | 0.07 | 0.30 | 0.57 | 0.02 | 0.00 | 0.01 | 0.02 | 0.04 | 0.24 | 0.66 | 0.02 |
| RIM101 | 0.00 | 0.83 | 0.17 | 0.00 | 0.00 | 0.00 | 0.00 | 0.00 | 0.26 | 0.44 | 0.26 | 0.04 | 0.00 | 0.00 | 0.00 | 0.00 | 0.21 | 0.40 | 0.39 | 0.00 | 0.00 |
| RLM1 | 1.00 | 0.00 | 0.00 | 0.00 | 0.00 | 0.00 | 0.00 | 0.00 | 0.03 | 0.06 | 0.17 | 0.59 | 0.16 | 0.00 | 0.00 | 0.03 | 0.06 | 0.17 | 0.59 | 0.16 | 0.00 |
| RME1 | 0.91 | 0.09 | 0.00 | 0.00 | 0.00 | 0.00 | 0.00 | 0.06 | 0.19 | 0.16 | 0.11 | 0.30 | 0.19 | 0.00 | 0.05 | 0.18 | 0.17 | 0.11 | 0.29 | 0.20 | 0.00 |
| ROX1 | 0.87 | 0.13 | 0.00 | 0.00 | 0.00 | 0.00 | 0.00 | 0.00 | 0.37 | 0.14 | 0.00 | 0.41 | 0.08 | 0.00 | 0.00 | 0.28 | 0.23 | 0.00 | 0.40 | 0.09 | 0.00 |
| RPH1 | 0.90 | 0.10 | 0.00 | 0.00 | 0.00 | 0.00 | 0.00 | 0.00 | 0.17 | 0.47 | 0.29 | 0.07 | 0.00 | 0.00 | 0.00 | 0.13 | 0.45 | 0.33 | 0.09 | 0.00 | 0.00 |
| RPN4 | 0.02 | 0.89 | 0.09 | 0.00 | 0.00 | 0.00 | 0.00 | 0.02 | 0.18 | 0.28 | 0.29 | 0.22 | 0.01 | 0.00 | 0.00 | 0.02 | 0.16 | 0.28 | 0.48 | 0.07 | 0.00 |
| RSC3 | 0.00 | 0.00 | 0.34 | 0.47 | 0.19 | 0.00 | 0.00 | 0.61 | 0.35 | 0.05 | 0.00 | 0.00 | 0.00 | 0.00 | 0.00 | 0.00 | 0.17 | 0.40 | 0.43 | 0.00 | 0.00 |
| RSC30 | 0.00 | 0.02 | 0.23 | 0.47 | 0.26 | 0.02 | 0.00 | 0.36 | 0.29 | 0.11 | 0.10 | 0.11 | 0.03 | 0.00 | 0.00 | 0.00 | 0.08 | 0.28 | 0.36 | 0.27 | 0.01 |
| RSF2 | 0.57 | 0.41 | 0.02 | 0.00 | 0.00 | 0.00 | 0.00 | 0.16 | 0.36 | 0.32 | 0.12 | 0.05 | 0.00 | 0.00 | 0.06 | 0.27 | 0.36 | 0.23 | 0.08 | 0.00 | 0.00 |
| RTG1 | 0.97 | 0.03 | 0.00 | 0.00 | 0.00 | 0.00 | 0.00 | 0.00 | 0.01 | 0.26 | 0.38 | 0.34 | 0.01 | 0.00 | 0.00 | 0.01 | 0.25 | 0.38 | 0.36 | 0.01 | 0.00 |
| RTG3 | 0.57 | 0.24 | 0.13 | 0.05 | 0.01 | 0.00 | 0.00 | 0.00 | 0.04 | 0.22 | 0.28 | 0.31 | 0.15 | 0.00 | 0.00 | 0.03 | 0.15 | 0.24 | 0.31 | 0.26 | 0.01 |
| SFL1 | 0.00 | 0.74 | 0.26 | 0.00 | 0.00 | 0.00 | 0.00 | 0.00 | 0.21 | 0.55 | 0.23 | 0.00 | 0.00 | 0.00 | 0.00 | 0.00 | 0.13 | 0.51 | 0.36 | 0.00 | 0.00 |
| SFP1 | 0.77 | 0.21 | 0.02 | 0.00 | 0.00 | 0.00 | 0.00 | 0.00 | 0.00 | 0.00 | 0.01 | 0.14 | 0.76 | 0.09 | 0.00 | 0.00 | 0.00 | 0.01 | 0.11 | 0.78 | 0.10 |
| SIP4 | 0.43 | 0.50 | 0.07 | 0.01 | 0.00 | 0.00 | 0.00 | 0.02 | 0.17 | 0.28 | 0.28 | 0.24 | 0.00 | 0.00 | 0.00 | 0.05 | 0.22 | 0.30 | 0.40 | 0.03 | 0.00 |
| SKN7 | 0.00 | 0.15 | 0.79 | 0.06 | 0.00 | 0.00 | 0.00 | 0.71 | 0.26 | 0.04 | 0.00 | 0.00 | 0.00 | 0.00 | 0.00 | 0.09 | 0.58 | 0.33 | 0.00 | 0.00 | 0.00 |
| SKO1 | 0.83 | 0.17 | 0.00 | 0.00 | 0.00 | 0.00 | 0.00 | 0.00 | 0.06 | 0.36 | 0.43 | 0.15 | 0.00 | 0.00 | 0.00 | 0.00 | 0.39 | 0.41 | 0.20 | 0.00 | 0.00 |
| SMK1 | 0.29 | 0.48 | 0.18 | 0.04 | 0.00 | 0.00 | 0.00 | 0.00 | 0.01 | 0.04 | 0.10 | 0.44 | 0.41 | 0.00 | 0.00 | 0.00 | 0.01 | 0.04 | 0.30 | 0.66 | 0.00 |
| SMP1 | 0.62 | 0.32 | 0.06 | 0.01 | 0.00 | 0.00 | 0.00 | 0.00 | 0.00 | 0.00 | 0.02 | 0.16 | 0.75 | 0.07 | 0.00 | 0.00 | 0.00 | 0.01 | 0.11 | 0.78 | 0.10 |
| SNF1 | 1.00 | 0.00 | 0.00 | 0.00 | 0.00 | 0.00 | 0.00 | 0.32 | 0.16 | 0.02 | 0.00 | 0.00 | 0.50 | 0.00 | 0.32 | 0.16 | 0.02 | 0.00 | 0.00 | 0.50 | 0.00 |
| SNT2 | 0.01 | 0.35 | 0.49 | 0.14 | 0.02 | 0.00 | 0.00 | 0.01 | 0.09 | 0.23 | 0.27 | 0.35 | 0.05 | 0.00 | 0.00 | 0.00 | 0.02 | 0.11 | 0.52 | 0.34 | 0.00 |
| SOK2 | 0.00 | 0.62 | 0.38 | 0.00 | 0.00 | 0.00 | 0.00 | 0.03 | 0.29 | 0.32 | 0.29 | 0.07 | 0.00 | 0.00 | 0.00 | 0.03 | 0.14 | 0.38 | 0.41 | 0.03 | 0.00 |
| SPT10 | 0.47 | 0.42 | 0.10 | 0.01 | 0.00 | 0.00 | 0.00 | 0.00 | 0.02 | 0.10 | 0.21 | 0.50 | 0.18 | 0.00 | 0.00 | 0.01 | 0.04 | 0.15 | 0.50 | 0.31 | 0.00 |
| SPT15 | 0.83 | 0.15 | 0.02 | 0.00 | 0.00 | 0.00 | 0.00 | 0.00 | 0.00 | 0.00 | 0.00 | 0.06 | 0.82 | 0.12 | 0.00 | 0.00 | 0.00 | 0.00 | 0.04 | 0.83 | 0.13 |
| SPT2 | 0.98 | 0.02 | 0.00 | 0.00 | 0.00 | 0.00 | 0.00 | 0.00 | 0.31 | 0.16 | 0.02 | 0.19 | 0.31 | 0.00 | 0.00 | 0.31 | 0.16 | 0.02 | 0.17 | 0.33 | 0.00 |
| SPT23 | 1.00 | 0.00 | 0.00 | 0.00 | 0.00 | 0.00 | 0.00 | 0.00 | 0.23 | 0.21 | 0.05 | 0.07 | 0.44 | 0.00 | 0.00 | 0.23 | 0.21 | 0.05 | 0.07 | 0.44 | 0.00 |
| SRD1 | 0.98 | 0.02 | 0.00 | 0.00 | 0.00 | 0.00 | 0.00 | 0.00 | 0.00 | 0.38 | 0.36 | 0.25 | 0.00 | 0.00 | 0.00 | 0.00 | 0.36 | 0.39 | 0.25 | 0.00 | 0.00 |
| STB2 | 0.66 | 0.31 | 0.03 | 0.00 | 0.00 | 0.00 | 0.00 | 0.00 | 0.33 | 0.43 | 0.19 | 0.05 | 0.00 | 0.00 | 0.00 | 0.23 | 0.40 | 0.24 | 0.14 | 0.00 | 0.00 |
| STB3 | 0.73 | 0.24 | 0.03 | 0.00 | 0.00 | 0.00 | 0.00 | 0.00 | 0.01 | 0.03 | 0.05 | 0.20 | 0.55 | 0.15 | 0.00 | 0.01 | 0.02 | 0.05 | 0.18 | 0.56 | 0.18 |
| STB4 | 0.81 | 0.19 | 0.00 | 0.00 | 0.00 | 0.00 | 0.00 | 0.00 | 0.21 | 0.49 | 0.25 | 0.04 | 0.00 | 0.00 | 0.00 | 0.16 | 0.44 | 0.33 | 0.07 | 0.00 | 0.00 |
| STB5 | 0.73 | 0.25 | 0.02 | 0.00 | 0.00 | 0.00 | 0.00 | 0.07 | 0.23 | 0.34 | 0.22 | 0.14 | 0.00 | 0.00 | 0.05 | 0.14 | 0.33 | 0.31 | 0.17 | 0.00 | 0.00 |
| STE12 | 1.00 | 0.00 | 0.00 | 0.00 | 0.00 | 0.00 | 0.00 | 0.00 | 0.22 | 0.22 | 0.05 | 0.45 | 0.06 | 0.00 | 0.00 | 0.22 | 0.22 | 0.05 | 0.45 | 0.06 | 0.00 |
| STP1 | 0.00 | 0.00 | 0.82 | 0.17 | 0.01 | 0.00 | 0.00 | 0.41 | 0.22 | 0.21 | 0.13 | 0.03 | 0.00 | 0.00 | 0.00 | 0.00 | 0.32 | 0.25 | 0.40 | 0.03 | 0.00 |
| STP2 | 0.00 | 0.13 | 0.35 | 0.34 | 0.17 | 0.00 | 0.00 | 0.01 | 0.07 | 0.15 | 0.25 | 0.37 | 0.16 | 0.00 | 0.00 | 0.00 | 0.00 | 0.03 | 0.28 | 0.69 | 0.01 |
| STP3 | 0.00 | 0.03 | 0.87 | 0.11 | 0.00 | 0.00 | 0.00 | 0.00 | 0.45 | 0.41 | 0.11 | 0.03 | 0.00 | 0.00 | 0.00 | 0.00 | 0.01 | 0.38 | 0.57 | 0.03 | 0.00 |
| STP4 | 0.00 | 0.17 | 0.75 | 0.08 | 0.00 | 0.00 | 0.00 | 0.00 | 0.42 | 0.39 | 0.16 | 0.03 | 0.00 | 0.00 | 0.00 | 0.00 | 0.04 | 0.42 | 0.51 | 0.03 | 0.00 |
| SUA7 | 0.06 | 0.70 | 0.24 | 0.00 | 0.00 | 0.00 | 0.00 | 0.11 | 0.38 | 0.35 | 0.16 | 0.00 | 0.00 | 0.00 | 0.01 | 0.07 | 0.34 | 0.35 | 0.23 | 0.00 | 0.00 |
| SUM1 | 0.81 | 0.16 | 0.02 | 0.00 | 0.00 | 0.00 | 0.00 | 0.05 | 0.18 | 0.19 | 0.16 | 0.33 | 0.09 | 0.00 | 0.02 | 0.13 | 0.22 | 0.20 | 0.32 | 0.11 | 0.00 |
| SUT1 | 0.00 | 0.68 | 0.29 | 0.03 | 0.00 | 0.00 | 0.00 | 0.44 | 0.36 | 0.19 | 0.02 | 0.00 | 0.00 | 0.00 | 0.00 | 0.25 | 0.41 | 0.29 | 0.05 | 0.00 | 0.00 |
| SUT2 | 0.68 | 0.28 | 0.04 | 0.00 | 0.00 | 0.00 | 0.00 | 0.00 | 0.02 | 0.06 | 0.11 | 0.31 | 0.49 | 0.02 | 0.00 | 0.01 | 0.04 | 0.10 | 0.29 | 0.54 | 0.02 |
| SWI4 | 0.00 | 0.86 | 0.14 | 0.00 | 0.00 | 0.00 | 0.00 | 0.28 | 0.19 | 0.05 | 0.24 | 0.24 | 0.00 | 0.00 | 0.00 | 0.22 | 0.23 | 0.07 | 0.42 | 0.06 | 0.00 |
| SWI5 | 0.00 | 0.93 | 0.07 | 0.00 | 0.00 | 0.00 | 0.00 | 0.24 | 0.21 | 0.11 | 0.23 | 0.21 | 0.00 | 0.00 | 0.00 | 0.21 | 0.23 | 0.10 | 0.41 | 0.05 | 0.00 |
| TBF1 | 0.65 | 0.30 | 0.06 | 0.00 | 0.00 | 0.00 | 0.00 | 0.01 | 0.06 | 0.17 | 0.27 | 0.41 | 0.09 | 0.00 | 0.00 | 0.03 | 0.12 | 0.23 | 0.45 | 0.16 | 0.00 |
| TBS1 | 0.73 | 0.24 | 0.03 | 0.00 | 0.00 | 0.00 | 0.00 | 0.18 | 0.38 | 0.35 | 0.08 | 0.00 | 0.00 | 0.00 | 0.11 | 0.30 | 0.44 | 0.16 | 0.00 | 0.00 | 0.00 |
| TEA1 | 0.49 | 0.39 | 0.11 | 0.02 | 0.00 | 0.00 | 0.00 | 0.00 | 0.00 | 0.01 | 0.05 | 0.30 | 0.62 | 0.01 | 0.00 | 0.00 | 0.01 | 0.03 | 0.19 | 0.74 | 0.03 |
| TEC1 | 0.52 | 0.37 | 0.10 | 0.01 | 0.00 | 0.00 | 0.00 | 0.00 | 0.00 | 0.02 | 0.07 | 0.35 | 0.56 | 0.00 | 0.00 | 0.00 | 0.01 | 0.03 | 0.27 | 0.69 | 0.01 |
| THI2 | 0.21 | 0.38 | 0.24 | 0.12 | 0.04 | 0.00 | 0.00 | 0.04 | 0.10 | 0.15 | 0.16 | 0.36 | 0.18 | 0.00 | 0.00 | 0.02 | 0.08 | 0.13 | 0.37 | 0.39 | 0.01 |
| THO2 | 0.94 | 0.06 | 0.00 | 0.00 | 0.00 | 0.00 | 0.00 | 0.15 | 0.17 | 0.13 | 0.05 | 0.08 | 0.43 | 0.00 | 0.14 | 0.17 | 0.13 | 0.05 | 0.07 | 0.44 | 0.00 |
| TOD6 | 0.23 | 0.44 | 0.25 | 0.07 | 0.01 | 0.00 | 0.00 | 0.00 | 0.02 | 0.05 | 0.12 | 0.38 | 0.43 | 0.00 | 0.00 | 0.00 | 0.02 | 0.04 | 0.26 | 0.67 | 0.02 |
| TOS8 | 0.93 | 0.07 | 0.00 | 0.00 | 0.00 | 0.00 | 0.00 | 0.00 | 0.00 | 0.42 | 0.35 | 0.23 | 0.00 | 0.00 | 0.00 | 0.00 | 0.38 | 0.36 | 0.26 | 0.00 | 0.00 |
| TYE7 | 0.86 | 0.14 | 0.00 | 0.00 | 0.00 | 0.00 | 0.00 | 0.00 | 0.15 | 0.49 | 0.24 | 0.12 | 0.00 | 0.00 | 0.00 | 0.11 | 0.47 | 0.27 | 0.15 | 0.00 | 0.00 |
| UGA3 | 0.00 | 0.39 | 0.58 | 0.03 | 0.00 | 0.00 | 0.00 | 0.25 | 0.56 | 0.18 | 0.02 | 0.00 | 0.00 | 0.00 | 0.00 | 0.09 | 0.36 | 0.42 | 0.13 | 0.00 | 0.00 |
| UME1 | 0.99 | 0.01 | 0.00 | 0.00 | 0.00 | 0.00 | 0.00 | 0.00 | 0.28 | 0.19 | 0.04 | 0.12 | 0.39 | 0.00 | 0.00 | 0.27 | 0.19 | 0.04 | 0.11 | 0.39 | 0.00 |
| UME6 | 0.00 | 0.36 | 0.56 | 0.08 | 0.00 | 0.00 | 0.00 | 0.18 | 0.47 | 0.31 | 0.04 | 0.00 | 0.00 | 0.00 | 0.00 | 0.01 | 0.32 | 0.43 | 0.24 | 0.00 | 0.00 |
| UPC2 | 1.00 | 0.00 | 0.00 | 0.00 | 0.00 | 0.00 | 0.00 | 0.00 | 0.00 | 0.25 | 0.42 | 0.33 | 0.00 | 0.00 | 0.00 | 0.00 | 0.25 | 0.42 | 0.33 | 0.00 | 0.00 |
| URC2 | 0.89 | 0.11 | 0.00 | 0.00 | 0.00 | 0.00 | 0.00 | 0.02 | 0.12 | 0.25 | 0.30 | 0.30 | 0.01 | 0.00 | 0.02 | 0.09 | 0.25 | 0.30 | 0.31 | 0.03 | 0.00 |
| VHR1 | 0.46 | 0.37 | 0.15 | 0.02 | 0.00 | 0.00 | 0.00 | 0.00 | 0.04 | 0.12 | 0.23 | 0.40 | 0.22 | 0.00 | 0.00 | 0.01 | 0.05 | 0.14 | 0.46 | 0.33 | 0.00 |
| WAR1 | 0.07 | 0.33 | 0.29 | 0.18 | 0.12 | 0.00 | 0.00 | 0.00 | 0.03 | 0.05 | 0.08 | 0.31 | 0.51 | 0.01 | 0.00 | 0.00 | 0.00 | 0.02 | 0.11 | 0.77 | 0.09 |
| WTM1 | 0.96 | 0.04 | 0.00 | 0.00 | 0.00 | 0.00 | 0.00 | 0.00 | 0.00 | 0.01 | 0.03 | 0.22 | 0.71 | 0.04 | 0.00 | 0.00 | 0.01 | 0.03 | 0.21 | 0.72 | 0.04 |
| XBP1 | 0.55 | 0.38 | 0.07 | 0.00 | 0.00 | 0.00 | 0.00 | 0.00 | 0.23 | 0.32 | 0.28 | 0.15 | 0.01 | 0.00 | 0.00 | 0.06 | 0.31 | 0.36 | 0.25 | 0.02 | 0.00 |
| YAP1 | 1.00 | 0.00 | 0.00 | 0.00 | 0.00 | 0.00 | 0.00 | 0.00 | 0.00 | 0.04 | 0.54 | 0.42 | 0.00 | 0.00 | 0.00 | 0.00 | 0.04 | 0.54 | 0.42 | 0.00 | 0.00 |
| YAP3 | 1.00 | 0.00 | 0.00 | 0.00 | 0.00 | 0.00 | 0.00 | 0.00 | 0.00 | 0.00 | 0.46 | 0.54 | 0.00 | 0.00 | 0.00 | 0.00 | 0.00 | 0.46 | 0.54 | 0.00 | 0.00 |
| YAP5 | 1.00 | 0.00 | 0.00 | 0.00 | 0.00 | 0.00 | 0.00 | 0.00 | 0.00 | 0.11 | 0.15 | 0.45 | 0.30 | 0.00 | 0.00 | 0.00 | 0.11 | 0.15 | 0.45 | 0.30 | 0.00 |
| YAP6 | 1.00 | 0.00 | 0.00 | 0.00 | 0.00 | 0.00 | 0.00 | 0.00 | 0.00 | 0.00 | 0.47 | 0.53 | 0.00 | 0.00 | 0.00 | 0.00 | 0.00 | 0.47 | 0.53 | 0.00 | 0.00 |
| YAP7 | 1.00 | 0.00 | 0.00 | 0.00 | 0.00 | 0.00 | 0.00 | 0.00 | 0.00 | 0.30 | 0.40 | 0.30 | 0.00 | 0.00 | 0.00 | 0.00 | 0.30 | 0.40 | 0.30 | 0.00 | 0.00 |
| YDR266C | 0.78 | 0.22 | 0.00 | 0.00 | 0.00 | 0.00 | 0.00 | 0.03 | 0.25 | 0.22 | 0.14 | 0.35 | 0.00 | 0.00 | 0.02 | 0.18 | 0.27 | 0.15 | 0.38 | 0.00 | 0.00 |
| YDR520C | 0.89 | 0.11 | 0.00 | 0.00 | 0.00 | 0.00 | 0.00 | 0.02 | 0.12 | 0.26 | 0.30 | 0.29 | 0.01 | 0.00 | 0.02 | 0.10 | 0.25 | 0.30 | 0.31 | 0.03 | 0.00 |
| YER130C | 0.74 | 0.26 | 0.00 | 0.00 | 0.00 | 0.00 | 0.00 | 0.11 | 0.25 | 0.23 | 0.25 | 0.16 | 0.00 | 0.00 | 0.05 | 0.26 | 0.21 | 0.27 | 0.21 | 0.00 | 0.00 |
| YER184C | 0.87 | 0.12 | 0.01 | 0.00 | 0.00 | 0.00 | 0.00 | 0.00 | 0.35 | 0.44 | 0.17 | 0.03 | 0.00 | 0.00 | 0.00 | 0.27 | 0.47 | 0.23 | 0.03 | 0.00 | 0.00 |
| YGR067C | 0.49 | 0.37 | 0.13 | 0.01 | 0.00 | 0.00 | 0.00 | 0.13 | 0.27 | 0.28 | 0.21 | 0.11 | 0.00 | 0.00 | 0.04 | 0.16 | 0.27 | 0.30 | 0.22 | 0.01 | 0.00 |
| YHP1 | 0.95 | 0.05 | 0.00 | 0.00 | 0.00 | 0.00 | 0.00 | 0.00 | 0.00 | 0.10 | 0.36 | 0.54 | 0.00 | 0.00 | 0.00 | 0.00 | 0.10 | 0.34 | 0.57 | 0.00 | 0.00 |
| YKL222C | 0.41 | 0.39 | 0.15 | 0.05 | 0.01 | 0.00 | 0.00 | 0.01 | 0.04 | 0.09 | 0.12 | 0.29 | 0.43 | 0.02 | 0.00 | 0.01 | 0.05 | 0.09 | 0.28 | 0.53 | 0.05 |
| YLL054C | 0.00 | 0.75 | 0.25 | 0.00 | 0.00 | 0.00 | 0.00 | 0.28 | 0.45 | 0.19 | 0.08 | 0.00 | 0.00 | 0.00 | 0.00 | 0.19 | 0.42 | 0.28 | 0.11 | 0.00 | 0.00 |
| YLR278C | 0.82 | 0.18 | 0.00 | 0.00 | 0.00 | 0.00 | 0.00 | 0.00 | 0.25 | 0.47 | 0.25 | 0.03 | 0.00 | 0.00 | 0.00 | 0.18 | 0.44 | 0.31 | 0.06 | 0.00 | 0.00 |
| YML081W | 0.25 | 0.45 | 0.24 | 0.05 | 0.01 | 0.00 | 0.00 | 0.01 | 0.04 | 0.09 | 0.16 | 0.35 | 0.35 | 0.00 | 0.00 | 0.00 | 0.04 | 0.08 | 0.31 | 0.55 | 0.01 |
| YNR063W | 0.60 | 0.32 | 0.07 | 0.01 | 0.00 | 0.00 | 0.00 | 0.00 | 0.01 | 0.06 | 0.11 | 0.40 | 0.42 | 0.00 | 0.00 | 0.00 | 0.03 | 0.09 | 0.35 | 0.53 | 0.00 |
| YOX1 | 0.99 | 0.01 | 0.00 | 0.00 | 0.00 | 0.00 | 0.00 | 0.00 | 0.00 | 0.00 | 0.18 | 0.74 | 0.08 | 0.00 | 0.00 | 0.00 | 0.00 | 0.18 | 0.74 | 0.08 | 0.00 |
| YPR013C | 0.71 | 0.24 | 0.04 | 0.00 | 0.00 | 0.00 | 0.00 | 0.00 | 0.00 | 0.01 | 0.05 | 0.30 | 0.63 | 0.01 | 0.00 | 0.00 | 0.00 | 0.02 | 0.26 | 0.70 | 0.02 |
| YPR015C | 0.63 | 0.29 | 0.07 | 0.01 | 0.00 | 0.00 | 0.00 | 0.00 | 0.00 | 0.01 | 0.05 | 0.29 | 0.64 | 0.02 | 0.00 | 0.00 | 0.00 | 0.02 | 0.22 | 0.73 | 0.02 |
| YPR022C | 0.74 | 0.25 | 0.02 | 0.00 | 0.00 | 0.00 | 0.00 | 0.26 | 0.44 | 0.26 | 0.04 | 0.00 | 0.00 | 0.00 | 0.17 | 0.41 | 0.32 | 0.10 | 0.00 | 0.00 | 0.00 |
| YPR196W | 0.60 | 0.33 | 0.07 | 0.01 | 0.00 | 0.00 | 0.00 | 0.00 | 0.01 | 0.03 | 0.08 | 0.27 | 0.57 | 0.03 | 0.00 | 0.01 | 0.02 | 0.06 | 0.23 | 0.64 | 0.04 |
| YRM1 | 0.41 | 0.38 | 0.16 | 0.03 | 0.01 | 0.00 | 0.00 | 0.01 | 0.05 | 0.09 | 0.12 | 0.25 | 0.43 | 0.04 | 0.00 | 0.03 | 0.05 | 0.10 | 0.26 | 0.50 | 0.07 |
| YRR1 | 0.78 | 0.21 | 0.01 | 0.00 | 0.00 | 0.00 | 0.00 | 0.01 | 0.19 | 0.25 | 0.26 | 0.27 | 0.02 | 0.00 | 0.01 | 0.12 | 0.25 | 0.29 | 0.32 | 0.02 | 0.00 |
| ZAP1 | 0.87 | 0.13 | 0.00 | 0.00 | 0.00 | 0.00 | 0.00 | 0.11 | 0.21 | 0.33 | 0.23 | 0.12 | 0.00 | 0.00 | 0.09 | 0.19 | 0.32 | 0.26 | 0.14 | 0.00 | 0.00 |

**Supplementary Figure 34** ***(preceding page)*****: The impact of the addition of m6A to SMAC-seq on assay resolution and the potential ability to footprint individual transcription factors**. Shown is the fraction of motifs in the genome for each transcription factor in the yeast genome containing the indicated number of informative positions using GC alone, m6A alone, and GC + m6A methyltransferase.

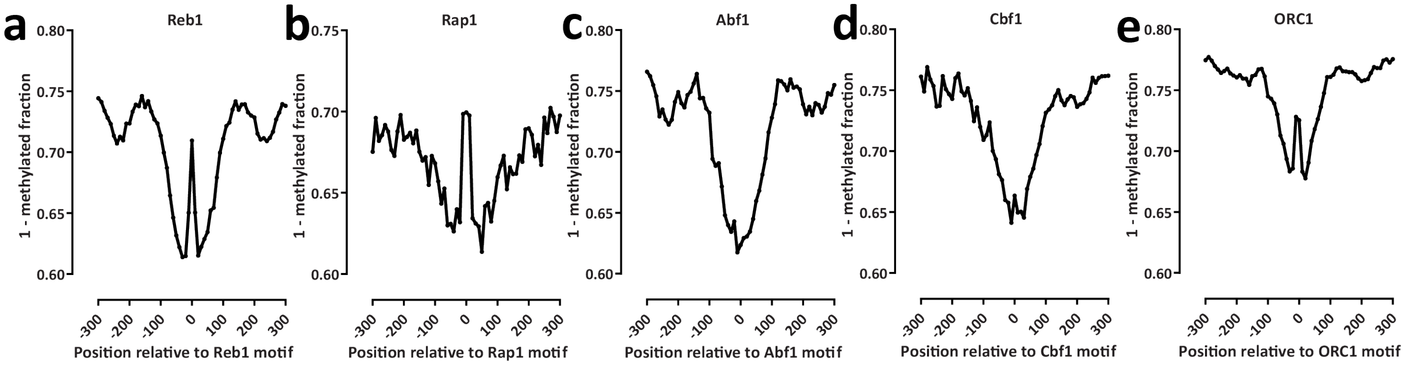


**Supplementary Figure 35: Single-molecule footprinting by transcription factors**. Shown is the average methylation status (averaged over 10bp) in the neighborhood of occupied (as measured by ChIP-exo or ChIP-seq) recognition motifs for several *S. cerevisiae* DNA binding proteins: (a) Reb1; (b) Rap1; (c) Abf1; (d) Cbf1. (e) ORC1. Strong footprinting is observed for Reb1, Rap1, and ORC1, while Abf1 and Cbf1 occupancy does not appear to be strongly protective against methylation.

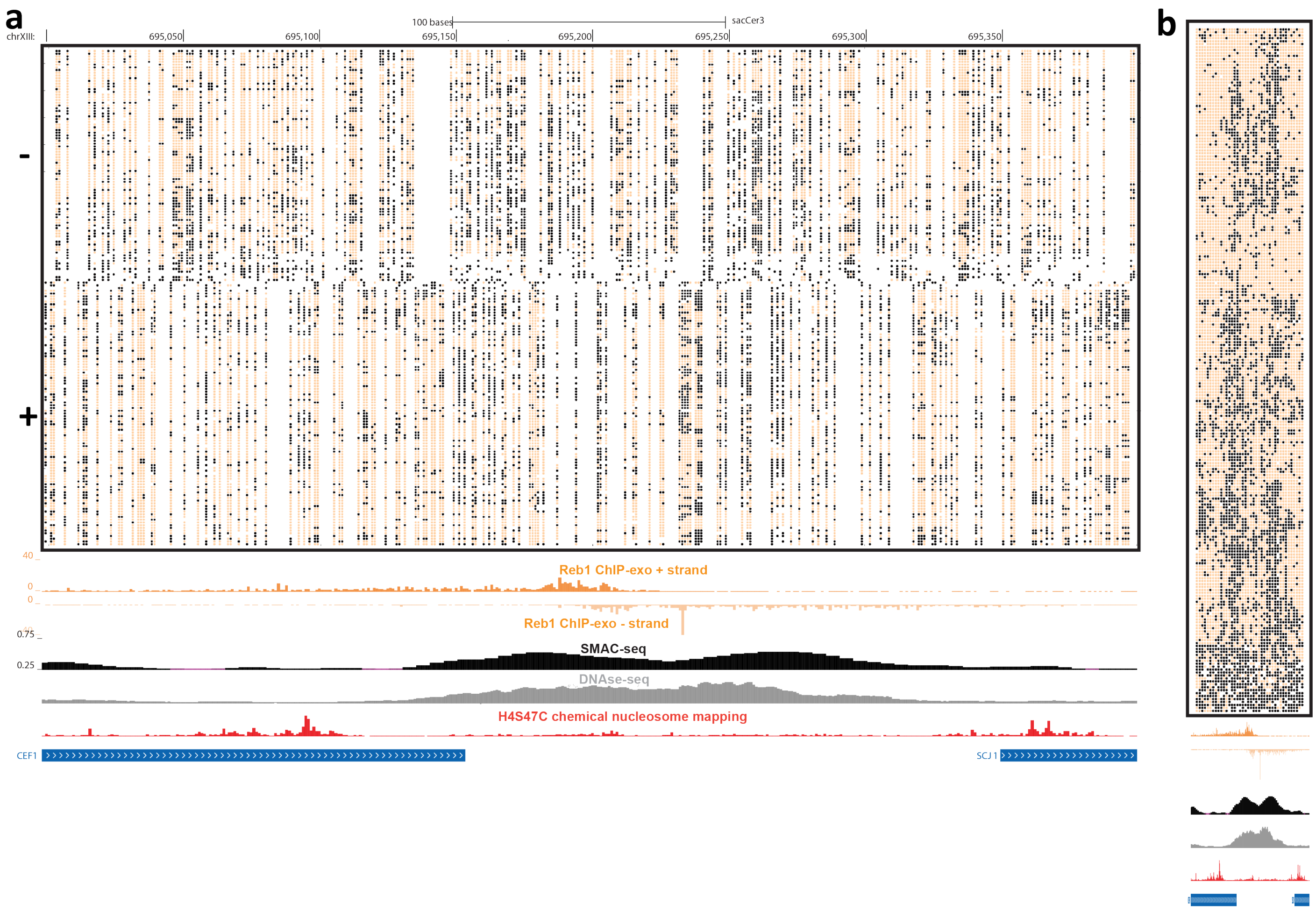


**Supplementary Figure 36: Single-molecule footprinting by Reb1 binding sites**. (a) Raw unfiltered nanopore reads fully spanning the 400-bp neighborhood of a Reb1 binding site on chrXVIII, at single-bp resolution; (b) Same as in (a), but at aggregated 10-bp resolution.

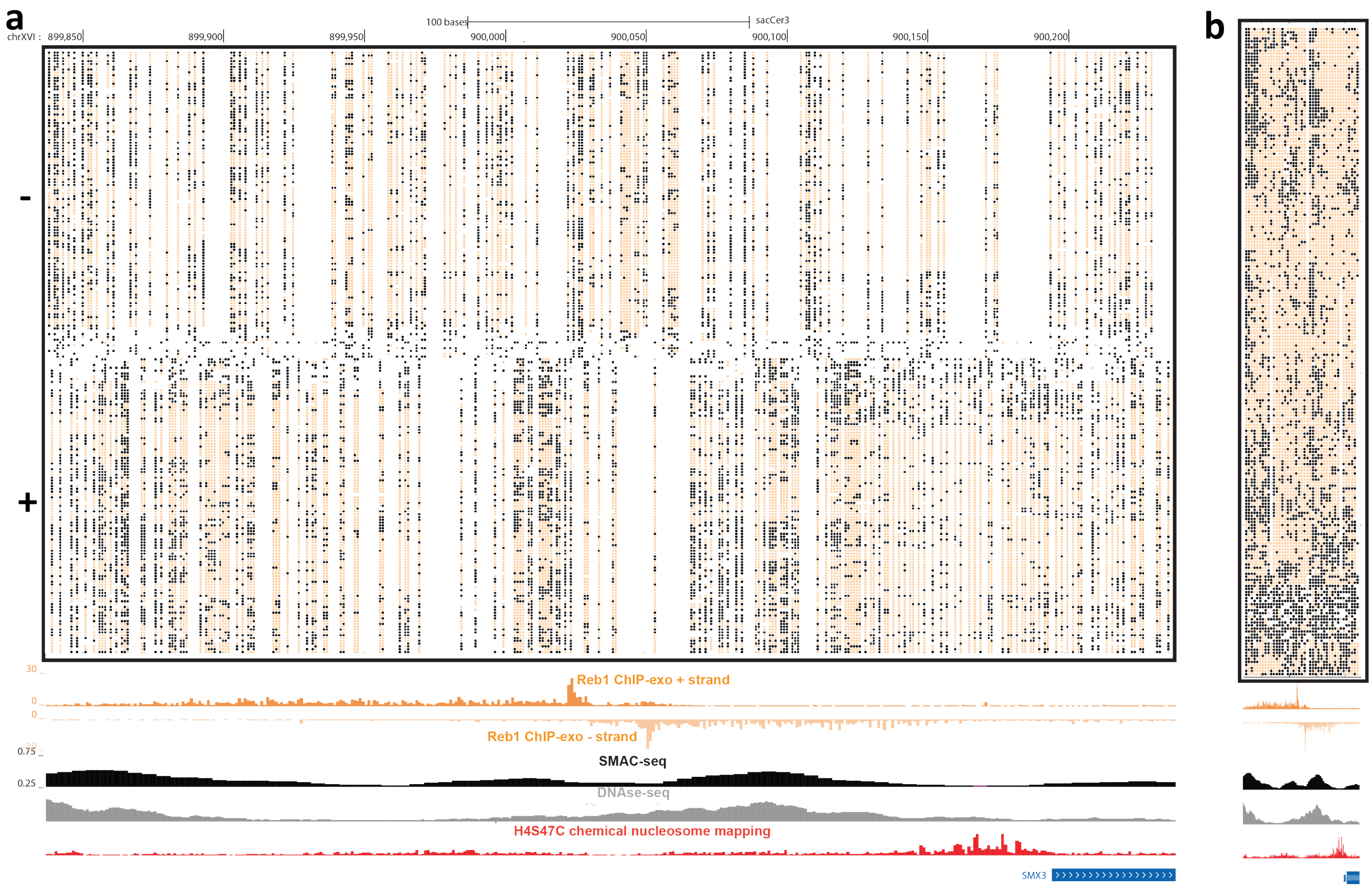


**Supplementary Figure 37: Single-molecule footprinting by Reb1 binding sites**. (a) Raw unfiltered nanopore reads fully spanning the 400-bp neighborhood of a Reb1 binding site on chrXVIII, at single-bp resolution; (b) Same as in (a), but at aggregated 10-bp resolution.

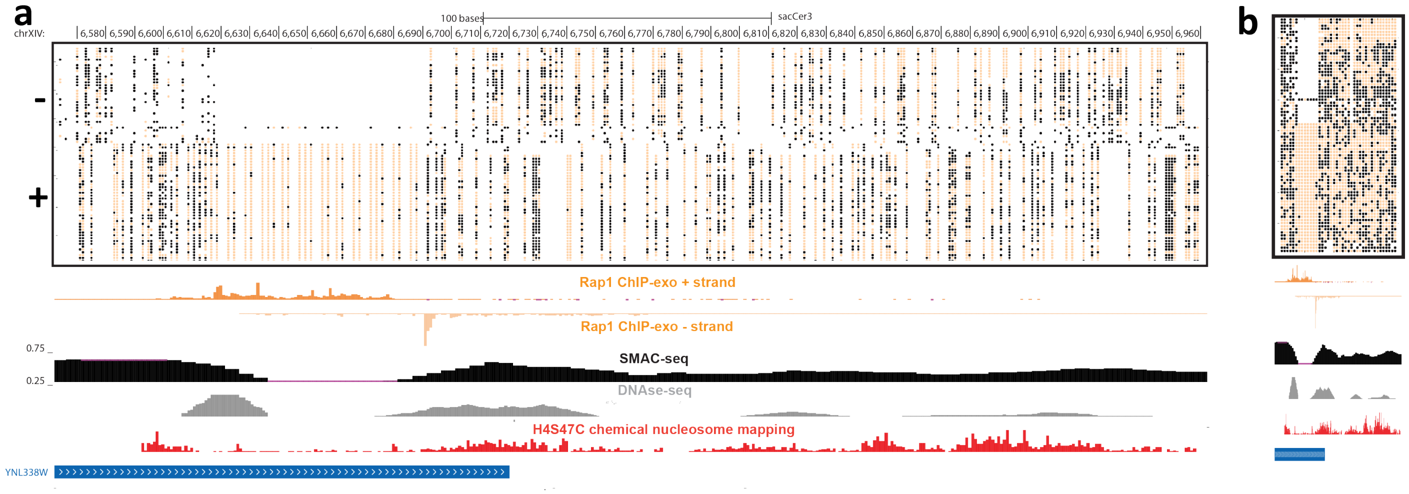


**Supplementary Figure 38: Single-molecule footprinting associated with Rap1 occupancy**. (a) Raw unfiltered nanopore reads fully spanning a 400-bp neighborhood of the subtelomeric region of chrXIV, at single-bp resolution; (b) Same as in (a), but at aggregated 10-bp resolution.

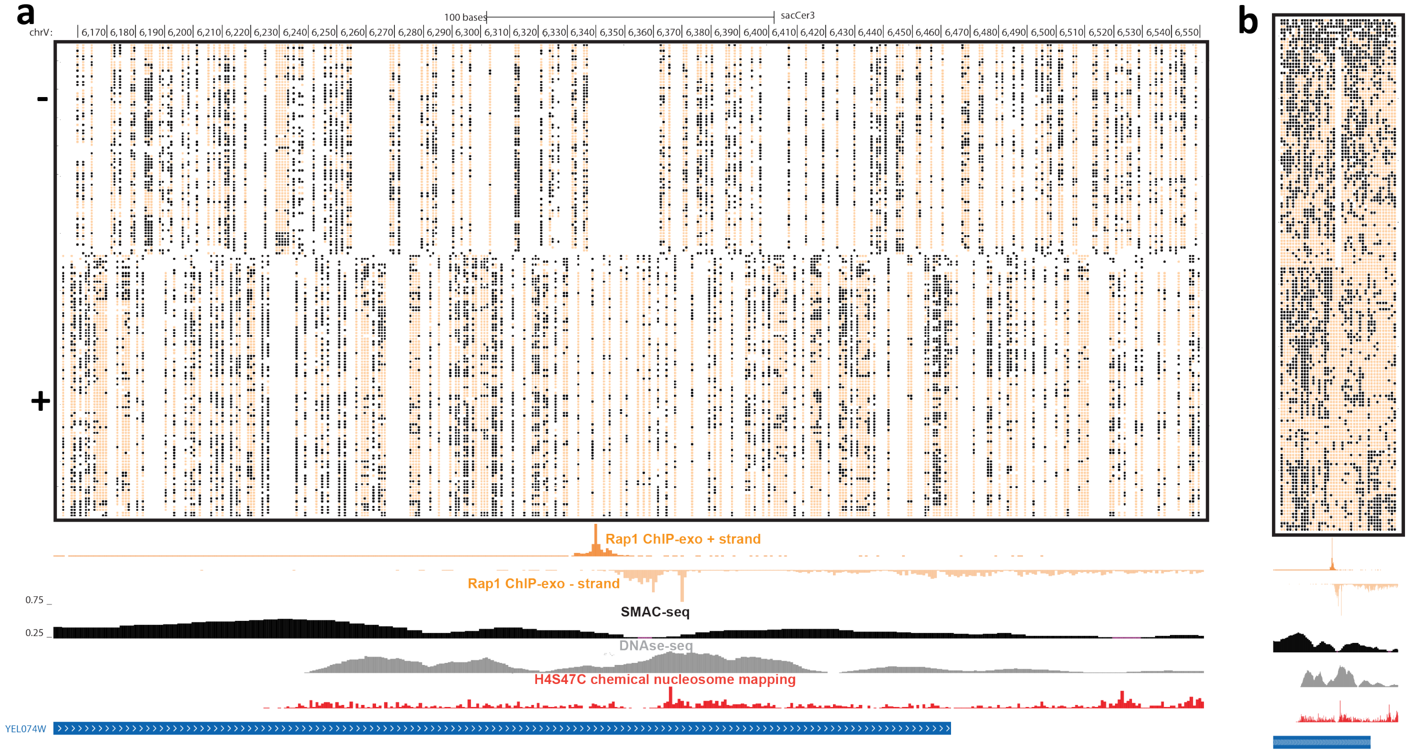


**Supplementary Figure 39: Single-molecule footprinting associated with Rap1 occupancy**. (a) Raw unfiltered nanopore reads fully spanning a 400-bp neighborhood of the subtelomeric region of chrXV, at single-bp resolution; (b) Same as in (a), but at aggregated 10-bp resolution.

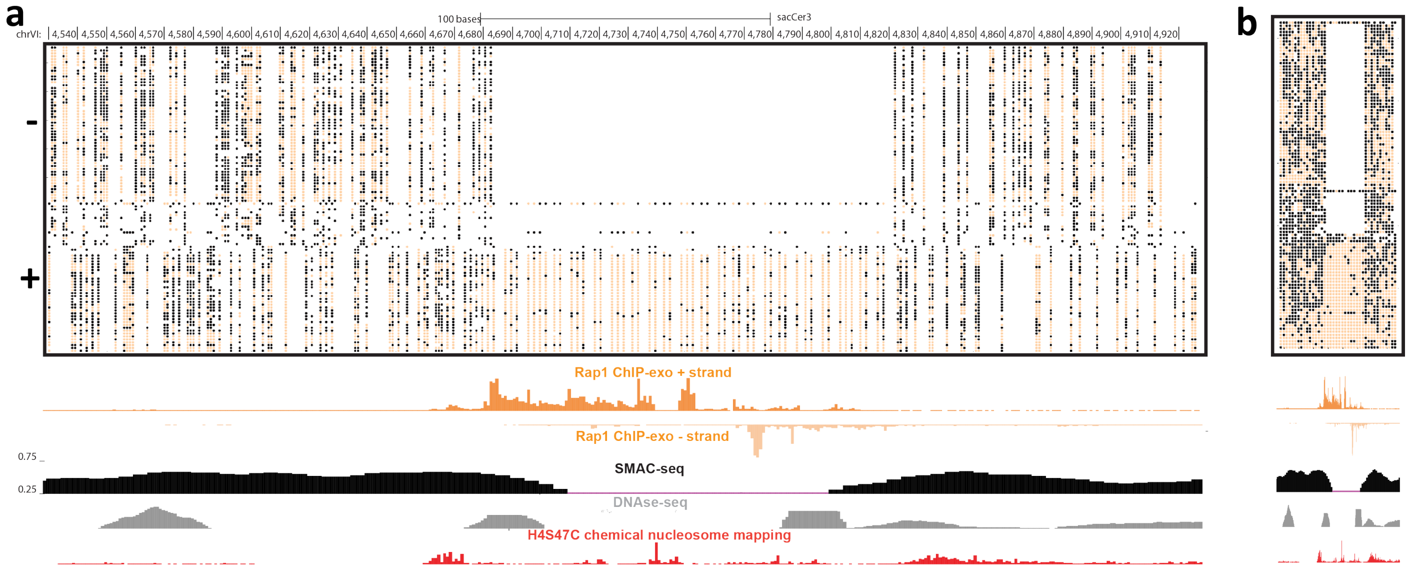


**Supplementary Figure 40: Single-molecule footprinting associated with Rap1 occupancy**. (a) Raw unfiltered nanopore reads fully spanning a 400-bp neighborhood of the subtelomeric region of chrVI, at single-bp resolution; (b) Same as in (a), but at aggregated 10-bp resolution.

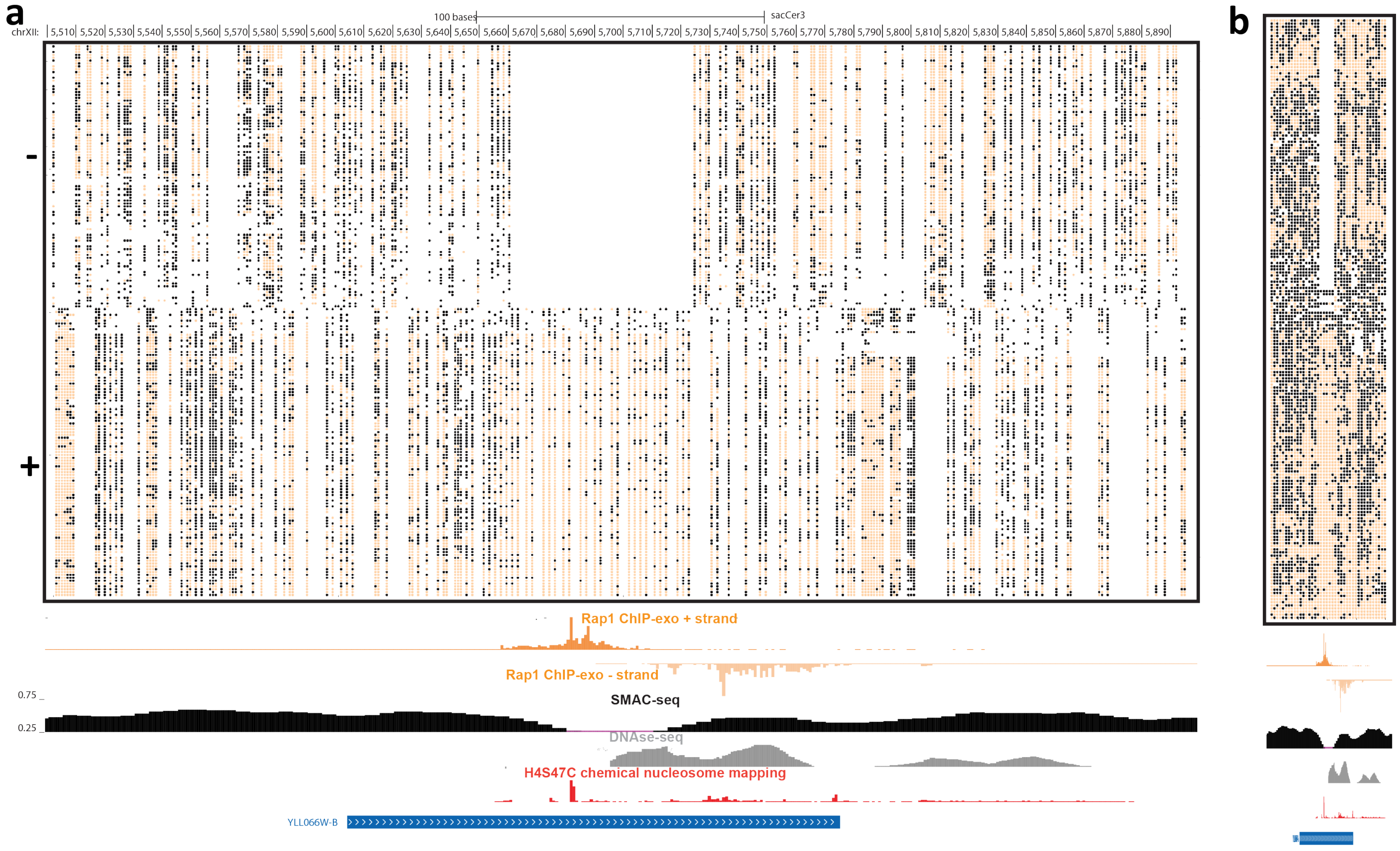


**Supplementary Figure 41: Single-molecule footprinting associated with Rap1 occupancy**. (a) Raw unfiltered nanopore reads fully spanning a 400-bp neighborhood of the subtelomeric region of chrXII, at single-bp resolution; (b) Same as in (a), but at aggregated 10-bp resolution.

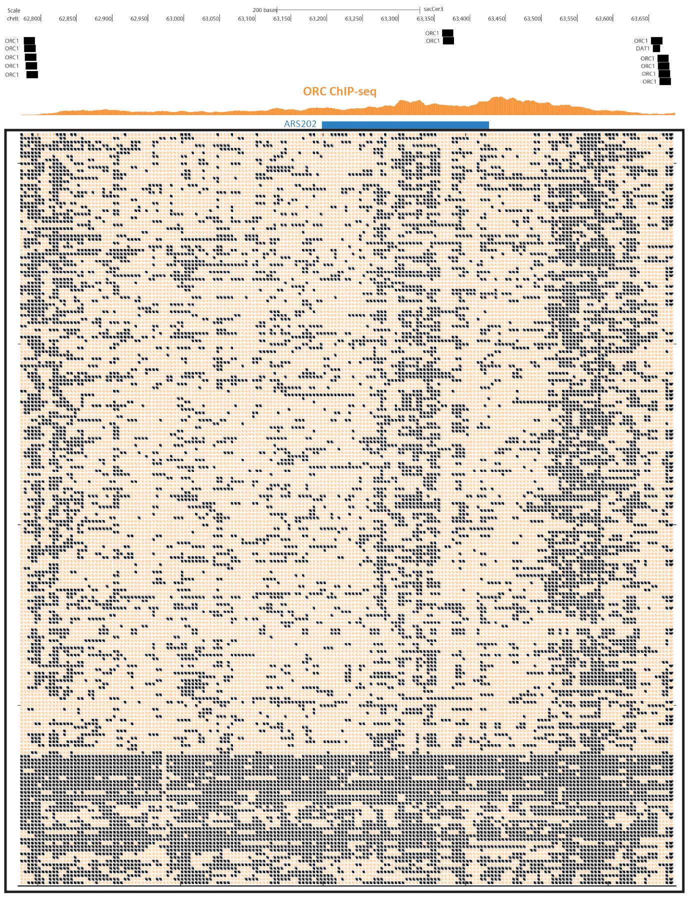


**Supplementary Figure 42: Single-molecule footprinting associated with ORC occupancy**. (a) Raw unfiltered nanopore reads fully spanning the neighborhood of an ARS site on chrII, at 5-bp aggregated resolution;

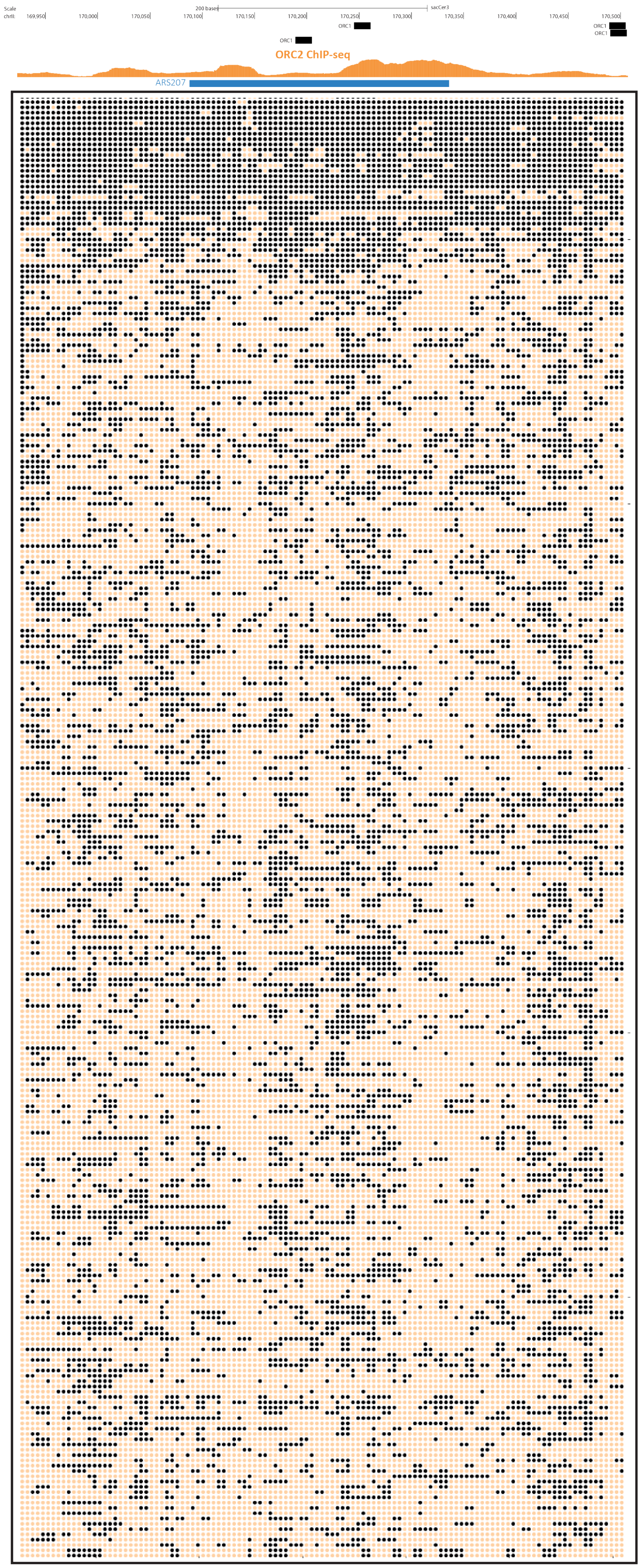


**Supplementary Figure 43: Single-molecule footprinting associated with ORC occupancy**. (a) Raw unfiltered nanopore reads fully spanning the neighborhood of an ARS site on chrII, at 5-bp aggregated resolution;

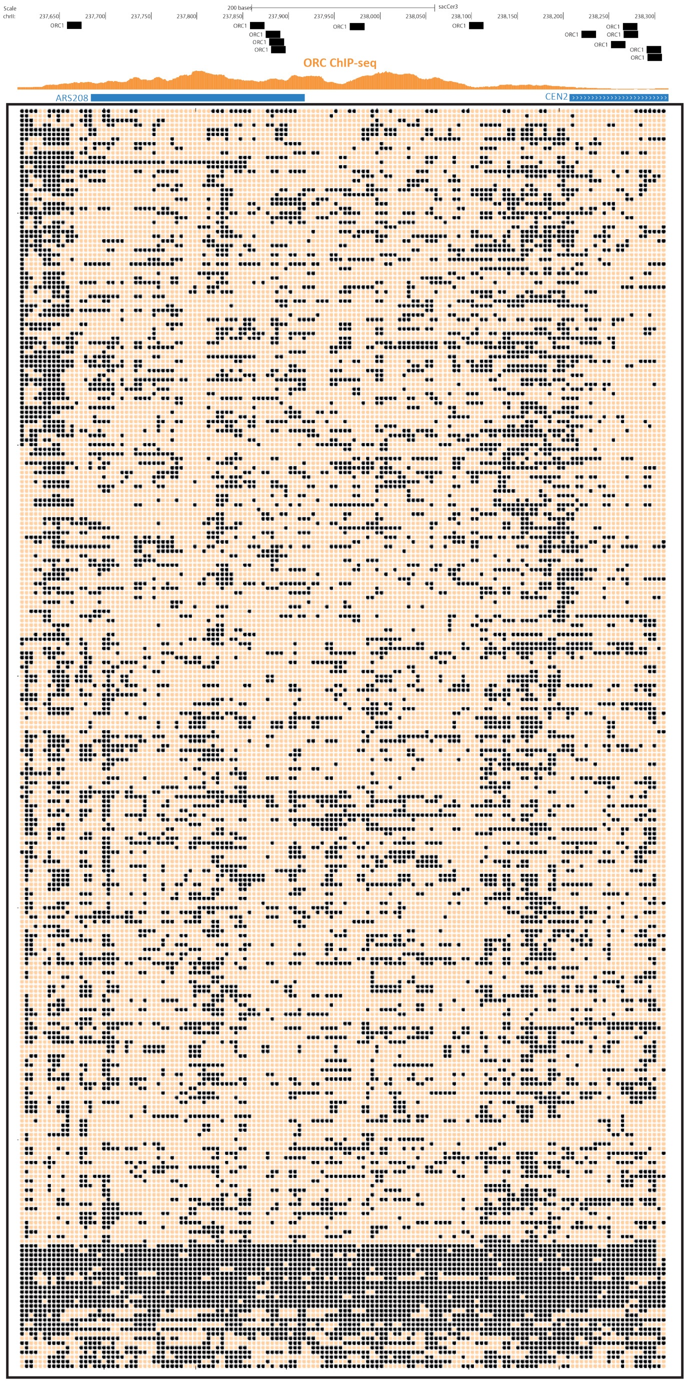


**Supplementary Figure 44: Single-molecule footprinting associated with ORC occupancy**. (a) Raw unfiltered nanopore reads fully spanning the neighborhood of an ARS site on chrII, at 5-bp aggregated resolution;

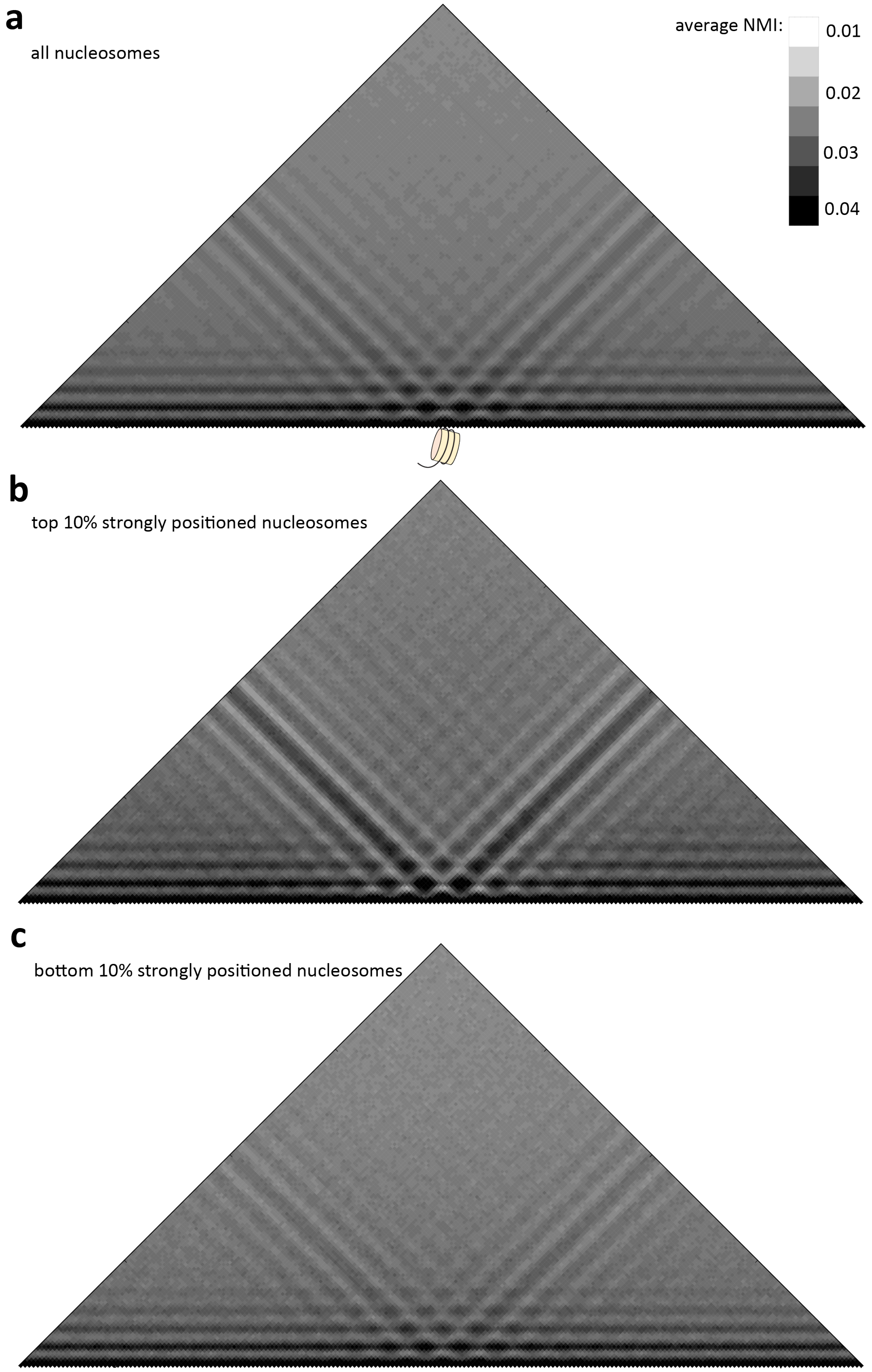


**Supplementary Figure 45: Metanucleosome NMI profiles in the yeast genome**. Shown are average NMI maps between all 20-bp segments centered on each positioned nucleosome in the genome (a), the top 10% strongly positioned nucleosomes (b), or the top 10% strongly positioned nucleosomes (c).

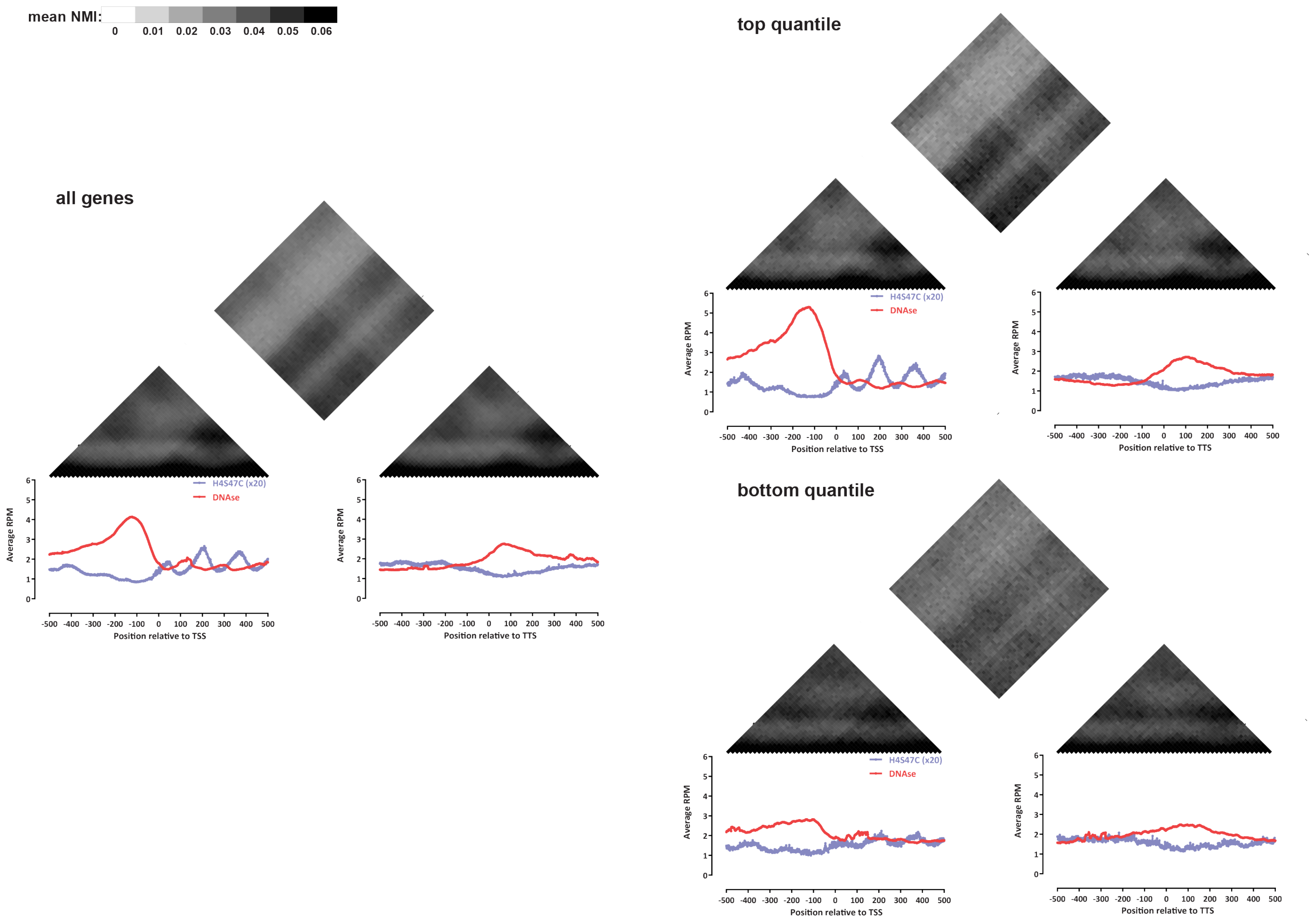


**Supplementary Figure 46: Patterns of coaccessibility between the 5′ and 3′ ends of genes**. Shown is the average NMI for the ±500bp regions in the 5′ and 3′ end of all yeast genes as well as the top and bottom 20% expression-ranked genes (calculated over 10-bp windows). Only genes ≥1000 bp in length are shown. Similar results are obtained using windows of size 20bp or 50bp (data not shown).

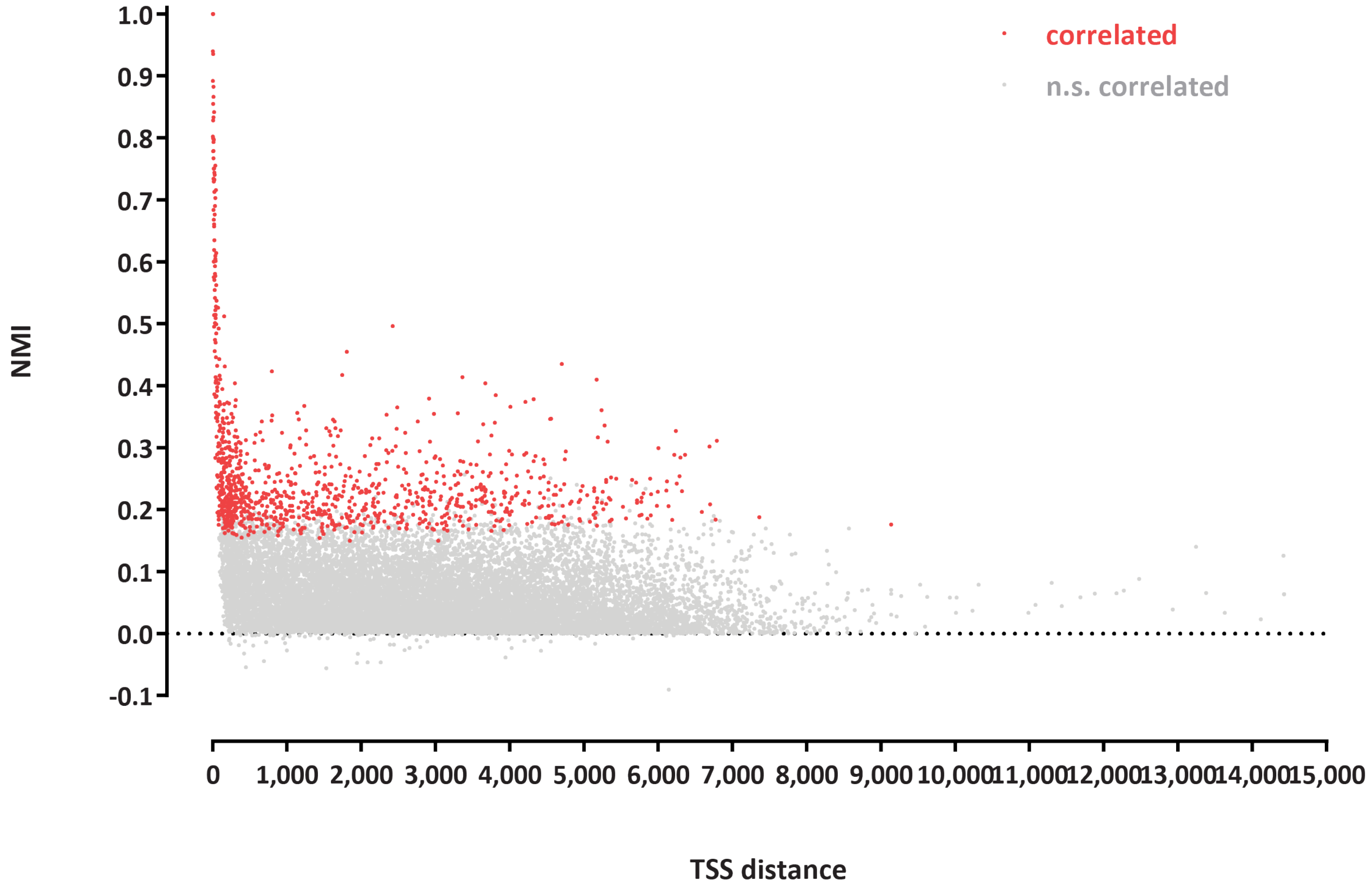


**Supplementary Figure 47: Accessibility correlation between TSSs in the yeast genome**. Shown are NMI values for each pair of significantly and non-significantly correlated TSSs (defined as the regions ±100 bp around the TSS).

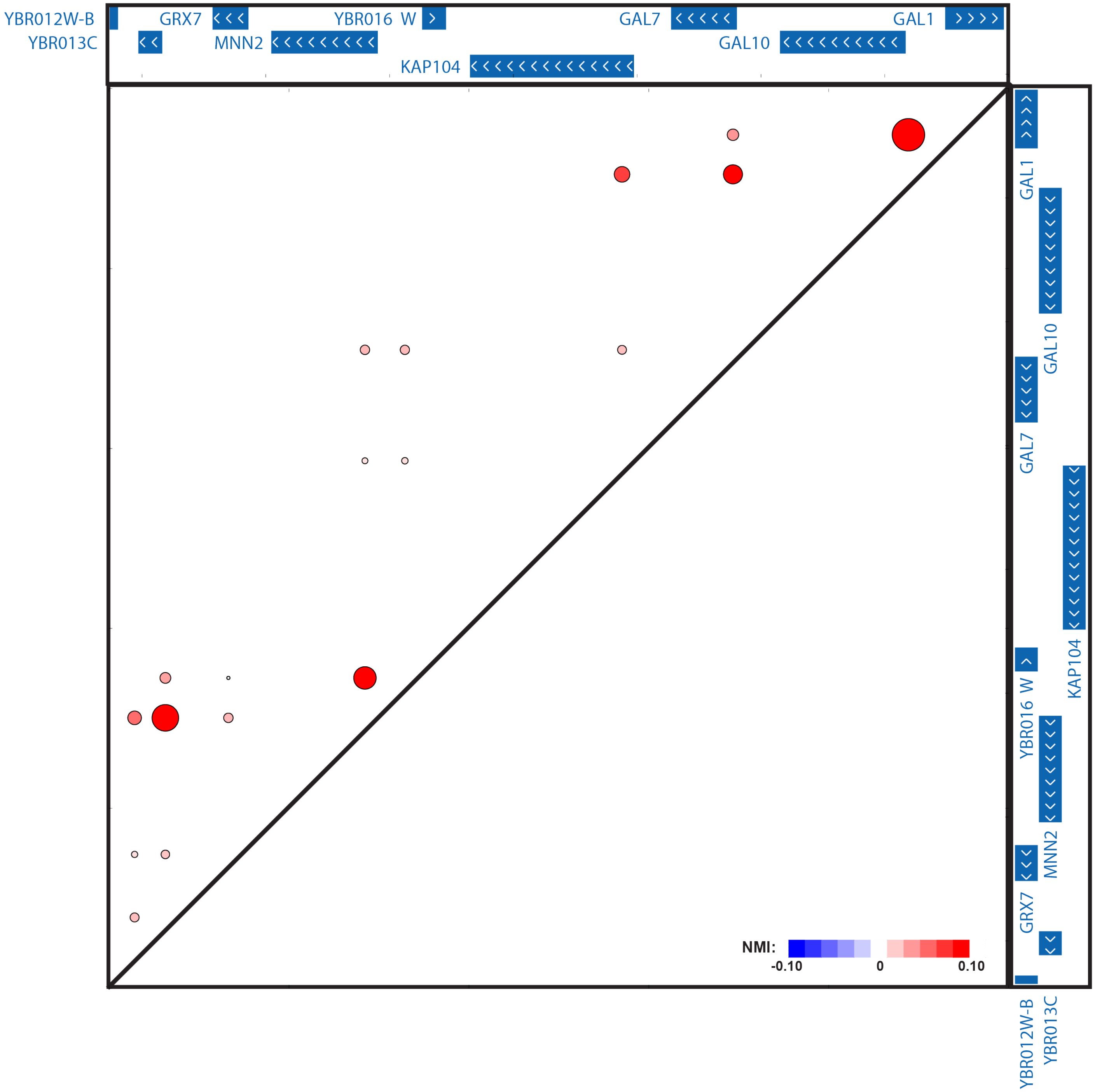


**Supplementary Figure 48: Example of accessibility correlation between TSSs**. Shown are accessibility correlations between all loci in the region with coordinates “chrII:265000-280000”. The size and color of the dots corresponds to their NMI scores.

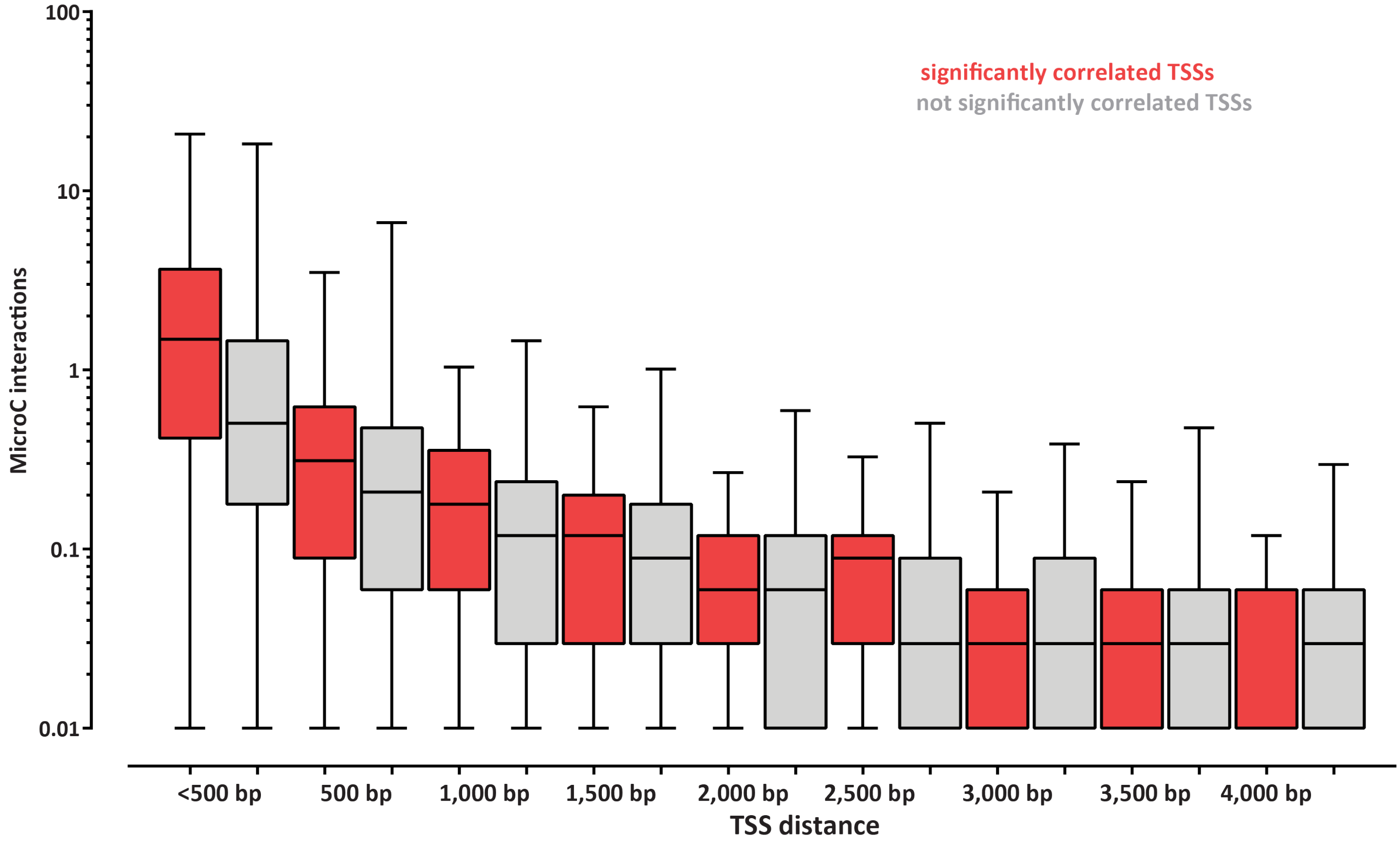


**Supplementary Figure 49: Accessibility correlation between TSSs and 3D interactions**. Shown are significantly and non-significantly correlated TSSs (defined as the regions ±100 bp around the TSS) split into distance bins and the number of 3D interactions between each group (measured by MicroC).

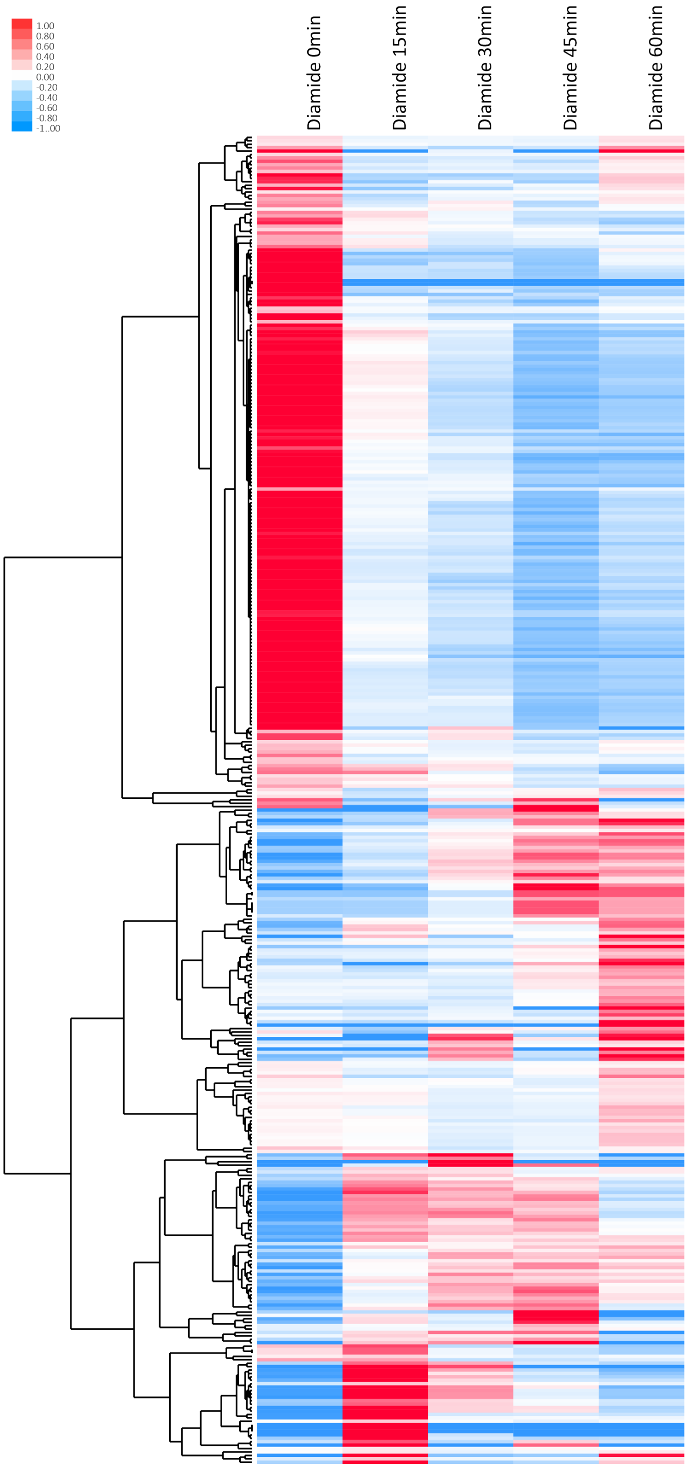


**Supplementary Figure 50: Gene expression changes upon diamide treatment**. Shown is RNA-seq data (mean and unit-variance normalized across time points) for all genes expressed at ≥50 FPKM in at least one time point.

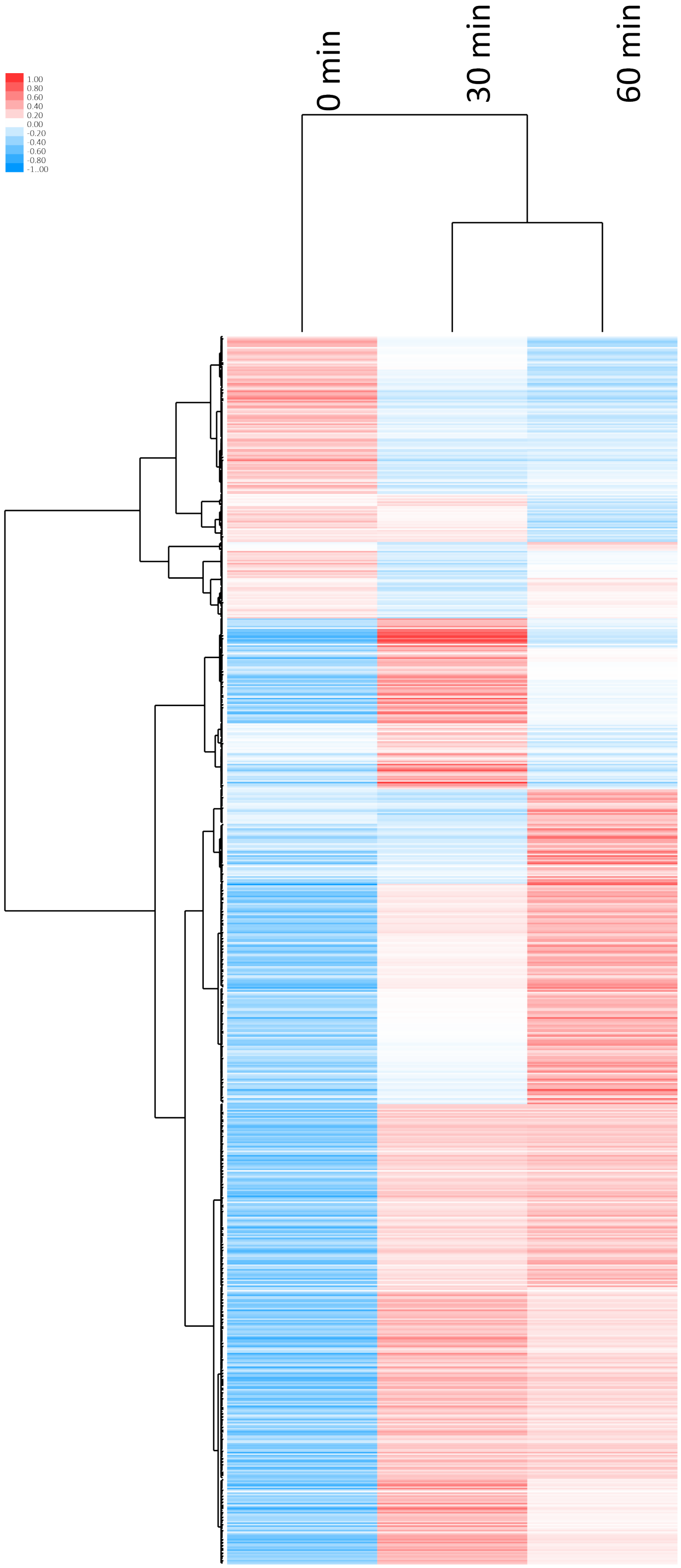


**Supplementary Figure 51: Dynamic changes in HSF1 occupancy upon diamide treatment**. Shown are ChIP-seq RPMs (mean and unit-variance normalized across time points) for all HSF1 peaks detected in least one time point.

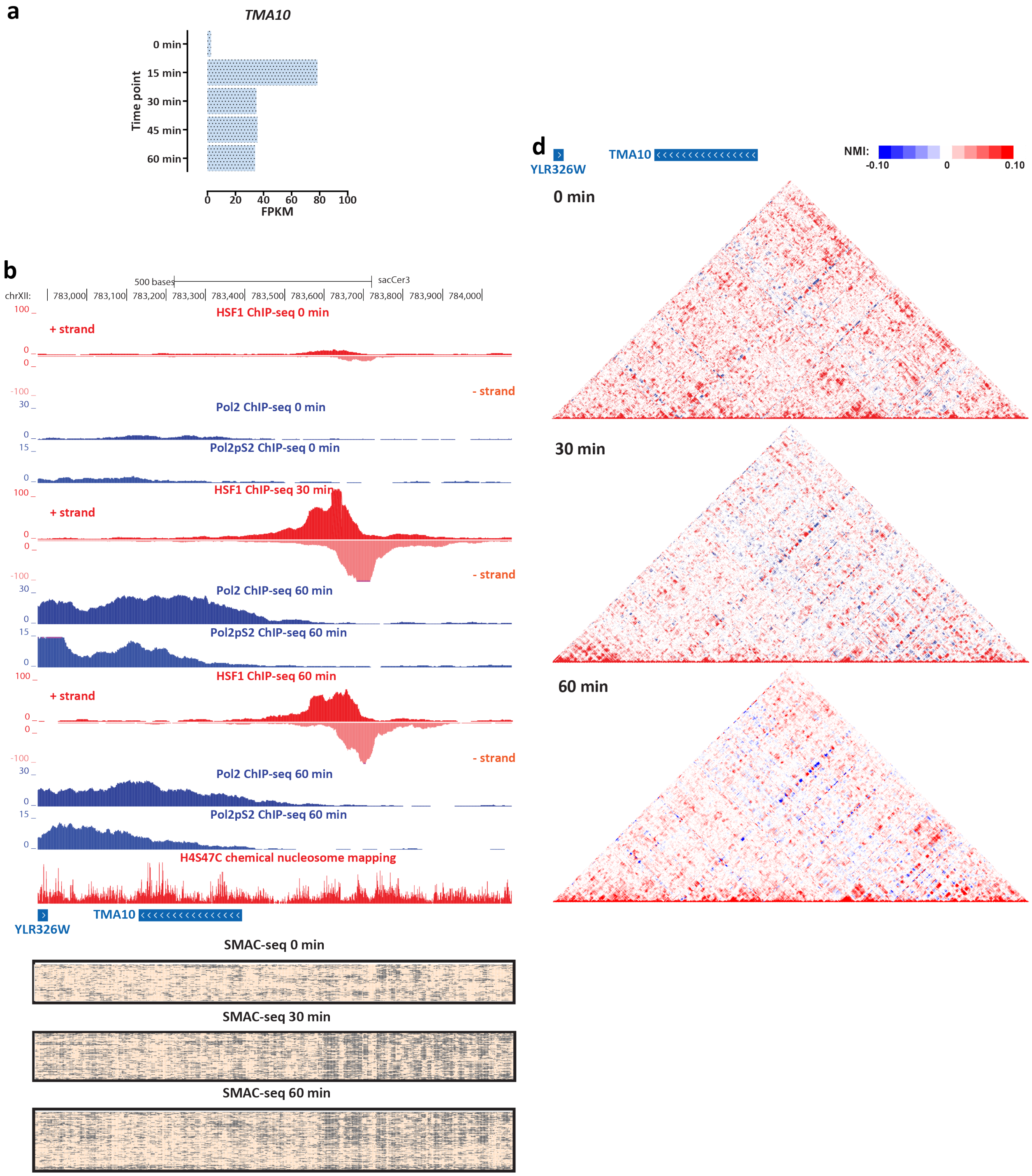


**Supplementary Figure 52: Coordinated changes in chromatin accessibility and nucleosomal occupancy during the yeast stress response**. Shown are changes in RNA Polymerase and HSF1 occupancy (measured by ChIP-seq), SMAC-seq profiles (1-bp resolution, 10-bp aggregate scores) and NMI profiles in the vicinity of the *TMA10* gene.

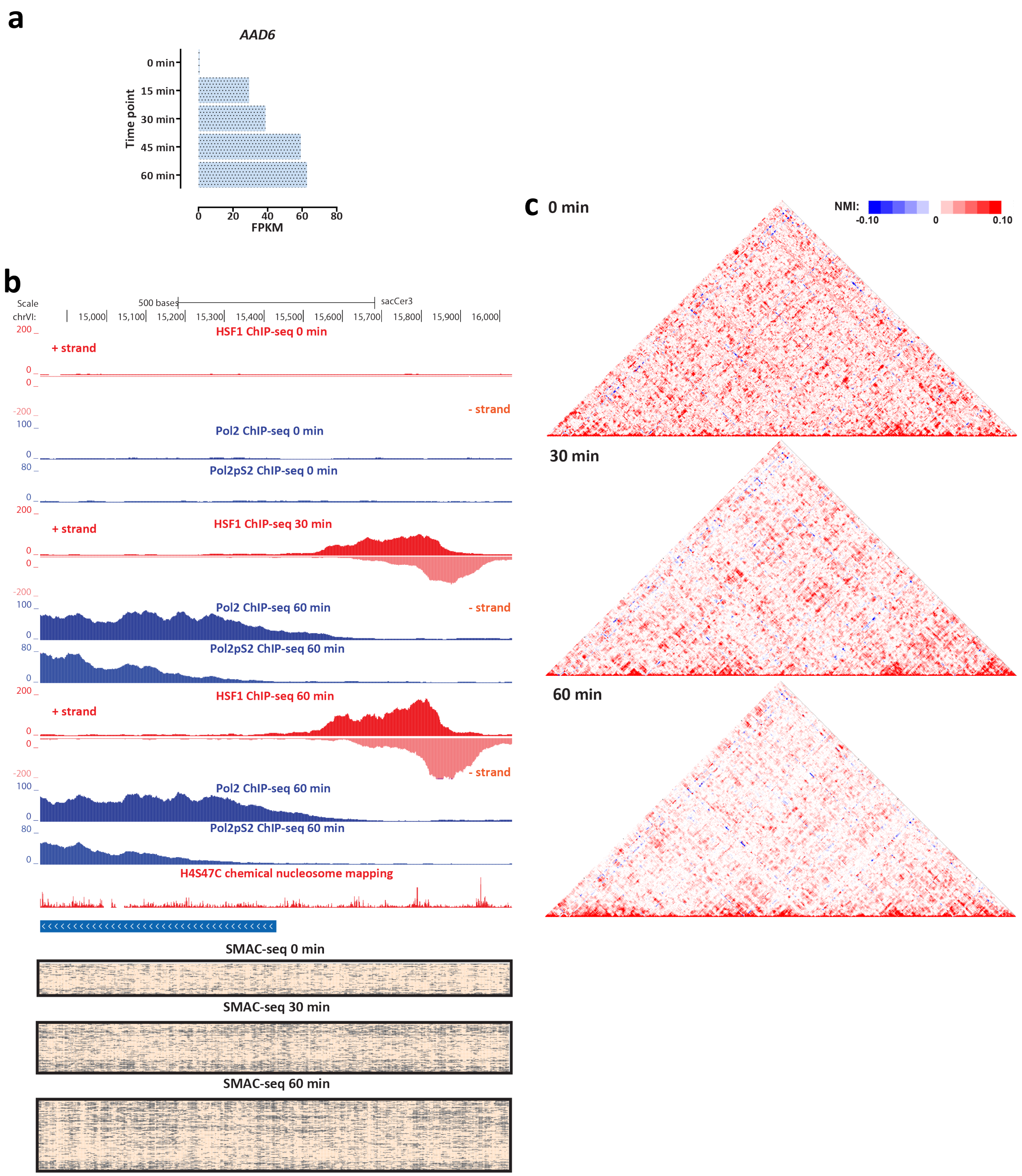


**Supplementary Figure 53: Coordinated changes in chromatin accessibility and nucleosomal occupancy during the yeast stress response**. Shown are changes in RNA Polymerase and HSF1 occupancy (measured by ChIP-seq), SMAC-seq profiles (1-bp resolution, 10-bp aggregate scores) and NMI profiles in the vicinity of the *AAD6* gene.

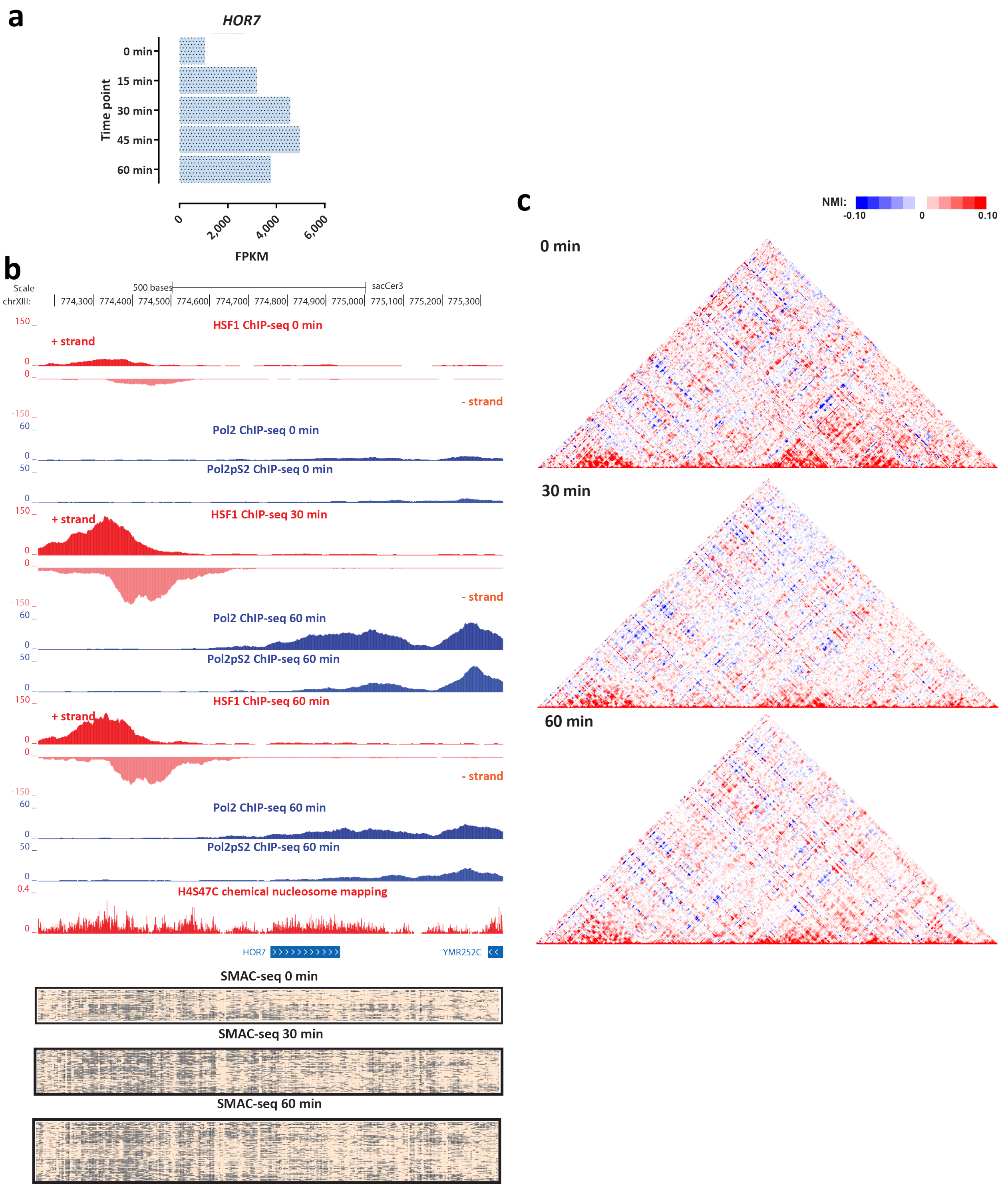


**Supplementary Figure 54: Coordinated changes in chromatin accessibility and nucleosomal occupancy during the yeast stress response**. Shown are changes in RNA Polymerase and HSF1 occupancy (measured by ChIP-seq), SMAC-seq profiles (1-bp resolution, 10-bp aggregate scores) and NMI profiles in the vicinity of the *HOR7* gene.

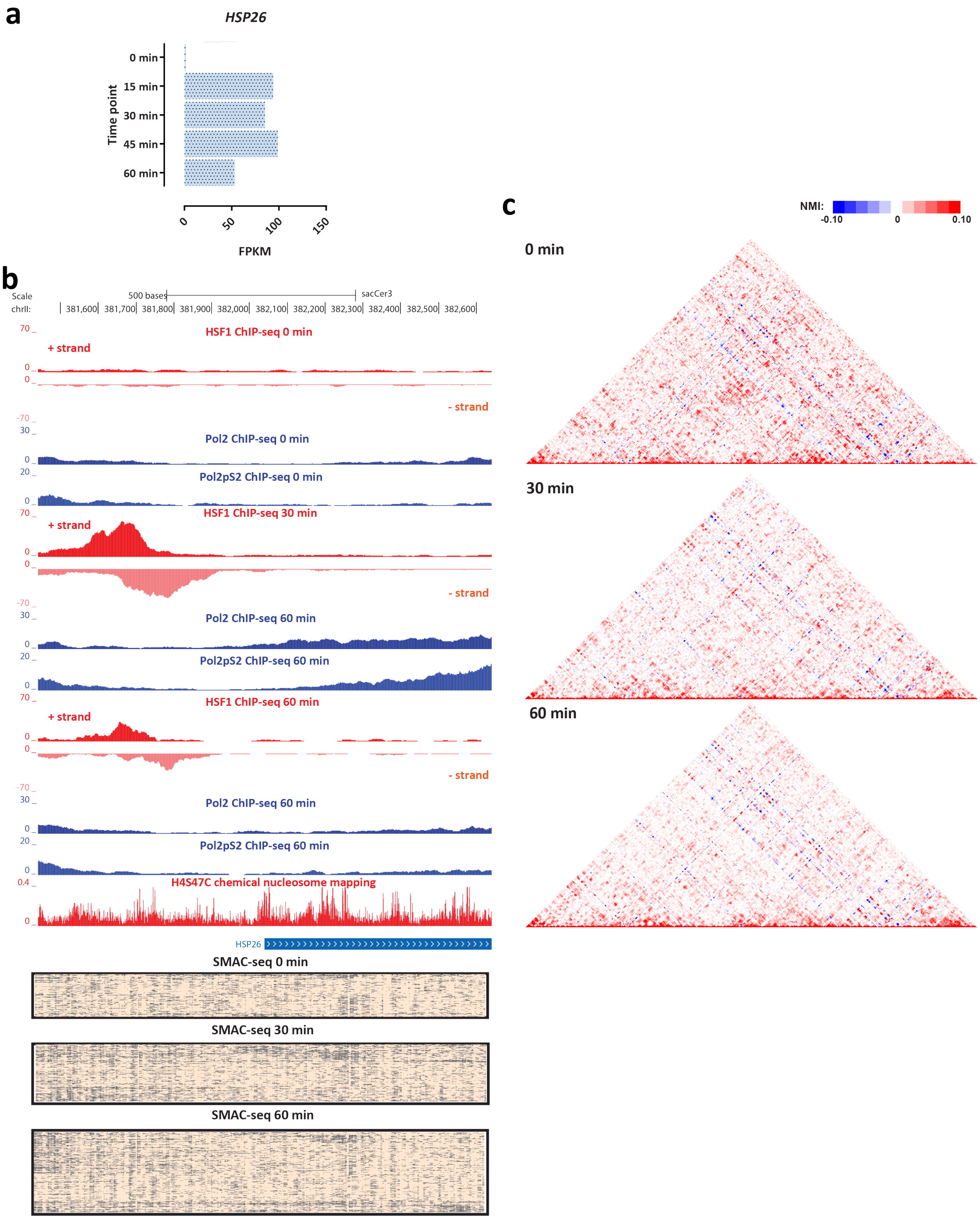


**Supplementary Figure 55: Coordinated changes in chromatin accessibility and nucleosomal occupancy during the yeast stress response**. Shown are changes in RNA Polymerase and HSF1 occupancy (measured by ChIP-seq), SMAC-seq profiles (1-bp resolution, 10-bp aggregate scores) and NMI profiles in the vicinity of the *HSP26* gene.

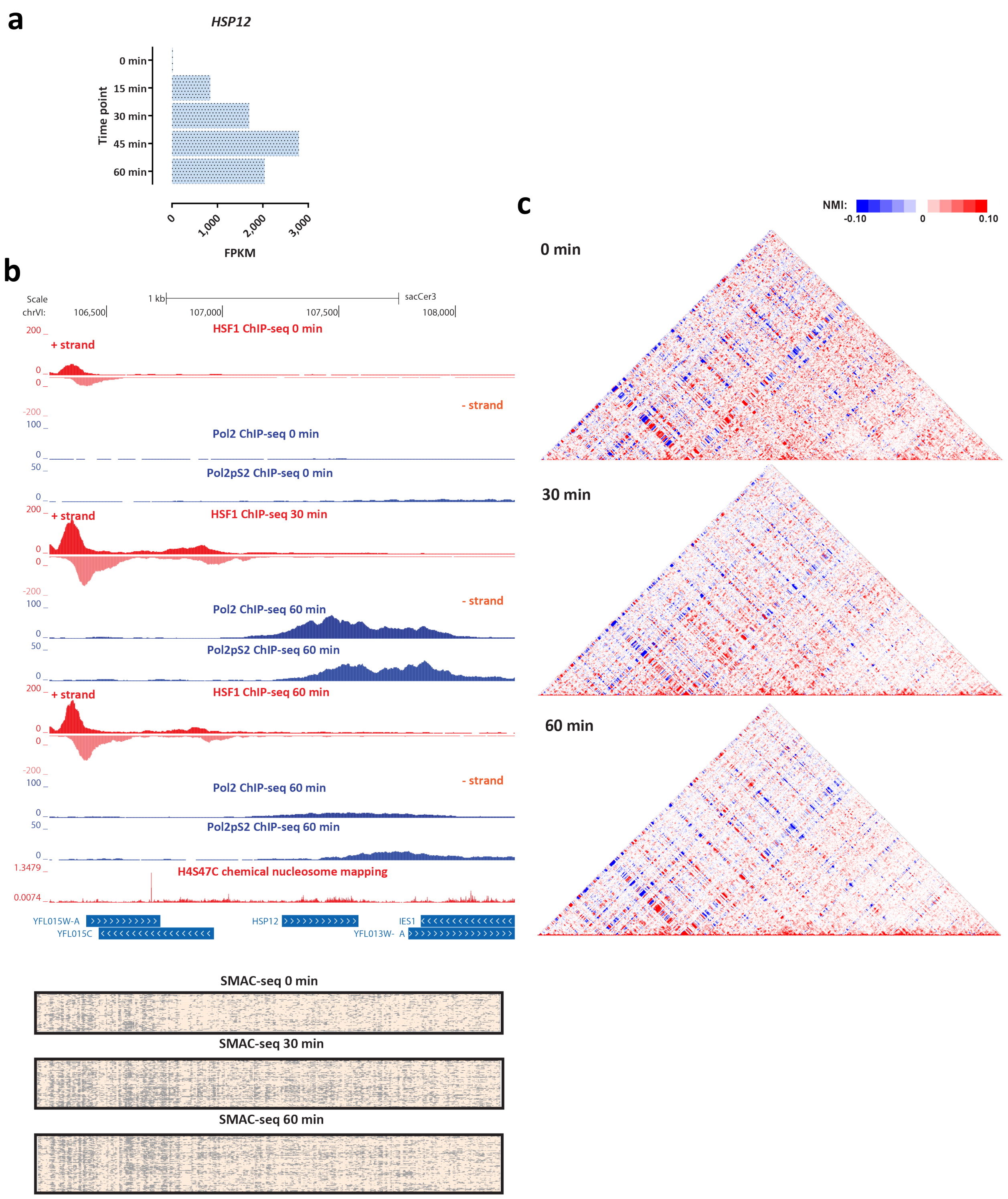


**Supplementary Figure 56: Coordinated changes in chromatin accessibility and nucleosomal occupancy during the yeast stress response**. Shown are changes in RNA Polymerase and HSF1 occupancy (measured by ChIP-seq), SMAC-seq profiles (1-bp resolution, 10-bp aggregate scores) and NMI profiles in the vicinity of the *HSP12* gene.

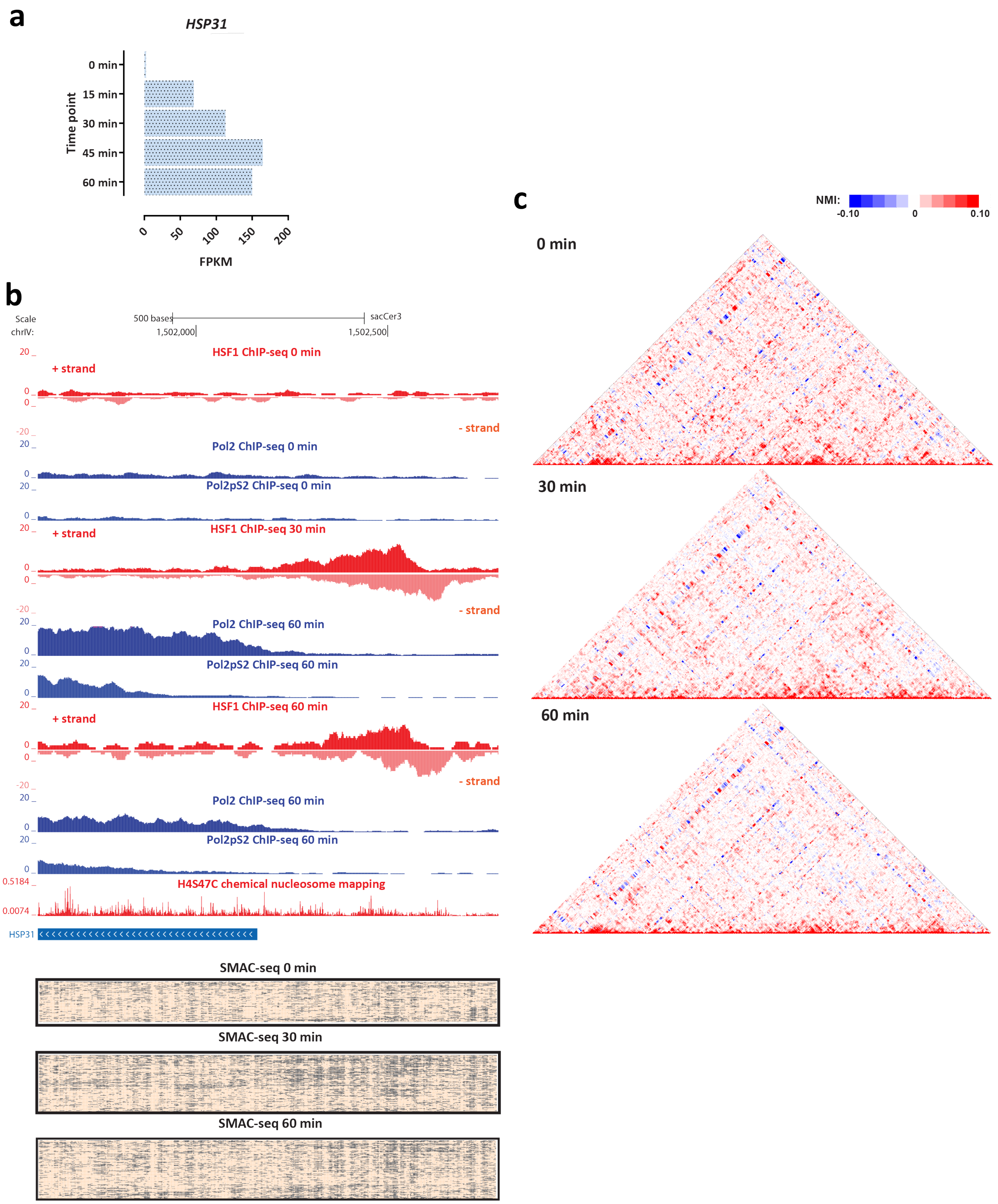


**Supplementary Figure 57: Coordinated changes in chromatin accessibility and nucleosomal occupancy during the yeast stress response**. Shown are changes in RNA Polymerase and HSF1 occupancy (measured by ChIP-seq), SMAC-seq profiles (1-bp resolution, 10-bp aggregate scores) and NMI profiles in the vicinity of the *HSP31* gene.

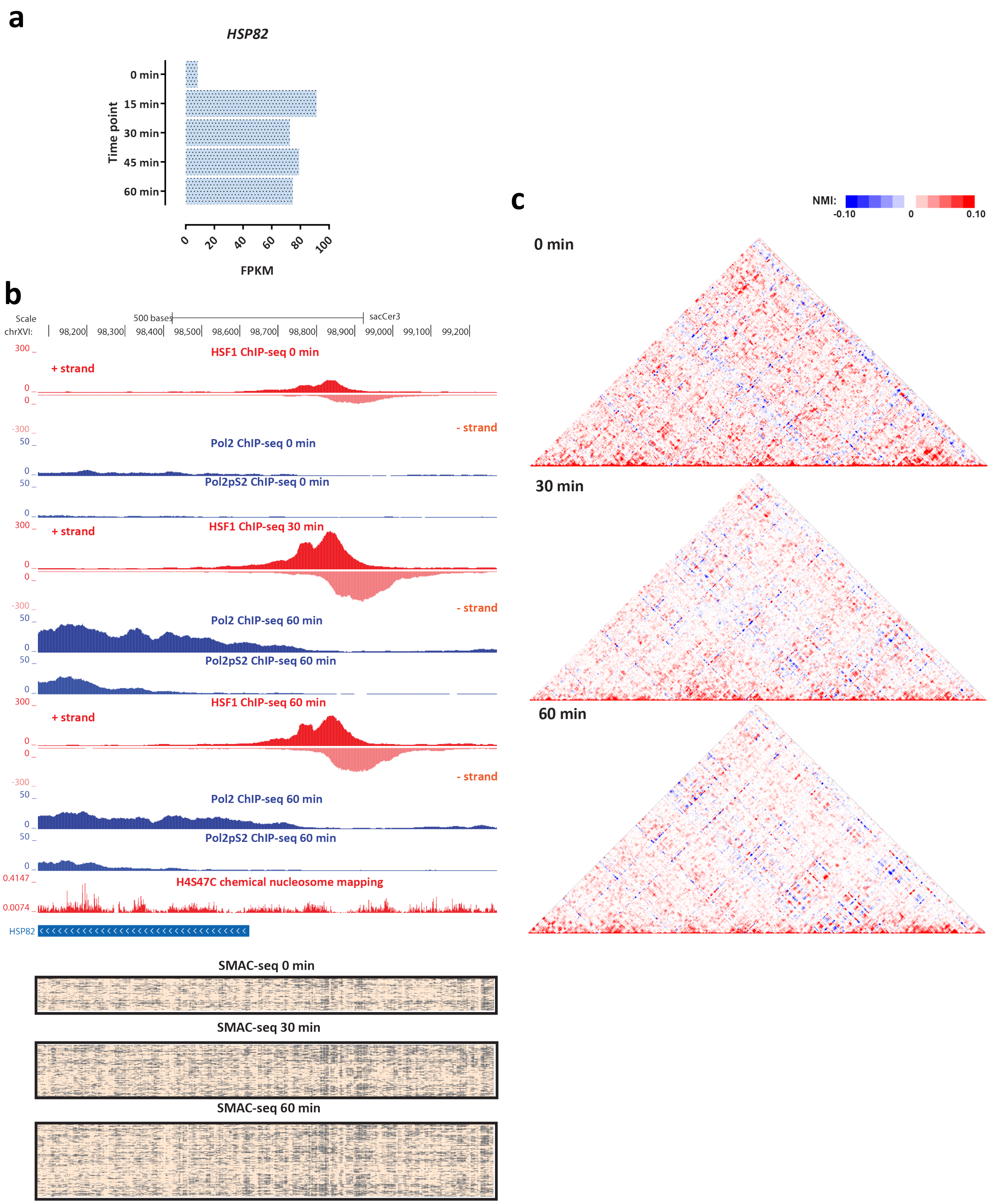


**Supplementary Figure 58: Coordinated changes in chromatin accessibility and nucleosomal occupancy during the yeast stress response**. Shown are changes in RNA Polymerase and HSF1 occupancy (measured by ChIP-seq), SMAC-seq profiles (1-bp resolution, 10-bp aggregate scores) and NMI profiles in the vicinity of the *HSP82* gene.

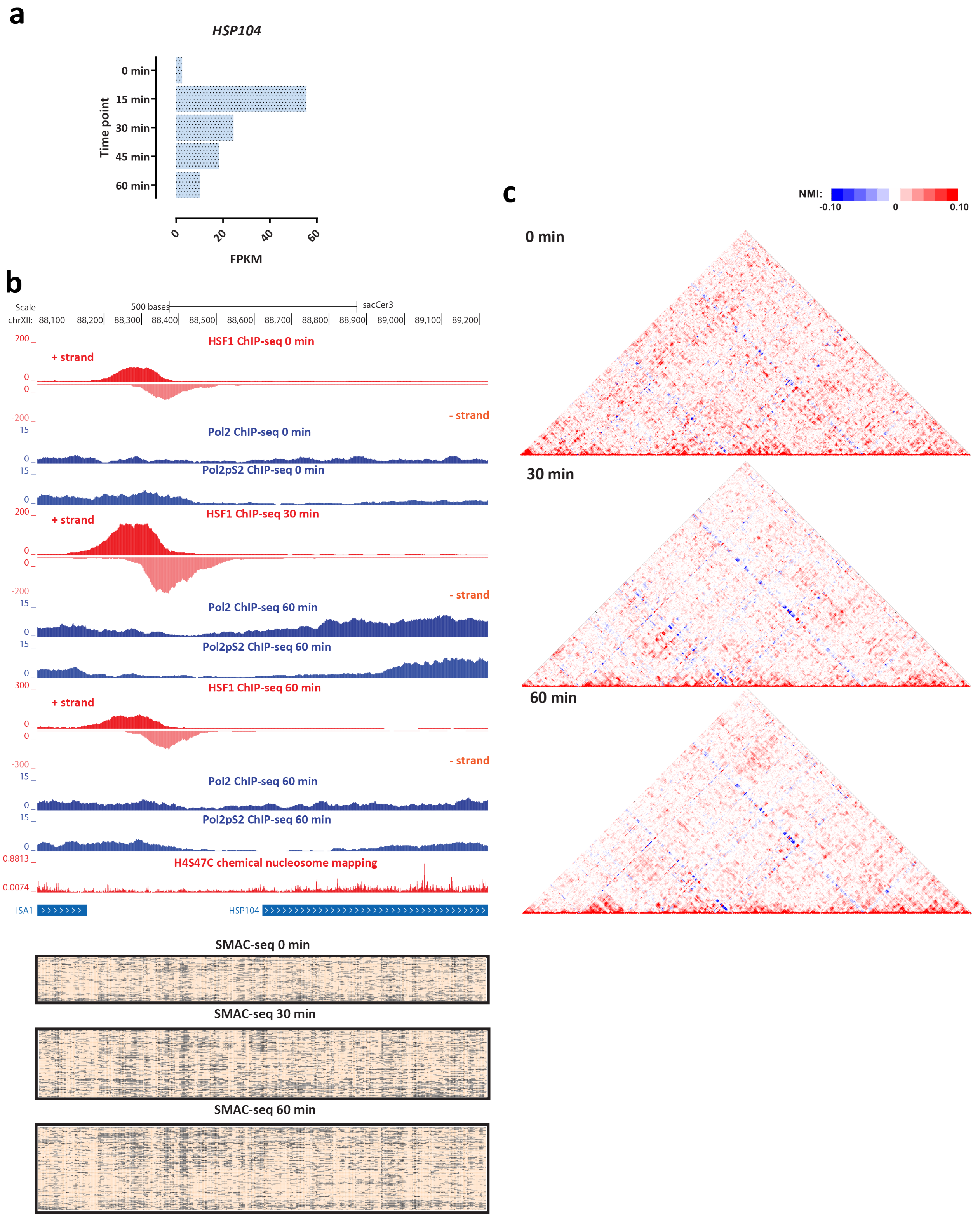


**Supplementary Figure 59: Coordinated changes in chromatin accessibility and nucleosomal occupancy during the yeast stress response**. Shown are changes in RNA Polymerase and HSF1 occupancy (measured by ChIP-seq), SMAC-seq profiles (1-bp resolution, 10-bp aggregate scores) and NMI profiles in the vicinity of the *HSP104* gene.

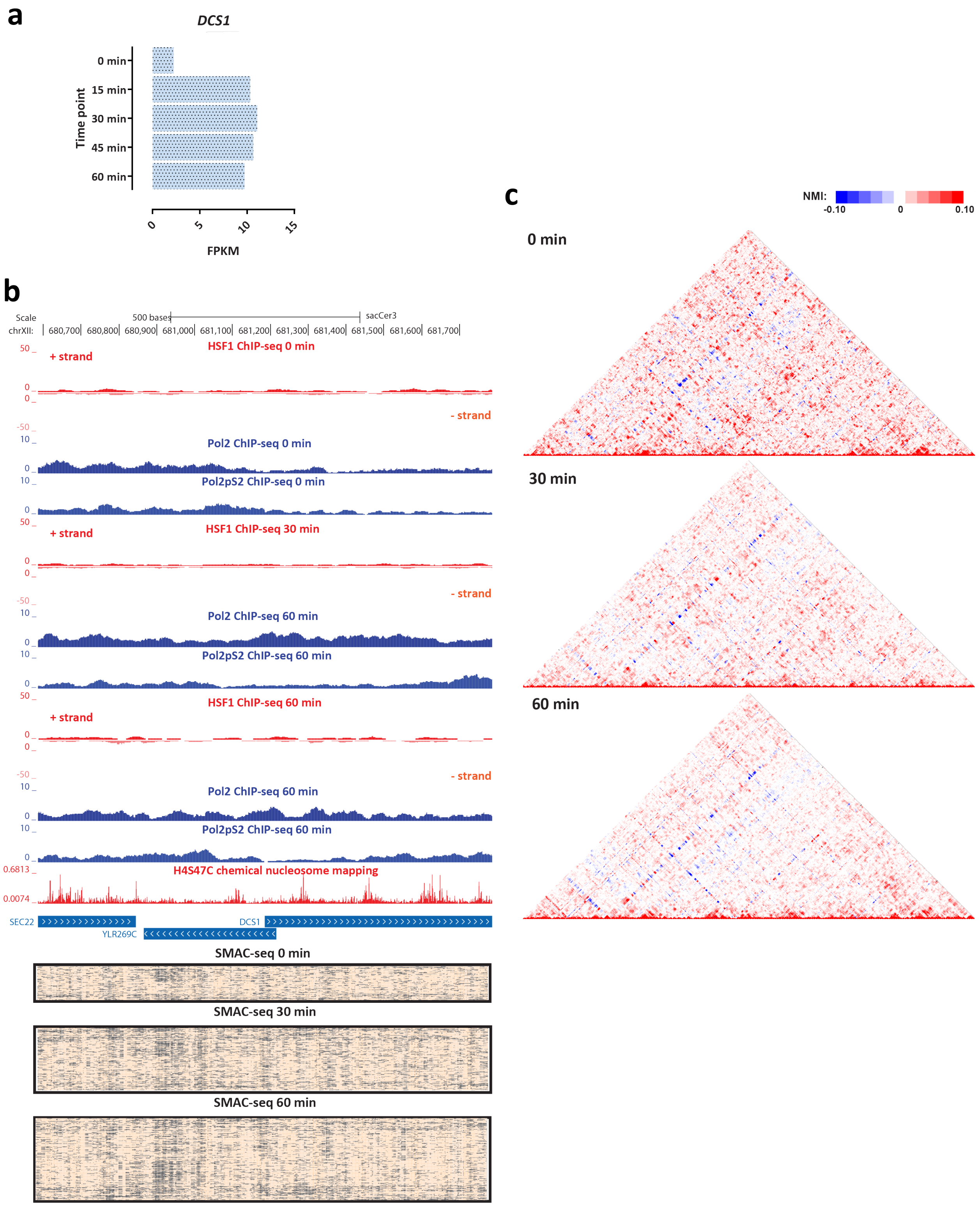


**Supplementary Figure 60: Coordinated changes in chromatin accessibility and nucleosomal occupancy during the yeast stress response**. Shown are changes in RNA Polymerase and HSF1 occupancy (measured by ChIP-seq), SMAC-seq profiles (1-bp resolution, 10-bp aggregate scores) and NMI profiles in the vicinity of the *DSC1* gene.

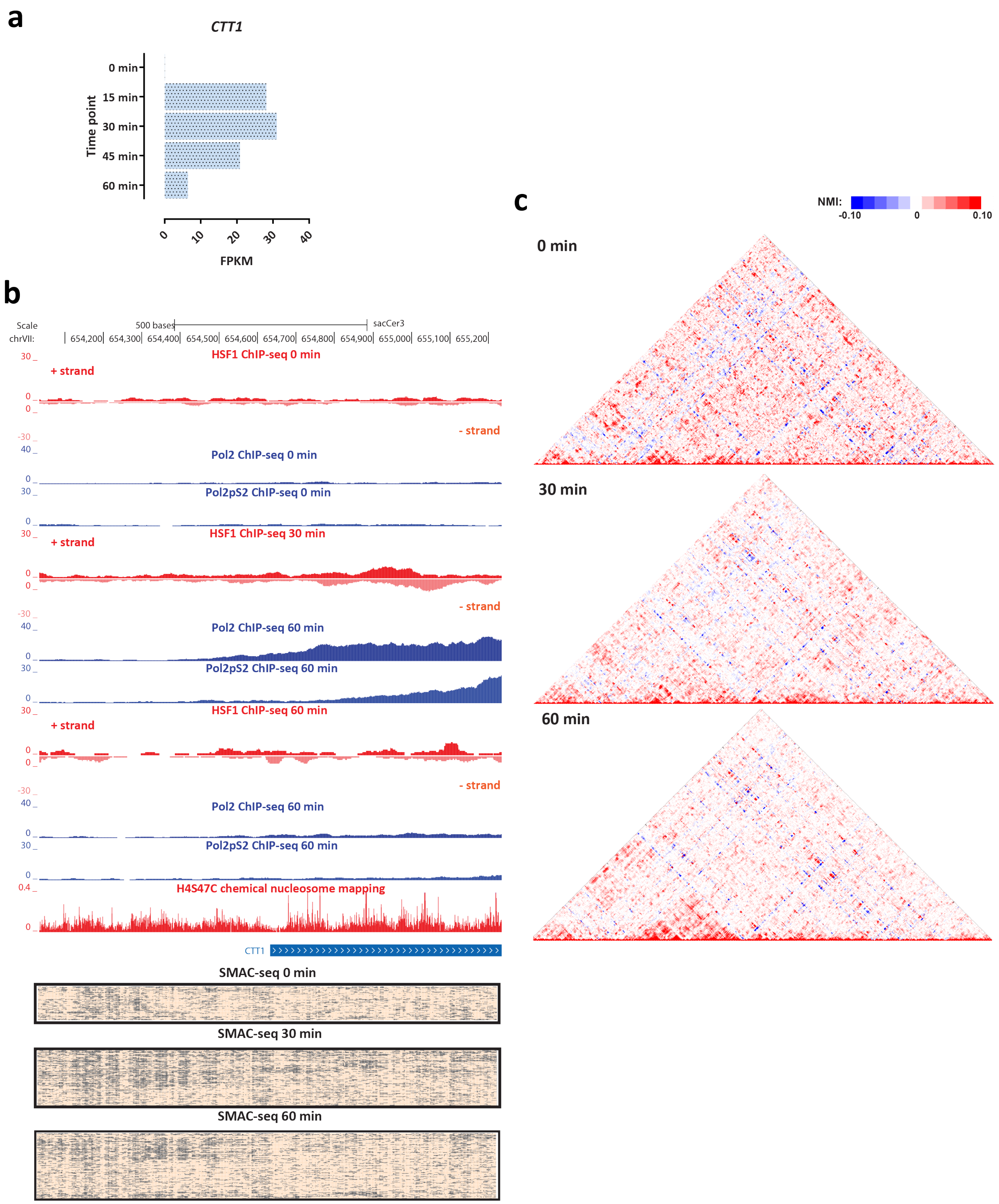


**Supplementary Figure 61: Coordinated changes in chromatin accessibility and nucleosomal occupancy during the yeast stress response**. Shown are changes in RNA Polymerase and HSF1 occupancy (measured by ChIP-seq), SMAC-seq profiles (1-bp resolution, 10-bp aggregate scores) and NMI profiles in the vicinity of the *CTT1* gene.

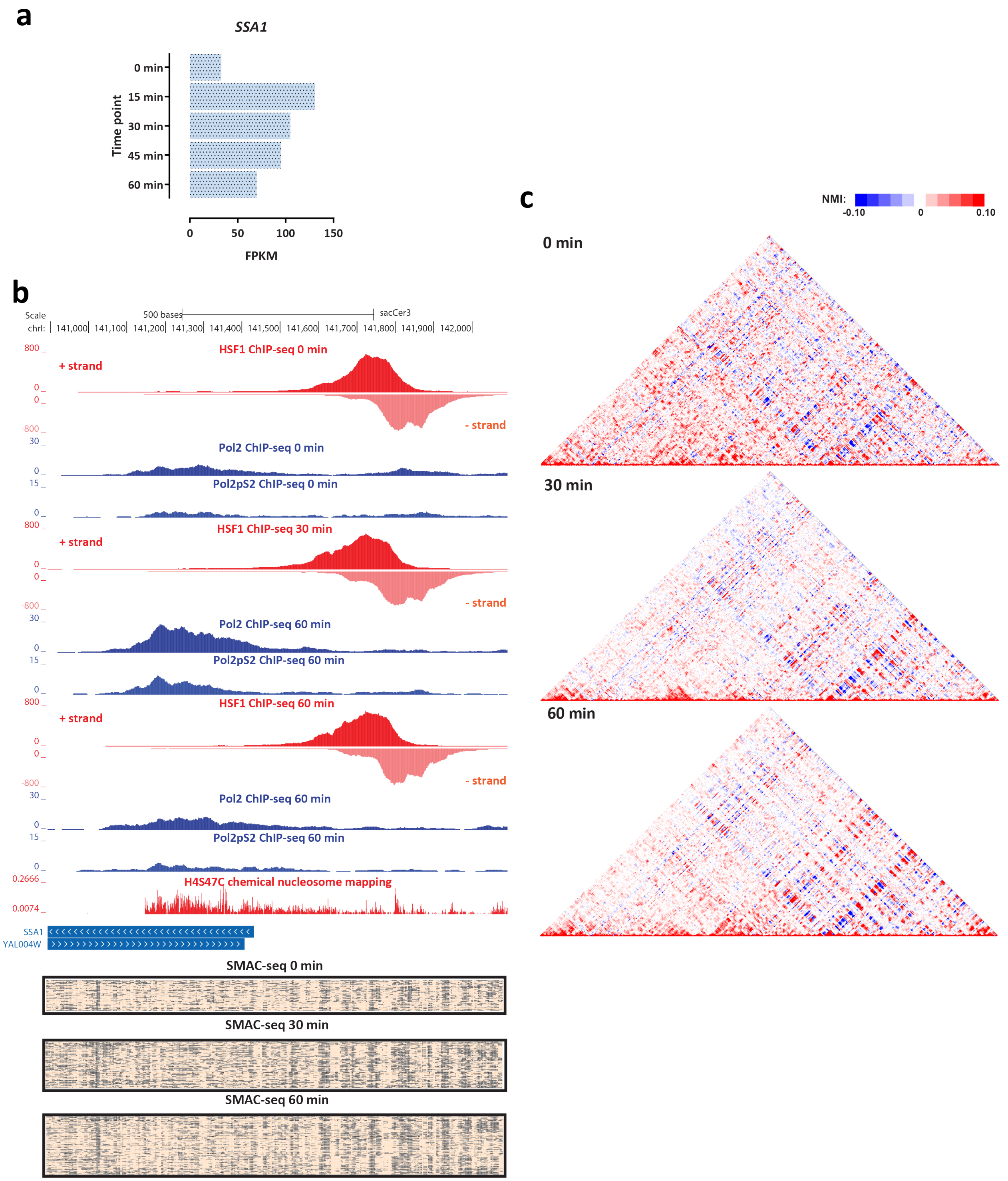


**Supplementary Figure 62: Coordinated changes in chromatin accessibility and nucleosomal occupancy during the yeast stress response**. Shown are changes in RNA Polymerase and HSF1 occupancy (measured by ChIP-seq), SMAC-seq profiles (1-bp resolution, 10-bp aggregate scores) and NMI profiles in the vicinity of the *SSA1* gene.

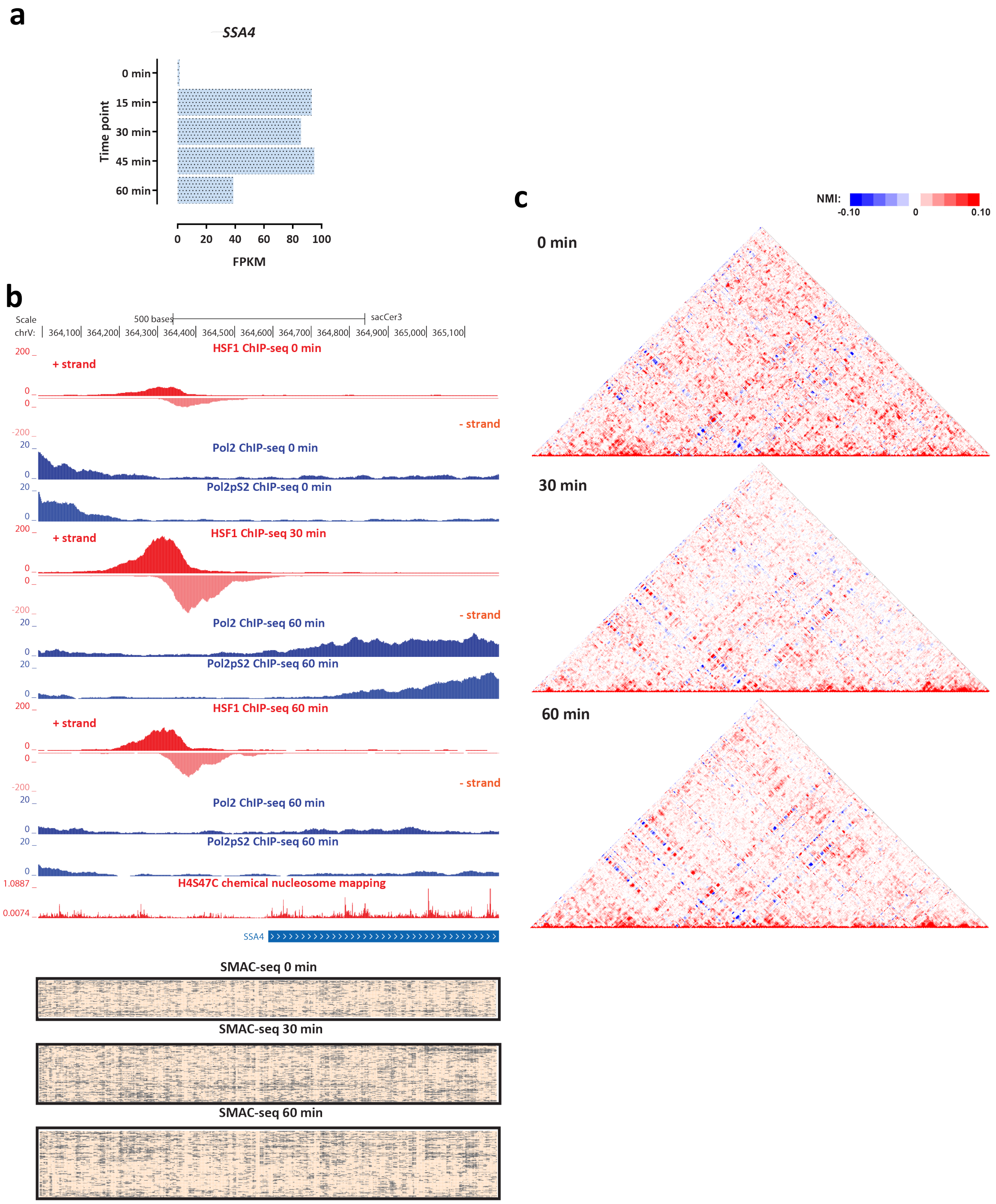


**Supplementary Figure 63: Coordinated changes in chromatin accessibility and nucleosomal occupancy during the yeast stress response**. Shown are changes in RNA Polymerase and HSF1 occupancy (measured by ChIP-seq), SMAC-seq profiles (1-bp resolution, 10-bp aggregate scores) and NMI profiles in the vicinity of the *SSA4* gene.

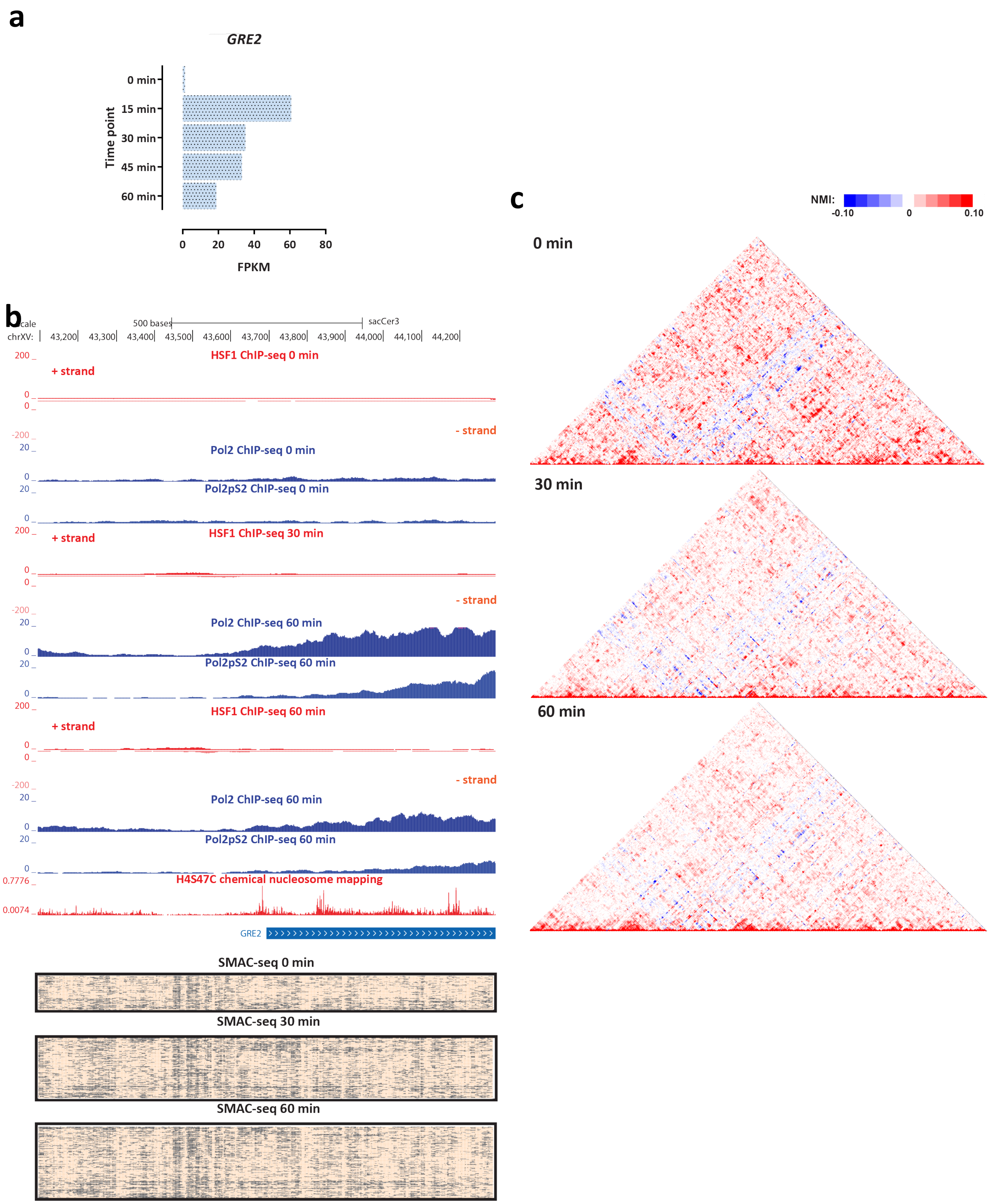


**Supplementary Figure 64: Coordinated changes in chromatin accessibility and nucleosomal occupancy during the yeast stress response**. Shown are changes in RNA Polymerase and HSF1 occupancy (measured by ChIP-seq), SMAC-seq profiles (1-bp resolution, 10-bp aggregate scores) and NMI profiles in the vicinity of the *GRE2* gene.

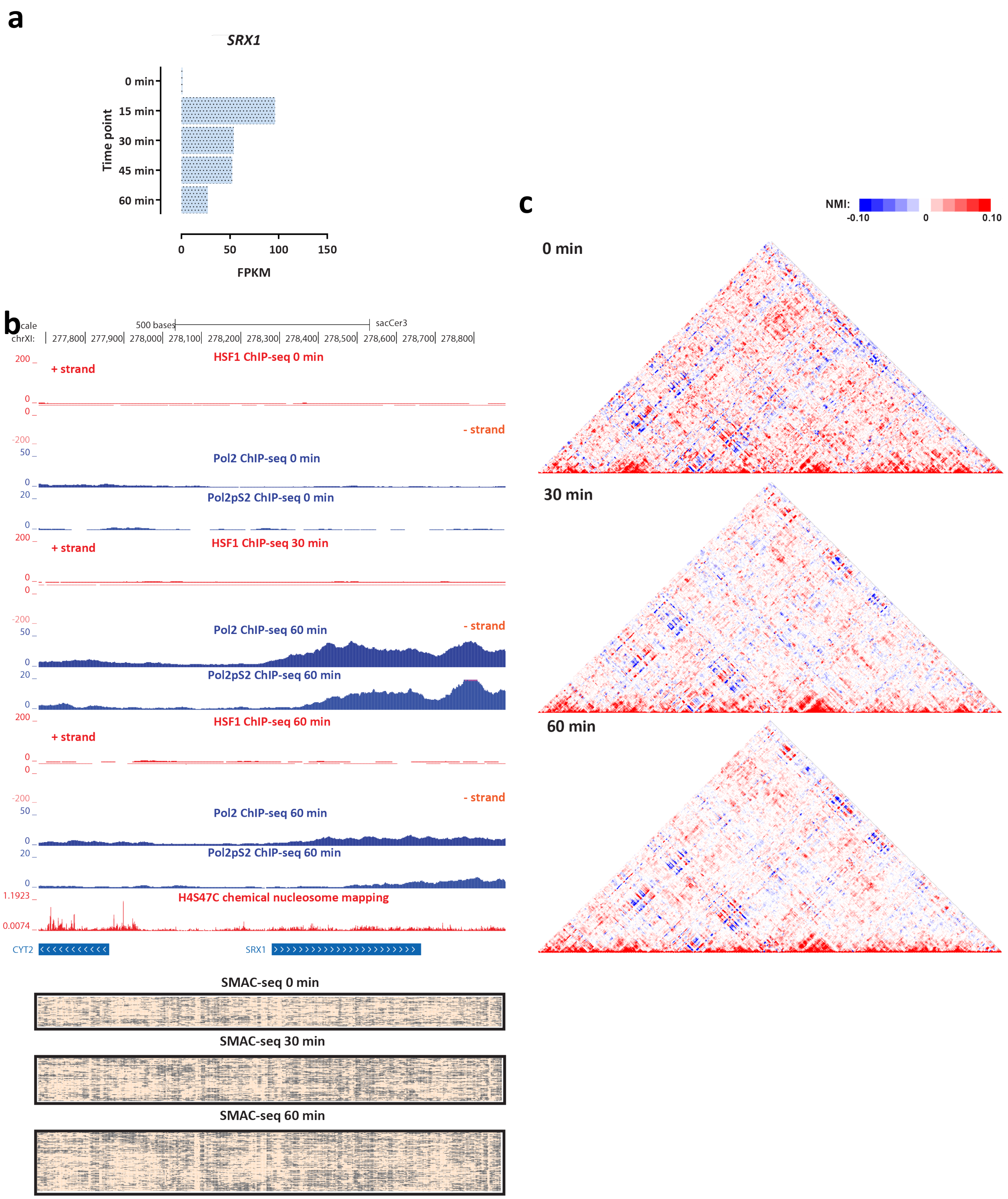


**Supplementary Figure 65: Coordinated changes in chromatin accessibility and nucleosomal occupancy during the yeast stress response**. Shown are changes in RNA Polymerase and HSF1 occupancy (measured by ChIP-seq), SMAC-seq profiles (1-bp resolution, 10-bp aggregate scores) and NMI profiles in the vicinity of the *SRX1* gene.
